# Platelet DKK1 promotes tolerogenic dendritic cells and non-healing responses in cutaneous leishmaniasis

**DOI:** 10.1101/2025.05.27.656395

**Authors:** Olivia C. Ihedioha, Anutr Sivakoses, Haley Q. Marcarian, Malini Sajeev, Diane McMahon-Pratt, Alfred L.M. Bothwell

## Abstract

Dickkopf-1(DKK1) is a classical Wnt antagonist, which is released at the initiation of *Leishmania major* infection through platelet TLR1/2 activation. Using BALB/c mice deficient in platelet MyD88 (MyD88^PKO^) or platelet DKK1 (DKK1^PKO^), we assessed whether the transmission of activation signals through MyD88 and subsequent release of DKK1 are critical in regulating the immune response to *L. major*. At the site of infection, the levels of neutrophil platelet aggregates and activated neutrophils of MyD88^(PKO)^ and DKK1^(PKO)^ mice were reduced. Further, these mice mounted anti-leishmanial Th1-responses and failed to develop progressive lesions. In contrast, WT BALB/c-infected mice developed progressive disease associated with elevated IL-10-producing Th1 and Th2 T cells. Further, elevated CD206^+^ M2 macrophages and tolerogenic DC-10 cells, which favor parasite proliferation, were observed. In vitro, DKK1 promoted DC IL-10 production and blocked TNFα-induction of IL-12. Overall, these results indicate that platelet-DKK1 promotes disease progression through the induction of tolerogenic DCs and subsequent pathological Th2 and IL-10-Th1 T cell-responses.

**Summary statement:** Platelet DKK1 produced in response to TLR1/2 signaling by *Leishmania* parasites drives development of non-healing responses. In vivo/ in vitro analyses indicate that the mechanism underlying this process is the promotion of tolerogenic dendritic cells, driving Th2 and Th1-IL-10 T cell responses.

**Graphical abstract:** 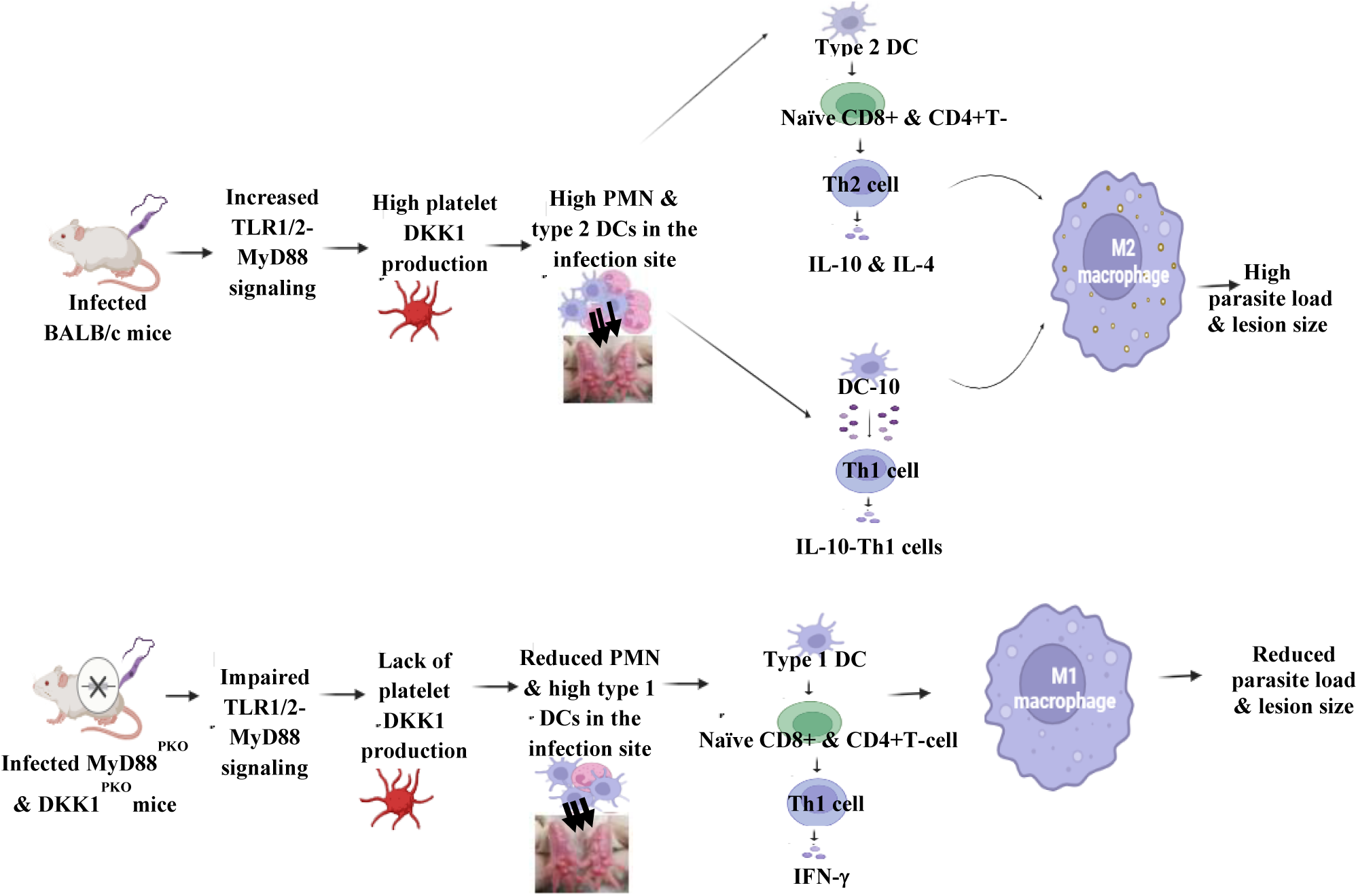

## Introduction

*Leishmania* parasites cause cutaneous leishmaniasis and exhibit a wide range of clinical manifestations, from self-healing lesions to chronic debilitating infections (de Vries and Schallig, 2022). The interplay between infecting agents and host factors determines the control or progression of *Leishmania* infection. An effective immune response against leishmaniasis relies on the combined action of the innate and adaptive immune responses of the infected host (Sacks and Noben-Trauth, 2002). Thus, the CD4^+^ and CD8^+^T cells, as well as cells of the innate immune response such as neutrophils, dendritic cells (DCs) and macrophages, are involved in regulating disease outcome (de Franca et al., 2024, Costa-da-Silva et al., 2022). Conventional dendritic cell type 1(cDC1) and type 2 (cDC2) are critical dendritic cell subsets determining the disease outcome because they preferentially drive Th1 and Th2 responses, respectively (Hilligan and Ronchese, 2020). Production of IL-12 by cDC1 skews the differentiation of naïve T cells to host-protective Th1 cells (Suzue et al., 2008, Ashour et al., 2020). This response promotes classical (M1) macrophage activation by cognate interactions and/or the release of interferon (IFN-γ) and tumor necrosis factor (TNF-α) that are required for the control of infection (Bogdan et al., 2024, Bogdan, 2020). On the other hand, the sustained release of IL-4 and IL-10 in the draining lymph nodes of *L. major*-infected BALB/c mice is decisive for the differentiation and expansion of Th2 cells (Himmelrich et al., 2000, Vacas et al., 2020). Th2 cells, in turn, cause alternative (M2) macrophage activation, resulting in impaired killing of *Leishmania* and disease progression (Alshaweesh et al., 2022, Almeida et al., 2023). Thus, non-healing progressive cutaneous leishmaniasis has been linked to a predominance of Th2 cells (Himmelrich et al., 2000, Vacas et al., 2020), while healing is associated with anti-leishmanial Th1-responses (Scott and Novais, 2016). Additionally, studies have identified CD4^+^IFN-γ+Th1 cells as a source of IL-10 that suppresses the protective immune response against *Leishmania* species (Anderson et al., 2007, Owens et al., 2012, Resende et al., 2013). These IL-10-producing Th1 cells, which are activated early in a strong inflammatory setting, are critical mediators of immune suppression in chronic cutaneous leishmaniasis (Anderson et al., 2007). The detailed characterization of this population indicated that the IL-10- producing Th1 cells are T-bet^+^, Foxp3^−^, and CD25^−^ while maintaining an effector phenotype (Anderson et al., 2007). A subset of tolerogenic dendritic cells (DC-10), endowed with the ability to release IL-10 spontaneously, were capable of polarizing CD4^+^ T cells toward an IL-10- producing Th1 cell phenotype. The tolerogenic DC-10 cells display MHC II^low^, CD80^low^, CD86^low^ and CD40 ^low^ phenotypic markers (Comi et al., 2018). Also, IL-12 production is reported to be absent in monocyte-derived tolerogenic DC-10 cells (Gregori et al., 2010).

It has been established that the disease phenotype in murine leishmaniasis can be established within the first few hours post-infection (Sacks and Noben-Trauth, 2002). This early response is characteristic of the innate immune system, which detects invading pathogens through specific receptors that recognize “pathogen-associated molecular patterns” (Takeda et al., 2003). Pathogen recognition receptors, specifically the family of Toll-like receptors (TLRs), can sense pathogen-associated molecular patterns. TLR engagement initiates innate immune responses and can influence subsequent adaptive immune responses (Pasare and Medzhitov, 2005). A critical role of the myeloid differentiation protein 88 (MyD88), a key downstream protein of TLR signaling, has been demonstrated in the activation of innate immunity (Kaisho and Akira, 2001). Platelets are well-positioned to act as first responders, detecting invading pathogens and triggering the host’s immune response to fight the infection (Portier and Campbell, 2021). Previous studies have confirmed the expression of TLR1-7 and MyD88 in human and mouse platelets (Semple et al., 2011). TLR1, TLR2 and TLR4 are important representatives of these receptors mediating intracellular platelet activation generally via the MyD88-dependent pathway (Niklaus et al., 2022, De Stoppelaar et al., 2015). Activation of platelet TLR elicits a plethora of responses, including the release of inflammatory mediators, modulation of leukocyte inflammatory responses and heterotypic cell aggregation (Hally et al., 2020).

Our previous studies showed that leukocyte-platelet aggregation (LPA) is required for leukocyte migration to the infection site and induction of Th2 immune response during *Leishmania* infection (Ihedioha et al., 2023, Chae et al., 2016). This aberrantly high LPA formation is driven by DKK1 released by activated platelets following recognition of *L. major*-derived lipophosphoglycan (LPG) via TLR1/2 (Ihedioha et al., 2023). Since MyD88 is a downstream adaptor of the TLR1/2 (Niklaus et al., 2022), in the current study, we utilized mice with conditional deletion of MyD88 or DKK1 from platelets to establish that DKK1 released via the Myd88-dependent pathway determines disease outcome. These studies further show that DKK1 critically promotes the shift of dendritic cells towards the DC-10 phenotype, leading to inhibition of dendritic cell maturation and development of the chronic Th2 immune response critical for disease progression in *L. major* infection. Together with the regulation of PMN/LPA aggregation which selectively draws host cells to the site of infection, DKK1 controls the developing milieu promoting susceptibility to infection.

## Results

### Minimal P-selectin expression in activated platelets from infected MyD88^(PKO)^ mice on days 3 and 14 PI

P-selectin is a thrombo-inflammatory molecule and a well-established marker for measuring platelet activation (Fabricius et al., 2021). *L. major-*derived LPG is reported to interact with the immune cell-expressed TLR2 and modulate anti-leishmanial immune responses (Becker et al., 2003, Aebischer et al., 2005, Halliday et al., 2016, Ghosh et al., 2023). We have previously shown that *L. major-*derived LPG is the key virulence factor that activates platelet TLR1/2 to induce DKK1 production (Ihedioha et al., 2023). Since TLR1/2 utilizes MyD88 to initiate its signaling cascade, this study investigated whether LPG and MyD88 influence platelet activation by regulating platelet TLR2 signaling. Expression of P-selectin was determined in platelets obtained from infected BALB/c, DKK1^(PKO)^, MyD88^(PKO)^ and non-infected BALB/c mice for varying periods of time. Relative to the infected BALB/c and DKK1^(PKO)^ mice, P-selectin expression was significantly suppressed in platelets obtained from infected MyD88^(PKO)^ mice on days 3 and 14 PI (Fig.1B & C). In addition, the level of P-selectin expression was reduced on day 14 PI (compared to day 3 PI) in all experimentally infected mice (Fig.1B & C). As the MyD88 adaptor protein is intact and functional in DKK1^(PKO)^ infected mice, these data confirmed that transmission of activation signals through the MyD88 signaling pathway is crucial in platelet activation in the early stages of *L. major* infection.

**Figure 1:**
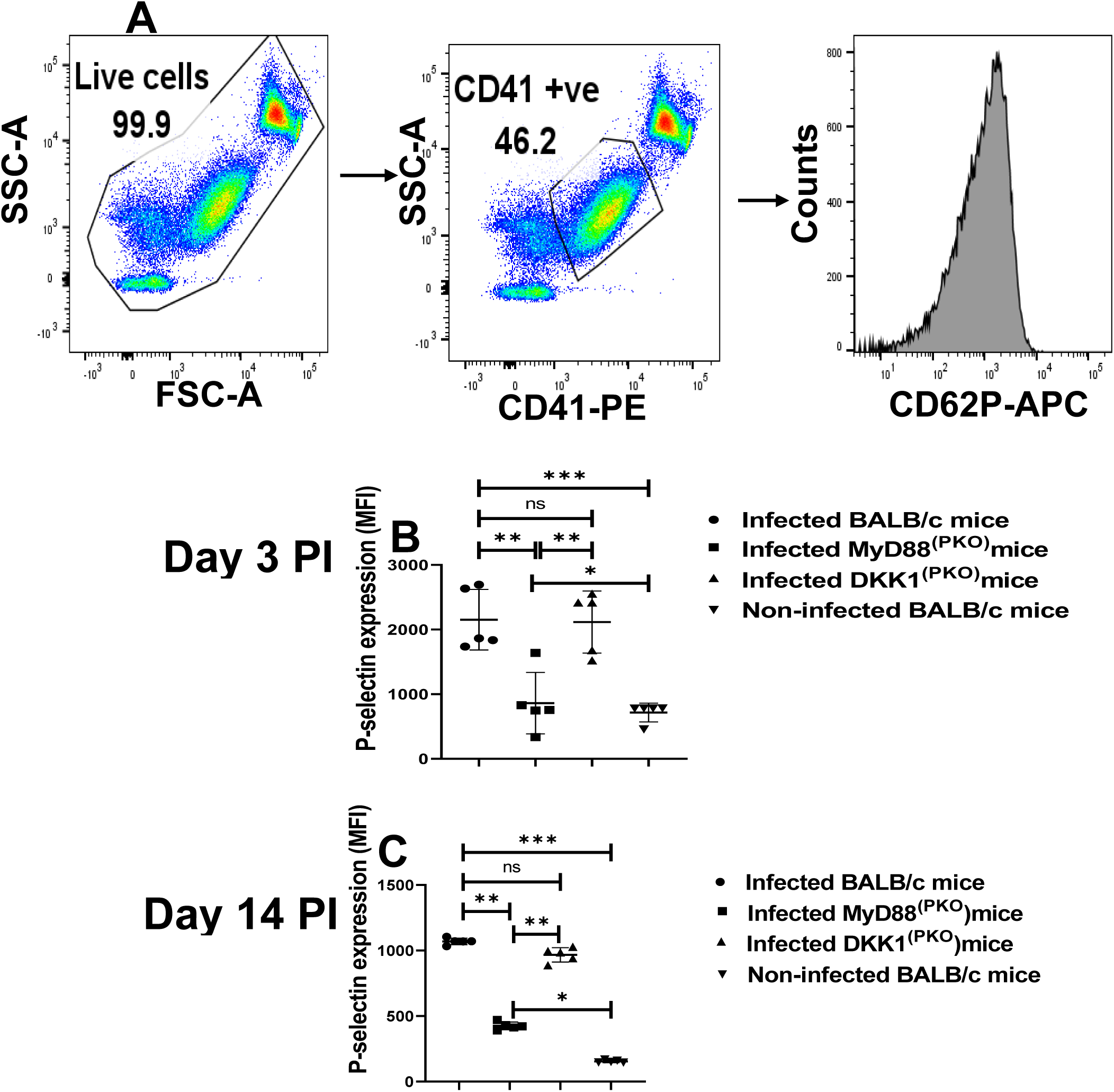
Minimal P-selectin expression in activated platelets from infected MyD88^(PKO)^mice. BALB/c, MyD88^(PKO)^, and DKK1^(PKO)^ mice were challenged with infective metacyclic promastigote (2 x 10^6^ parasites, n = 5) of WT *L. major* strain via the footpad. Control mice (n = 5) were given 0.9% NaCl saline. Blood was collected via the maxillary vein on days 3 and 14 PI. Isolated platelet samples were analyzed by flow cytometry for P-selectin expression. Representative dot plots and histogram indicate the analysis of P-selectin expression on day 3 PI **(A)**. The MFI obtained on days 3 and 14 PI from each experimental group indicates the expression of P-selectin by CD41^+^ cells **(B) & (C).** In all the experiments, WT-infected and non-infected mice served as positive and negative controls, respectively. Results are presented as mean (± SEM) and are representative of two independent experiments. One-way ANOVA with Bonferroni’s post hoc test was performed to analyze the data *p < 0.05, **p < 0.01, ***p < 0.001; ‘ns’ indicates not significant (p > 0.05).

### Decreased DKK1 production in infected MyD88^(PKO)^ and DKK1^(PKO)^ mice on day 3 PI

MyD88 is the key downstream signaling protein of TLR1/2 (Niklaus et al., 2022), and DKK1 production during *Leishmania* infection is primarily dependent on TLR1/2 (Ihedioha et al., 2023). Thus, to assess whether the deficiency of MyD88 impairs DKK1 production, we compared the plasma DKK1 levels in infected BALB/c, DKK1^(PKO)^, MyD88^(PKO)^ and non-infected BALB/c mice on day 3 PI. We confirmed significantly decreased production of plasma DKK1 in DKK1^(PKO)^ and MyD88^(PKO)^ infected mice compared to infected BALB/c mice (Fig. 2). This finding suggests that MyD88 activation signal is critical for DKK1 production from activated platelets in infected BALB/c mice.

**Figure 2:**
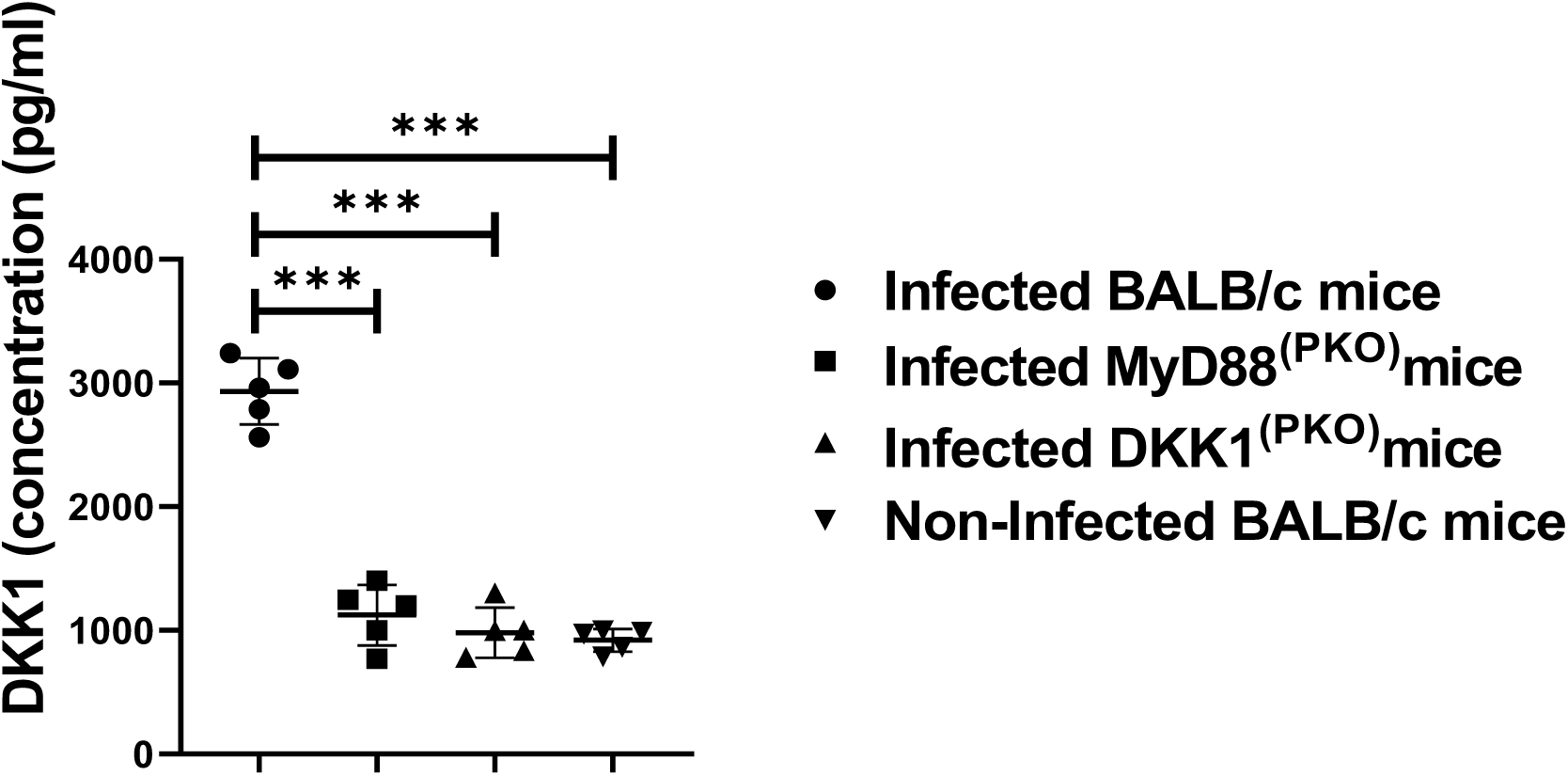
Decreased DKK1 production in infected MyD88^(PKO)^ and DKK1^(PKO)^ mice. BALB/c, MyD88^(PKO)^, and DKK1^(PKO)^ mice were challenged with infective metacyclic promastigote (2 x 10^6^ parasites, n = 5) of WT *L. major* strain via the footpad. Control mice (n = 5) were given 0.9% NaCl saline. Blood was collected via the maxillary vein on day 3 PI. Plasma samples were analyzed by ELISA for DKK1 production. In all experiments, Infected and non- infected BALB/c mice served as positive and negative controls, respectively. Results are presented as mean +/- SEM and are representative of three replicate experiments. One-way ANOVA with Bonferroni’s post hoc test was performed to analyze the data ***p < 0.001. (p > 0.05).

### Impaired LPA and NPA formation in infected MyD88^(PKO)^ and DKK1^(PKO)^ mice on days 3 and 14 PI

P-selectin (CD62P) and PSGL-1 mediate leukocyte platelet aggregation formed by adhesion between activated platelets and mature leukocytes under inflammatory conditions (Irving et al., 2004, Chae et al., 2016, Finsterbusch., 2018). Previously, we showed that LPG-activated platelets promote P-selectin expression, resulting in platelet leukocyte aggregation (LPA), which is essential for an early migration of leukocytes to the inflammatory site (Zuchtriegel et al., 2016, Ihedioha et al., 2023, Ihedioha et al., 2024). Furthermore, pre-treatment of mice with DKK1 inhibitor prior to infection with *L. major* reduced the elevation of leukocyte platelet aggregate formation found in blood of infected mice (Chae et al., 2016). These findings suggest that leukocyte platelet aggregation and leukocyte infiltration at the inflammatory site are driven by DKK1 production. Given that LPG activates the TLR1/2-MyD88 pathway to induce DKK1 production, we consider the possibility that the deletion of MyD88 protein or failure of DKK1 production might reduce LPA formation in blood obtained from DKK1^(PKO)^ and MyD88^(PKO)^ infected mice. Relative to the BALB/c-infected mice, LPA was significantly impaired in DKK1^(PKO)^ and MyD88^(PKO)^ infected mice on day 3 PI (Fig. 3A & B).

**Figure 3:**
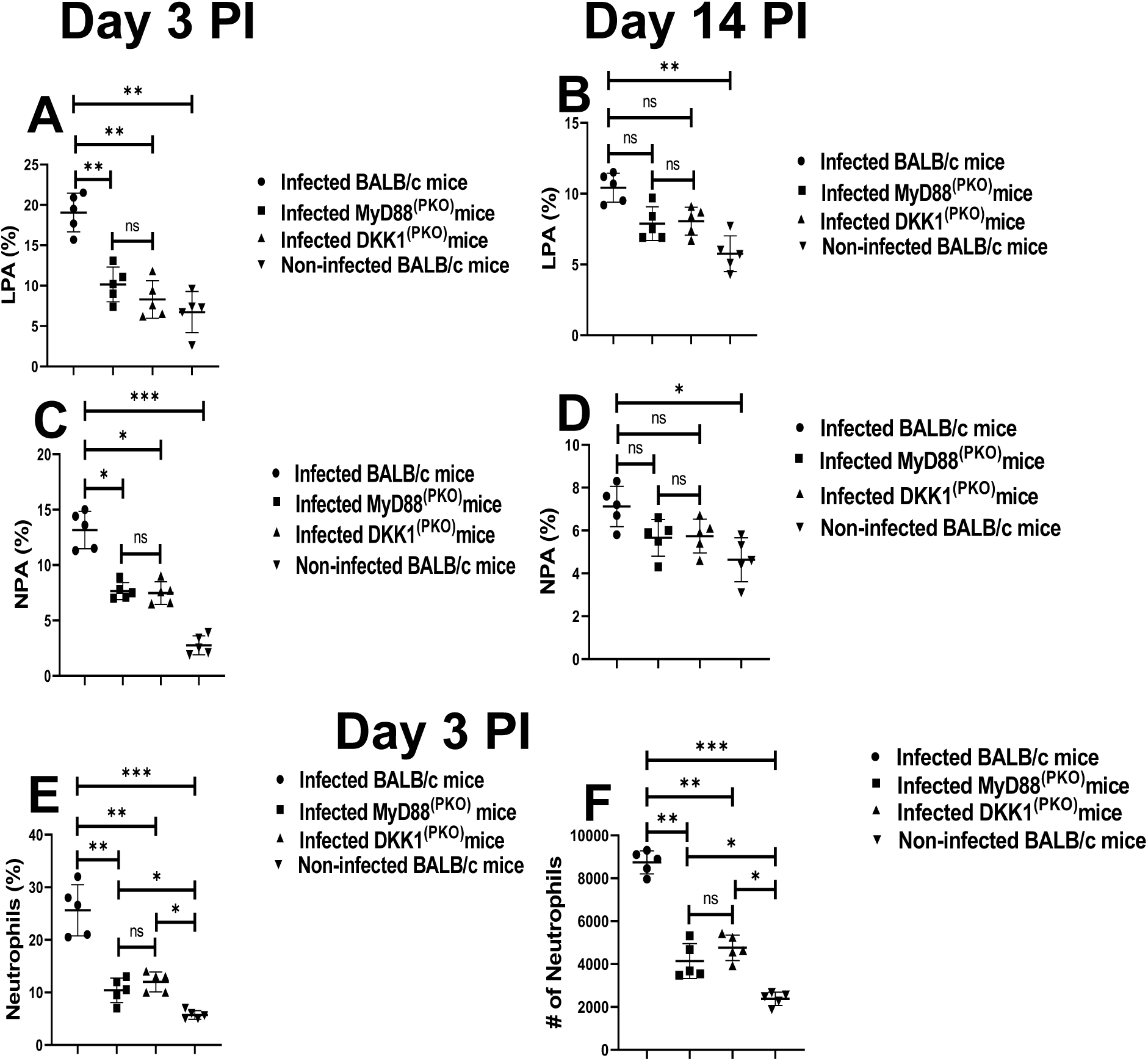
Reduced neutrophils, LPA and NPA formation in infected MyD88^(PKO)^ and DKK1^(PKO)^ mice. BALB/c, MyD88^(PKO)^, and DKK1^(PKO)^ mice were challenged with infective metacyclic promastigote (2 x 10^6^ parasites, n = 5) of WT *L. major* strain via the footpad. Control mice for LPA detection (n = 5) and NPA detection (n = 10/2 feet per mouse) were given 0.9% NaCl saline. Blood was collected via the maxillary vein on days 3 and 14 PI for the determination of LPA formation. Cells from the infected footpad were collected on days 3 and 14 post-infection (PI) to assess NPA aggregation and neutrophil infiltration. Samples were analyzed by flow cytometry for LPA, NPA and neutrophils. Representative flow cytometry dot plots showing the analysis of LPA and NPA performed on day 3 PI and **t**he dot plots from each sample in all the experimental groups are indicated in **Fig. S1**. The column graphs **(A) & (B)** indicate the percentage of LPA molecules by CD45^+^ cells, while the percentage of NPA formed by Ly6G^+^ and CD41^+^ cells **(A)** and infiltrated neutrophils **(B &C**) is indicated. In all the experiments, BALB/c infected and non- infected mice served as positive and negative controls, respectively. Results are presented as mean (± SEM) and are representative of two independent experiments. One-way ANOVA with Bonferroni’s post hoc test was performed to analyze the data *p < 0.05, **p < 0.01; ***p < 0.001; ‘ns’ indicates not significant (p > 0.05).

Further, we characterize the levels of neutrophil platelet aggregates (NPA) within infected tissue using infected BALB/c, DKK1^(PKO)^, MyD88^(PKO)^ and non-infected mice. NPA are required for the migration of activated neutrophils to the infection site. Consistent with the decreased circulating LPA in DKK1^(PKO)^ and MyD88^(PKO)^ infected mice, data showed reduced NPA at the infection site of DKK1^(PKO)^ and MyD88^(PKO)^ infected mice on day 3 PI compared to BALB/c-infected mice (Fig. 3C & D). On day 14 PI, the formation of LPA and NPA was reduced in all experimentally infected mice compared to day 3 PI (Fig. 3A, B, C, & D). On day 14 PI the LPA and NPA formation were comparable in all the infected groups of mice (Fig. 3B & D). Further, the percentage and number of neutrophils at the infection site of DKK1^(PKO)^ and MyD88^(PKO)^ infected mice were significantly reduced on day 3 PI (Fig. 3E & F). These data suggest that DKK1 produced via the TLR1/2- MyD88 pathway might be critical to the elevated LPA and NPA formation in infected BALB/c mice and the transit of these cells to the site of infection.

### Decreased MPO^+^, CD11b^+^ and MHC class II^+^ neutrophils in MyD88^(PKO)^ and DKK1^(PKO)^- infected mice on days 3 and 14 PI

Myeloperoxidase, a heme enzyme abundantly expressed by PMN, has been widely used as a marker of neutrophil activation (Arnhold, 2020). We have previously shown that DKK1 signaling via PMN-LRP6 increased neutrophil MPO expression in the infection site of *L. major* infected mice (Ihedioha et al., 2024). To determine whether failure to transmit activation signal due to deficiency of MyD88 or impaired DKK1 production regulates infiltration of activated neutrophils to the infection site, MPO^+^ neutrophils were assessed in cells obtained from the footpad of infected BALB/c, DKK1^(PKO)^, MyD88^(PKO)^ and non-infected mice. In comparison to the infected BALB/c mice, MPO^+^ neutrophils were significantly reduced in DKK1^(PKO)^ and MyD88^(PKO)^ infected mice (Fig. 4A & B).

**Figure 4:**
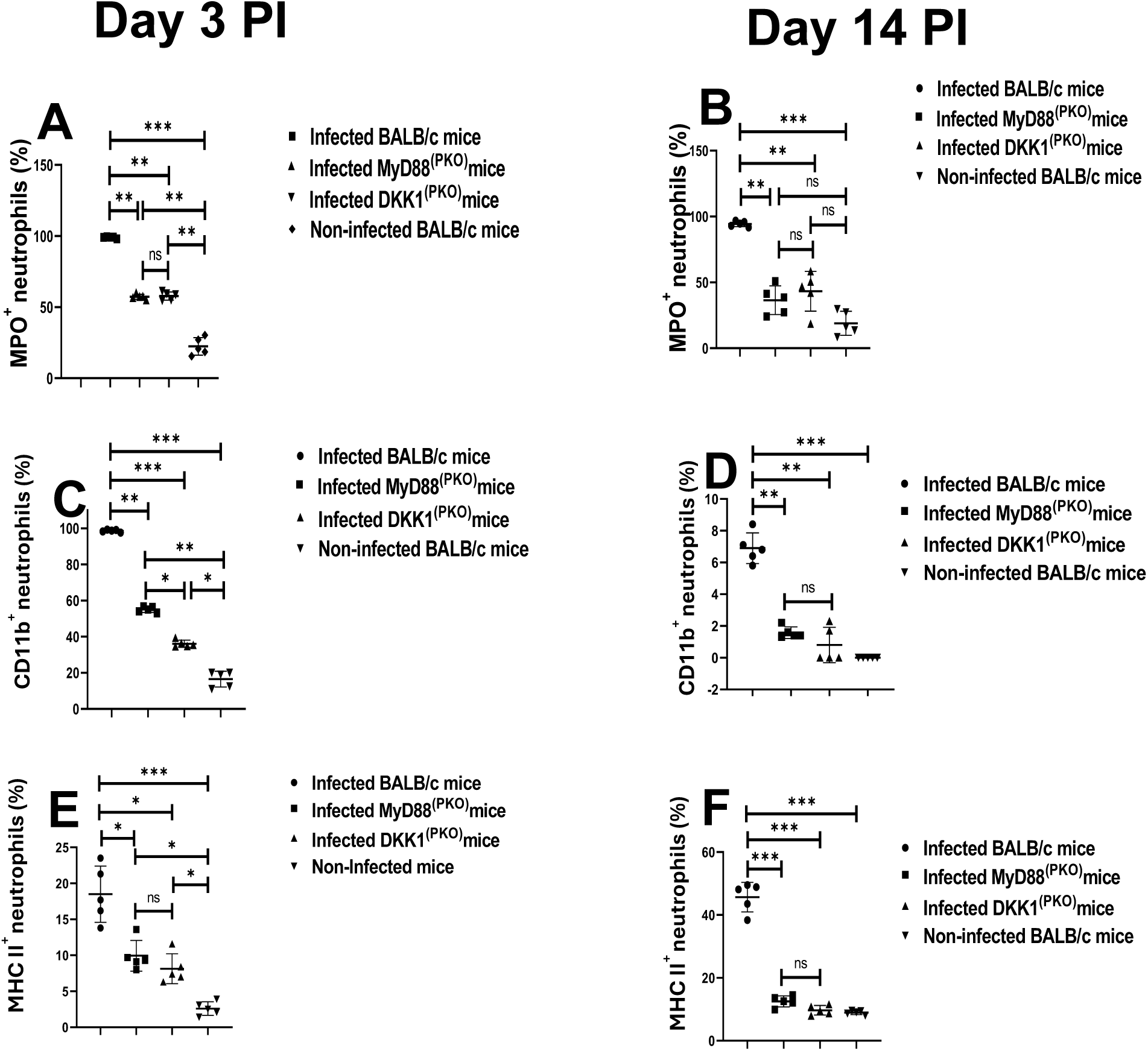
Decreased MPO^+^, CD11b^+^ and MHC class II^+^ neutrophils in MyD88^(PKO)^ and DKK1^(PKO)^ -infected mice. The WT-BALB/c, MyD88^(PKO)^ and DKK1^(PKO)^ mice were challenged with infective metacyclic promastigote (2 x 10^6^ parasites, n = 5) of *L. major* via the footpad. Non-infected BALB/c mice (n = 10/2 feet per mouse) were given 0.9% NaCl saline. Cells were isolated from the footpads of all infected and non-infected mice at days 3 and 14 PI. Samples were analyzed by flow cytometry for MPO^+^, CD11b^+^ and MHC class II^+^ neutrophils. A representative flow cytometry dot plot showing the analysis of MPO^+^, CD11b^+^ and MHC class II**^+^** neutrophils on day 3 PI and the dot plot of each sample in all the experimental groups is presented in **Figure S2.** The percentage of MPO^+^, CD11b^+^ and MHC class II^+^ cells in the different experimental groups is shown in column graphs **(A, B, C, D & E)**. In all experiments, infected and non-infected BALB/c mice served as positive and negative controls, respectively. Results are presented as mean (± SEM) and are representative of two independent experiments. One-way ANOVA with Bonferroni’s post hoc test was performed to analyze the data *p < 0.05; **p < 0.01; ***p < 0.001; ns-non-significant (p > 0.05).

Activated neutrophils are also detected using upregulated CD11b and MHC class II expression in neutrophils (Gosselin et al., 1993, Sandilands et al., 2005). To further assess whether MyD88 activation signals and subsequent DKK1 production are crucial in the migration of activated neutrophils to the infection site, CD11b^+^ and MHC class II^+^ neutrophils obtained from the footpads of infected BALB/c, DKK1^(PKO)^, MyD88 ^(PKO)^ and non-infected BALB/c mice were assessed. In comparison to the infected BALB/c mice, we observed a significant decrease in CD11b^+^ and MHC II^+^ neutrophils in DKK1^(PKO)^ and MyD88^(PKO)^ infected mice (Fig. 4C, D, E, & F). In addition, MPO^+^, CD11b^+^ and MHC II^+^ neutrophils were reduced on day 14 PI in all experimentally infected mice compared to day 3 PI (Fig. 4A, B, C, D, E, & F). Since fewer MPO^+^, CD11b^+^, and MHC class II^+^ neutrophils were found in DKK1^(PKO)^ and MyD88^(PKO)^ infected mice, these data suggest that neutrophil persistent activation and migration to the infection site in BALB/c mice is promoted by DKK1 release via the MyD88 signaling pathway.

### Increased CD38^+^CD206^-^ macrophages and CD8α^+^CD11b^-^ dendritic cells in infected DKK1^(PKO)^ and MyD88 ^(PKO)^ mice

Previously, we demonstrated that treatment with a DKK1 inhibitor diminished leukocyte accumulation at the inflammatory site following parasite infection (Chae et al., 2016). Although the role of PMN recruitment and LPA formation clearly depends on DKK1, we considered the possibility that the deletion of MyD88 or the absence of DKK1 in platelets might also impact the infiltration of macrophages and dendritic cells to the infection site. The migration of dendritic cells and macrophages to the infection site increased by day 14 PI compared to day 7 PI (Fig. 5A, B, C, & D). However, the infiltration of macrophages and dendritic cells to the infection site in DKK1^(PKO)^ and MyD88^(PKO)^ mice was significantly impaired on days 7 and 14 PI compared to infected BALB/c mice (Fig. 5A, B, C, & D). It is of interest that although LPA and PMN infiltration becomes comparable in all groups of mice by day 14, dendritic cells and macrophage remained at lower levels in DKK1^(PKO)^ and MyD88^(PKO)^ mice than in WT BALB/c mice.

**Figure 5:**
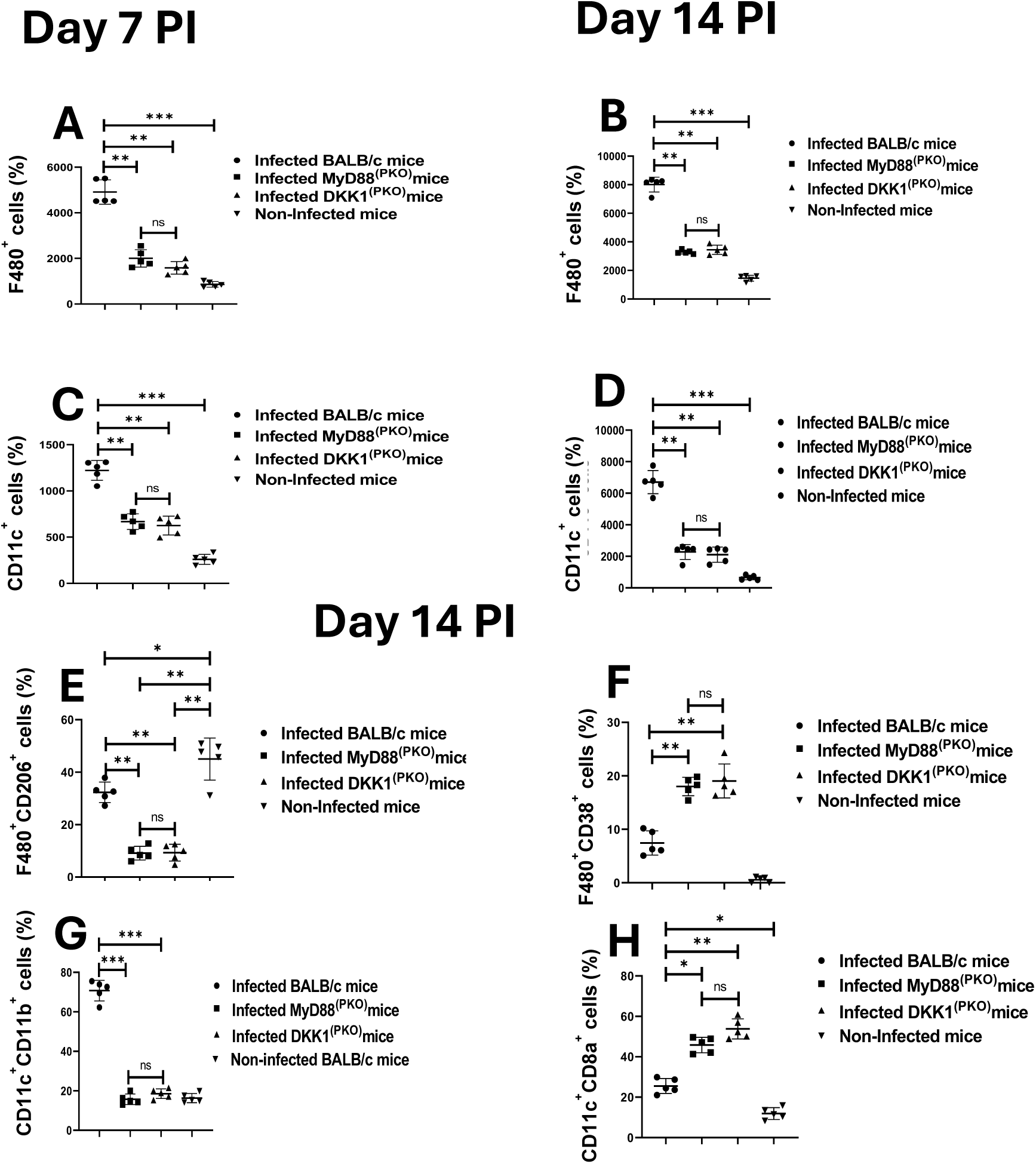
Increased CD38^+^ macrophages and CD8α^+^ dendritic cells in MyD88^(PKO)^ and DKK1^(PKO)^ -infected mice. One week and two weeks post-infection, cells from the infected foot of each mouse in BALB/c, MyD88^(PKO)^ ^and^ DKK1^(PKO)^ were harvested and counted for macrophages (**A & B**) and dendritic cells (**C &D**) by flow cytometry. Two weeks post-infection, the infected foot from each mouse in BALB/c, MyD88^(PKO)^ and DKK1^(PKO)^ were examined for CD206^+^ and CD38^+^ macrophages (**E & F**), as well as CD11b^+^ and CD8α^+^ dendritic cells (**G & H**) by flow cytometry. Representative flow cytometry dot plots showing the analysis of CD206^+^, CD38^+^ macrophages and CD11b^+^, CD8α^+^ dendritic cells performed on day 14 PI and a dot plot of each sample in all the experimental groups are presented in **Figure S3.** Results are presented as mean +/- SEM and are representative of two independent experiments. Data analysis was done using one-way ANOVA with Bonferroni’s post hoc test *p < 0.05, **p < 0.01, ***p < 0.001, and “ns” indicates non-significant.

Having observed a significant decrease in the frequency of macrophages and dendritic cells in infected DKK1^(PKO)^ and MyD88^(PKO)^ mice, we investigated whether DKK1 influences the M1-M2 balance, thereby shaping the disease outcome. Accordingly, we compared the abundance of M1 (F4/80^+^CD38^+^CD206^-^) and M2 (F4/80^+^CD38^-^CD206^+^) subsets in dermal macrophages obtained from infected BALB/c, DKK1^(PKO)^, MyD88^(PKO)^ and non-infected BALB/c mice. Relative to infected BALB/c mice, DKK1^(PKO)^ and MyD88^(PKO)^ infected mice showed a reduced F4/80^+^CD38^-^ CD206^+^ cell population and an increased F4/80^+^CD38^+^CD206^-^ cell population. Notably, the F4/80^+^CD38^-^CD206^+^ cell population dominates in the infection site in BALB/c mice (Fig. 5E & F).

The cDC1 and cDC2 subsets of conventional dendritic cells at the infection site were distinguished based on the surface expression of CD11c, CD11b and CD8α molecules, as previously reported (Szulc-Dąbrowska et al., 2023). In general, the percentage of cDC1 (CD11c^+^CD11b^−^CD8α^+^) and cDC2 (CD11c^+^CD11b^+^CD8α^-^) cells were compared in the experimental and control mice. We observed a significant decrease in CD11c^+^CD11b^+^ cells in infected DKK1^(PKO)^ and MyD88^(PKO)^ mice compared to infected BALB/c mice. However, the portion of CD11c^+^CD8α^+^ cells was significantly higher in infected DKK1^(PKO)^ and MyD88^(PKO)^ mice (Fig. 5G & H). This data suggests that DKK1 released via the MyD88-dependent pathway promotes the infiltration of M2 macrophages and cDC2 cells at the infection site in BALB/c mice.

### Impaired Th2 cytokine production and upregulated IFN-γ in MyD88^(PKO)^ and DKK1^(PKO)^ infected mice on weeks 2 and 15 PI

Previously, we have shown using in vitro studies that DKK1 preferentially induces Th2 cytokines (Chae et al., 2016). To confirm whether failure to transmit activation signals due to a deficiency of MyD88 or a defect in DKK1 production promotes a Th1 response, cytokine production from SLAG-activated lymph node cells obtained from infected BALB/c, DKK1^(PKO)^, MyD88^(PKO)^, and non-infected BALB/c mice was assessed. Compared to infected BALB/c, the production of IL-10 and IL-4 was significantly inhibited in infected DKK1^(PKO)^ and MyD88^(PKO)^ mice on weeks 2 and 15 PI. However, IFN-γ production was elevated in the mutant mice (Fig. 6A, B, C, D, E & F). These data confirmed that the Th2 response in BALB/c infected mice is dependent on DKK1 produced via the TLR1/2-MyD88-dependent pathway.

**Figure 6:**
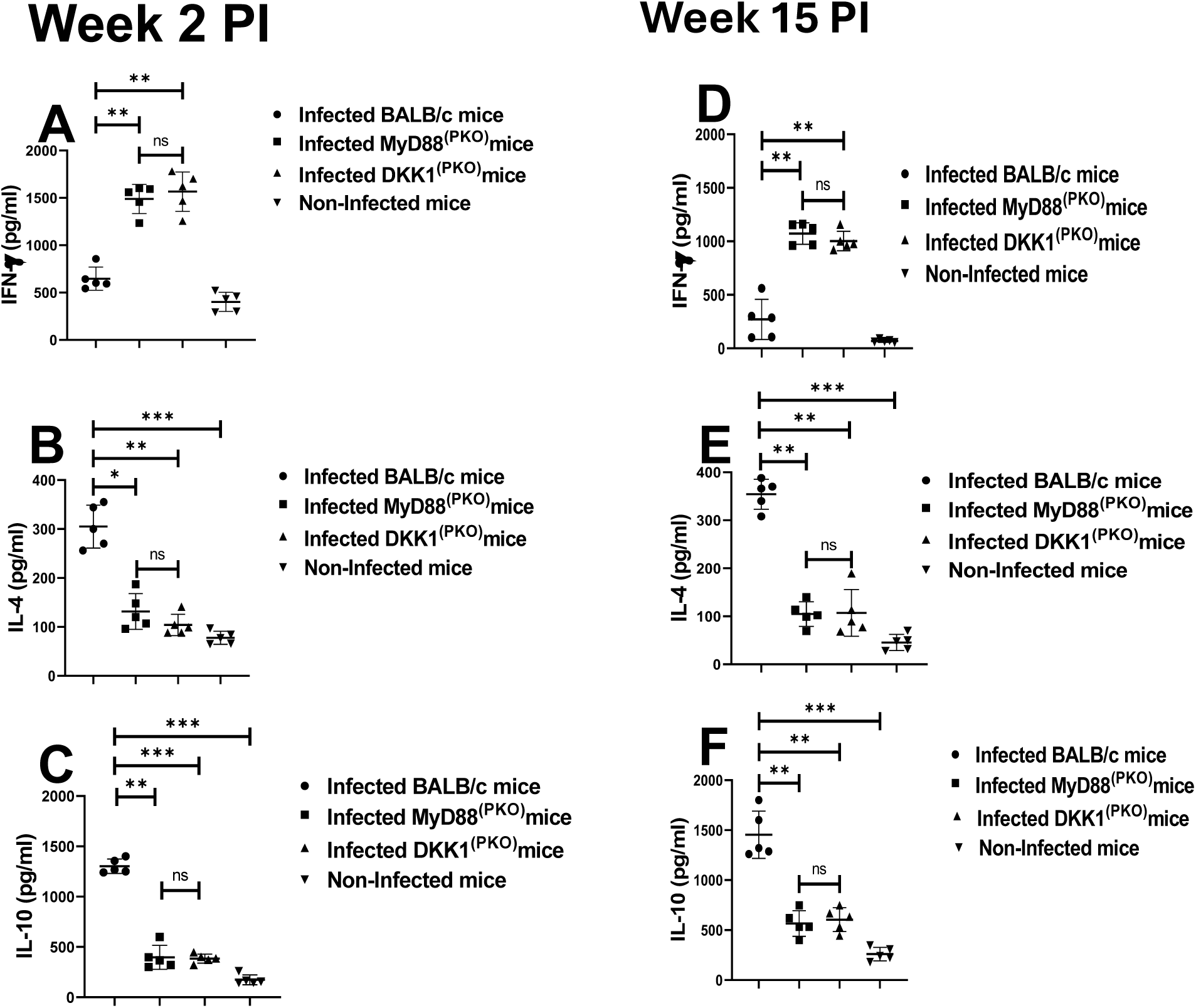
DKK1 promotes IL-10 induction in BALB/c-infected mice. BALB/c, MyD88^(PKO)^ and DKK1^(PKO)^ mice were challenged with infective metacyclic promastigote (2 x 10^6^ parasites, n = 5) of *L. major* via the footpad. Non- infected BALB/c mice (n = 5) were given 0.9% NaCl saline. Two or fifteen weeks post-infection, cells from draining and non-draining lymph nodes were isolated. Lymph node cells were incubated with SLAG (50 µg/ml derived from WT parasites). Cell culture supernatant samples obtained were analyzed by ELISA for cytokine production, as shown in the column graphs (**A- F**). In all the experiments, BALB/c infected and non-infected mice served as positive and negative controls, respectively. Results are presented as mean +/- SEM and are representative of two independent experiments. One-way ANOVA with Bonferroni’s *post hoc* test was performed to analyze the data *p < 0.05, **p < 0.01, ***p < 0.001, ‘ns’ indicates not significant (p > 0.05).

### Reduced IL-10-Th1 cells, CD4^+^IL-10^+^ and CD8^+^IL-10^+^ T-cells in MyD88^(PKO)^ and DKK1^(PKO)^ infected mice

Since MyD88^(PKO)^ and DKK1^(PKO)^ infected mice produce high levels of IFN-γ and low levels of IL-10 cytokines, we further evaluated the relative contribution of T-cell subsets in producing these cytokines. The results showed that the percentage of CD4^+^ T-cells was comparable in the draining lymph nodes of infected BALB/c, MyD88^(PKO)^ and DKK1^(PKO)^ mice (Fig. 7B). However, the percentage of CD8^+^T-cells was significantly lower in MyD88^(PKO)^ and DKK1^(PKO)^ infected mice compared to infected BALB/c mice (Fig. 7C). In addition, infected MyD88^(PKO)^ and DKK1^(PKO)^ mice showed a higher percentage of CD4^+^ and CD8^+^ T-cell-producing IFN-γ. Similarly, the MFI of IFN-γ-producing CD4^+^ and CD8^+^ T-cells was significantly higher in these mutant mice. However, there was an increased MFI and percentage of IL-10-producing CD4^+^ and CD8^+^ T- cells, while IFN-γ-producing CD4^+^ and CD8^+^ T-cells were low in the infected BALB/ mice (Fig. 7D, E, F, G, H, I, J & K) Interestingly, the percentage of IL-10-Th1 cells was elevated in BALB/c-infected mice compared to MyD88^(PKO)^ and DKK1^(PKO)^ infected mice (Fig. 7L). This suggests that DKK1 generated via the MyD88-dependent pathway promotes polarization of immune suppressive T-cells in BALB/c-infected mice.

**Figure 7:**
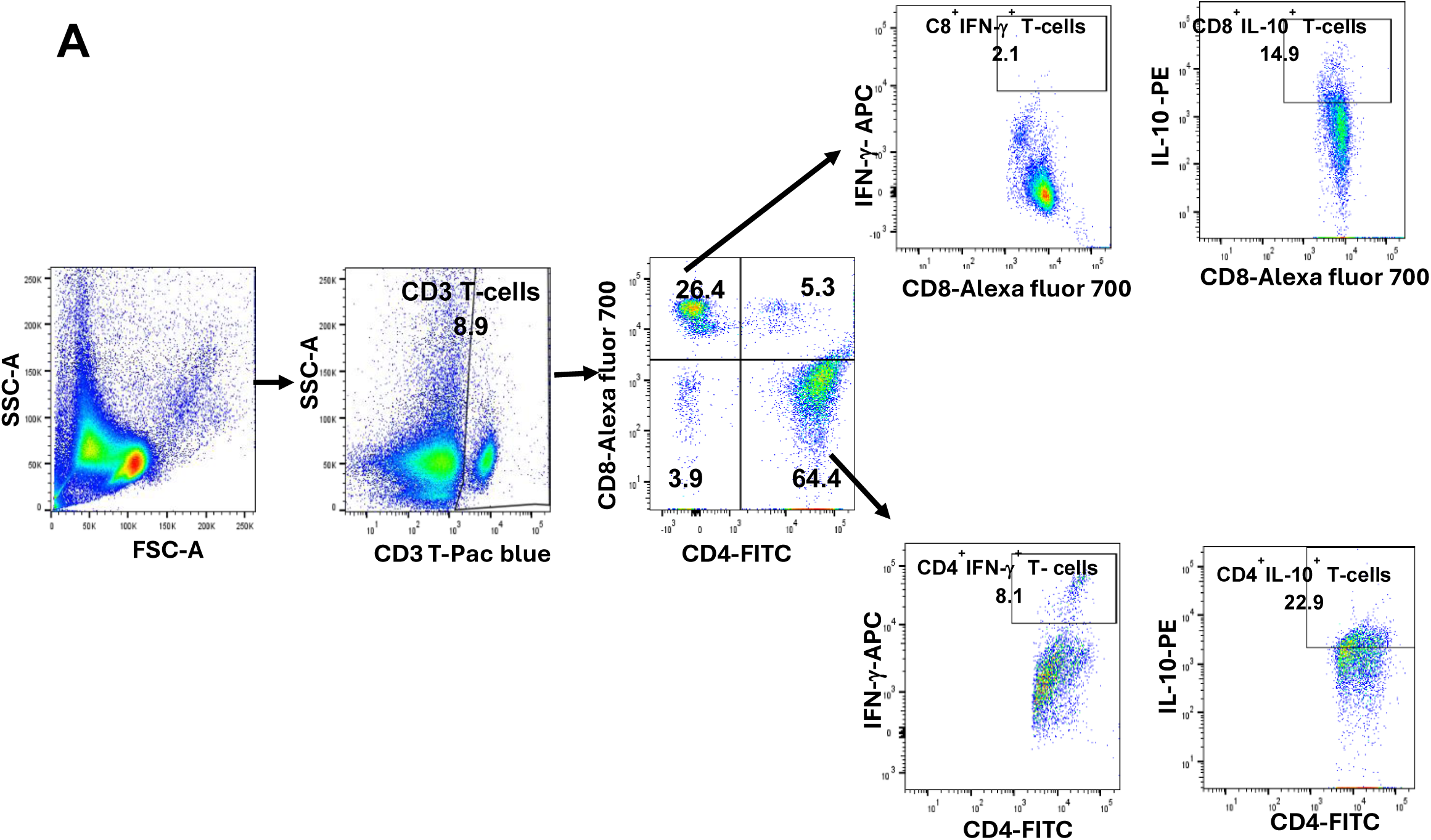
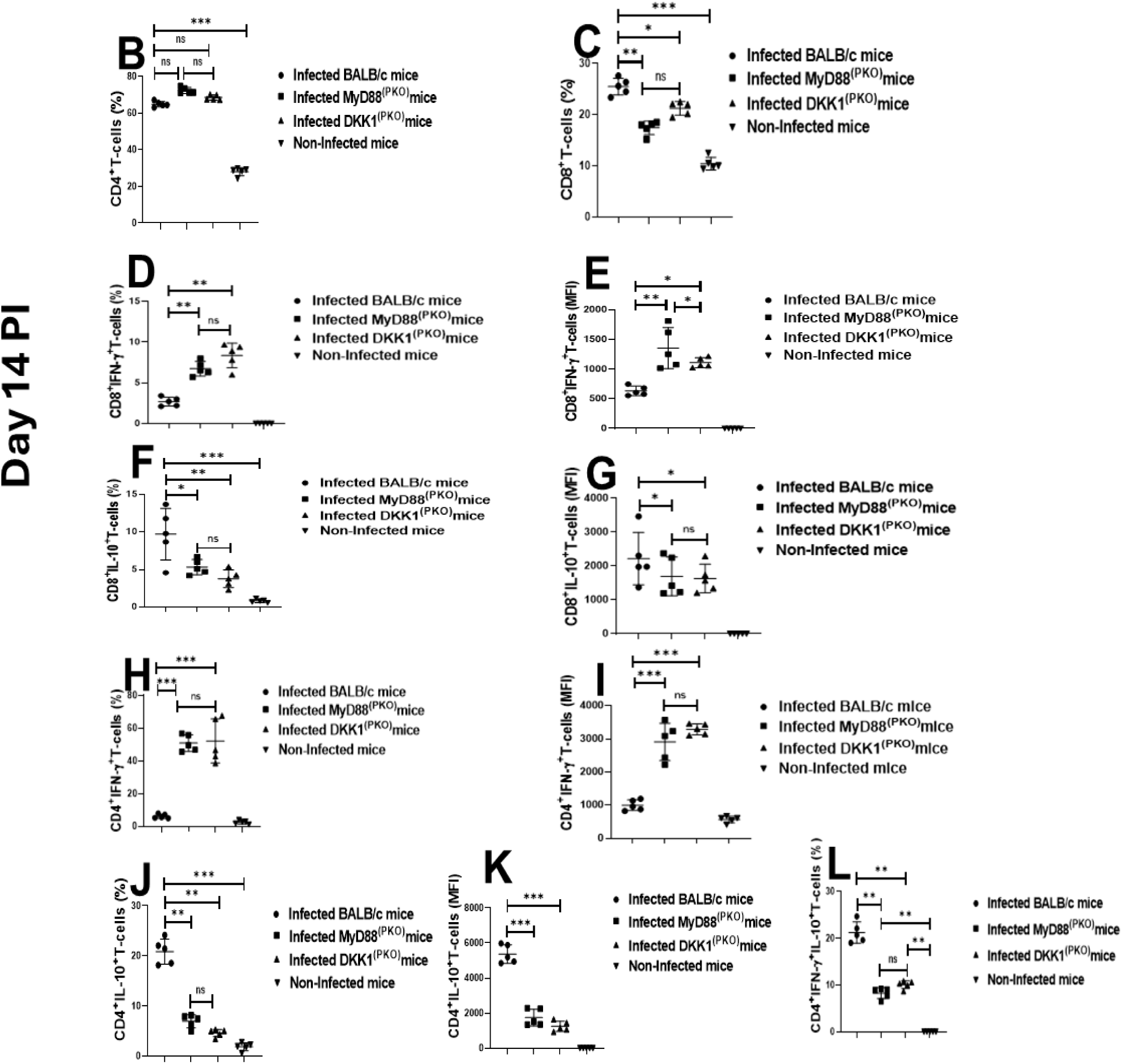
Impaired CD8^+^IL-10^+^IFN-γ^−^, CD4^+^IL-10^+^IFN-γ^−^ and CD4^+^IL-10^+^IFNg^+^ T-cells in MyD88^(PKO)^ and DKK1^(PKO)^ - infected mice on day 14 PI. BALB/c, MyD88^(PKO)^ and DKK1^(PKO)^ mice were challenged with infective metacyclic promastigote (2 x 10^6^ parasites, n = 5) of *L. major* via the footpad. Non-infected BALB/c mice (n = 5) were given 0.9% NaCl saline. Two weeks post-infection, the draining and non-draining lymph node cells were isolated. Lymph node cells were incubated with a cell stimulation cocktail for 5 hr. After a further 3 hr in BFA, cells were stained for intracellular IL-10 and IFN-γ. The stained lymph node cells from each mouse in BALB/c, MyD88^(PKO)^ and DKK1^(PKO)^ infected mice were determined for the percentage of CD4^+^ and CD8^+^ T-cells **(B & C)**, percentage and MFI of CD8^+^IFN-γ^+^ or CD8^+^IL-10^+^ T-cells (**D, E, F, & G**), percentage and MFI of CD4^+^IFN-γ^+^ or CD4^+^IL-10^+^ T-cells (**H, I, J & K**), and percentage of CD4^+^ IFN-γ^+^ IL-10^+^ T-cells (**L**) by flow cytometry. Representative flow cytometry dot plots showing the analyses CD4^+^ and CD8^+^ T-cells, CD8^+^IFN-γ^+^ or CD8^+^IL-10^+^ T-cells, CD4^+^IFN-γ^+^ or CD4^+^IL-10^+^ T-cells, and CD4^+^ IFN-γ^+^ IL-10^+^ T-cells performed on day 14 PI is indicated (**A**). A dot plot of each sample in all the experimental groups is presented in **Figure S4 (A, B, C, D, E and F).** Results are presented as mean +/- SEM. and are representative of triplicate experiments. Data analysis was done using one-way ANOVA with Bonferroni’s post hoc test *p < 0.05, **p < 0.01, ***p < 0.001, and “ns” indicates non-significant.

### rDKK1-treated dendritic cells display tolerogenic DC-10 phenotype

Tolerogenic DC-10 cells are potent inducers of IL-10-producing Th1 cells, well-known to block *Leishmania* resolution by suppressing Th1 cell-mediated immunity (Comi et al., 2018, Anderson et al., 2007). Tolerogenic DC-10 are usually detected using reduced expression of antigen-presenting molecules and elevated IL-10 production (Comi et al., 2018). Given that IL-10- producing Th1 cells are elevated in BALB/c-infected mice, we consider the possibility that DKK1 increases IL-10-producing Th1 cells by inducing tolerogenic DC-10. Thus, naïve dendritic cells were activated in vitro with r-TNF-α, r-DKK1, r-IL-10 or (r-TNF-α + rDKK1). Compared to r-TNF-activated dendritic cells, the expression and percentage of MHC II^+^, CD86^+^, and CD80^+^ molecules were impaired in r-DKK1-treated DCs (Fig. 8A, B, C, D, E, and F). Additionally, IL-10 production was elevated in r-DKK1-treated dendritic cells, while production of IL-12 by r-TNF-α treated dendritic cells was inhibited by r-DKK1 (Fig. 9A & B). The tolerogenic effect of rDKK1-treated dendritic cells was comparable to rIL-10-treated dendritic cells. These data suggest that DKK1 may regulate IL-10-Th1 cells by activation of tolerogenic DC-10 phenotype.

**Figure 8:**
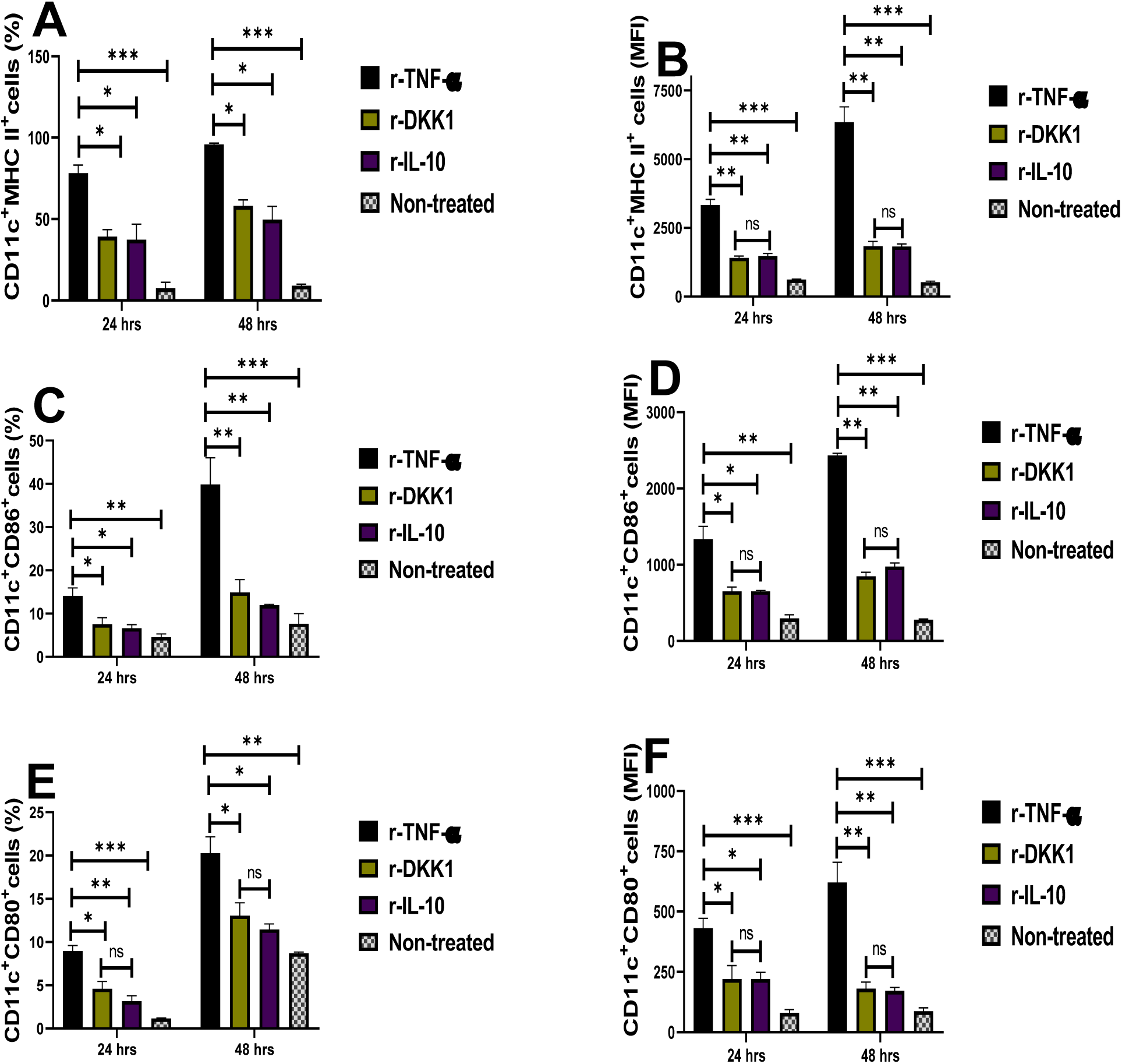
Minimal expression and percentage of MHC II^+^, CD86^+^ and CD80^+^ dendritic cells in the presence of recombinant DKK1. Monocyte-derived dendritic cells were incubated in rDKK1(100 ng/ml), rIL-10 (20 ng/ml) and rTNF-α (10 ng/ml). Cells harvested at 24 and 48 hrs post-incubation were used to determine the expression and percentage of MHC II+, CD86+, and CD80+ cells by flow cytometry. Representative flow cytometry dot plots generated 24 hrs post incubation showed the analysis of MHC II^+^, CD86^+^ and CD80^+^ as indicated in **Figure S5**. The percentage **(A, C, & E**) and MFI (**B, D & F**) of MHC II^+^, CD86^+^ and CD80^+^ in the different experimental conditions are shown in the bar graphs. Results are presented as mean (± SEM) and are representative of triplicate experiments. One-way ANOVA with Bonferroni’s post hoc test was performed to analyze the data (*p < 0.05, **p < 0.01; ***p < 0.001; and “ns” indicates non-significant.

**Figure 9:**
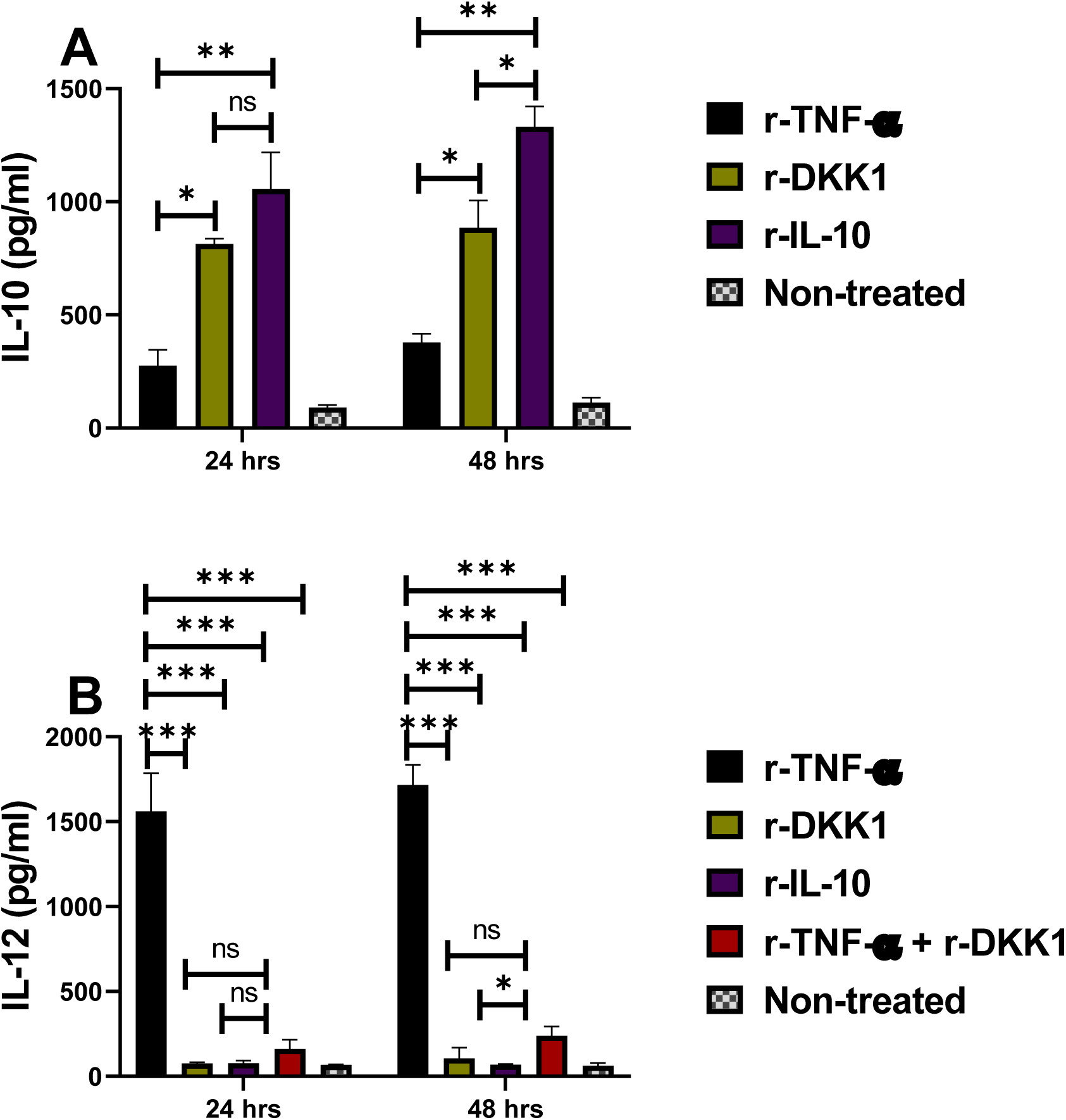
Elevated IL-10 and reduced IL-12 production in rDKK1-treated dendritic cells. Monocyte-derived dendritic cells were incubated in r-DKK1(100 ng/ml), r-IL-10 (20 ng/ml), r-TNF-α (10 ng/ml) or r-TNF-α + r-DKK1. Cell culture supernatants harvested 24 and 48 hrs post-incubation were used to determine IL-10 and IL-12 production by ELISA, as shown in the bar graphs **(A**) and (**B).** Results are presented as mean +/- SEM and are representative of triplicate experiments. One-way ANOVA with Bonferroni’s *post hoc* test was performed to analyze the data (*p < 0.05, **p < 0.01). ‘ns’ indicates not significant.

### Significant reduction in lesion size and parasite load in MyD88^(PKO)^ and DKK1^(PKO)^ infected mice

Lesion size was significantly reduced in MyD88^(PKO)^ and DKK1^(PKO)^ infected mice compared to infected BALB/c mice (Fig. 10A). In weeks 6 and 15 PI, the parasitic load was significantly decreased in MyD88^(PKO)^ and DKK1^(PKO)^ infected mice relative to infected BALB/c mice (Fig. 10B & C) and appeared to plateau, consistent with observed lesion development. Although previous studies have shown that administration of a DKK1 inhibitor can partially ameliorate disease, the disease still progressed (Chae et al., 2016). These data clearly demonstrate that platelet DKK1 released through MyD88 signaling is critical for disease progression, and release of DKK1 from platelets promotes parasite survival and proliferation in the infection site of BALB/c mice.

**Figure 10:**
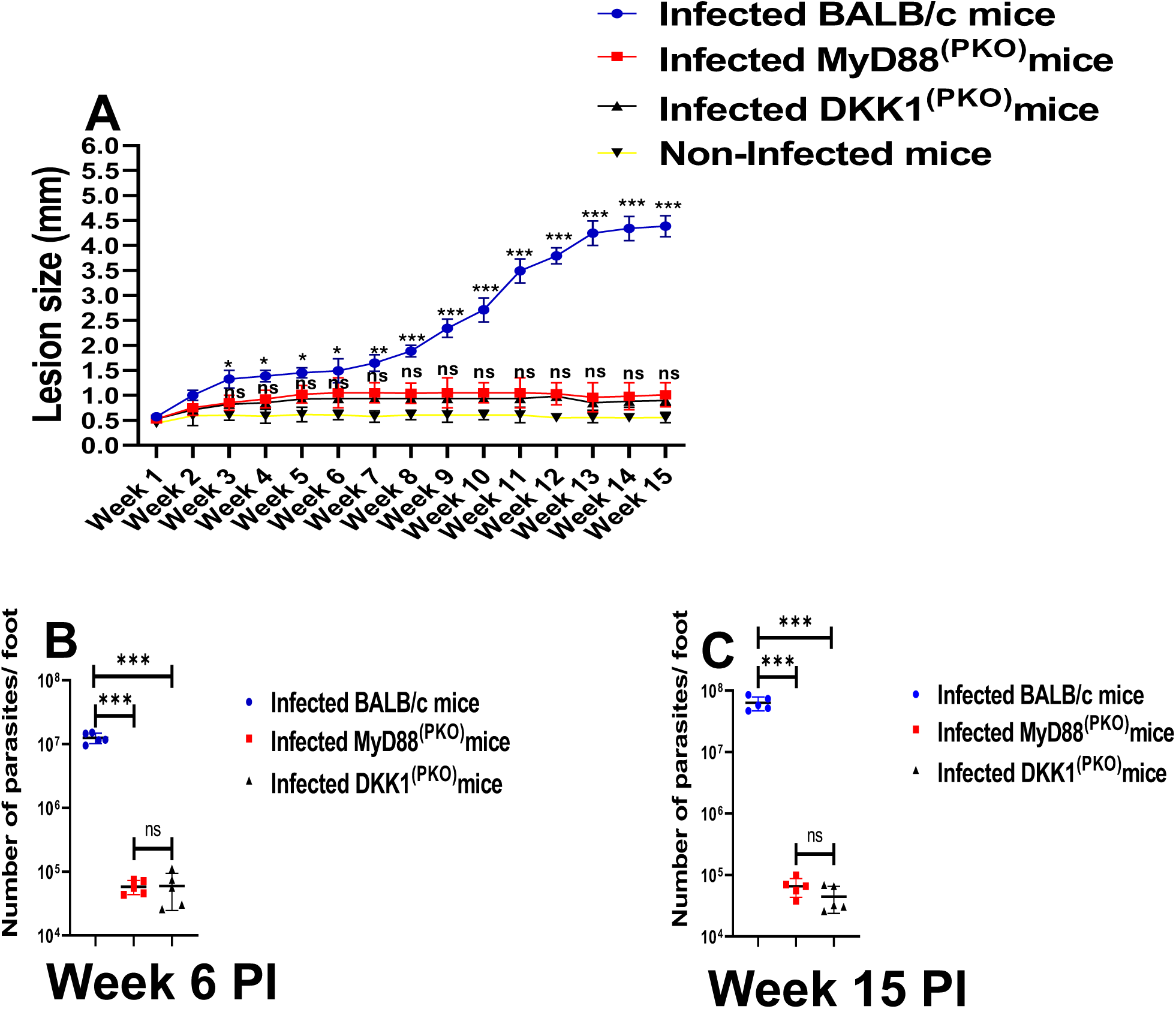
Lesion size and parasite burden decreased in MyD88^(PKO)^ and DKK1^(PKO)^ infected mice. The infected foot from each mouse in BALB/c, MyD88^(PKO)^ and DKK1^(PKO)^ infected mice were measured for lesion size weekly using a vernier caliper **(A)**, and parasite burden (at day week 6 and 15PI) was determined by limiting dilution assay **(B & C)**. Results are presented as mean +/- SEM. For Figure **(A),** mice in each infected group were compared with the non-infected group and data analysis was done using one-way ANOVA with Bonferroni’s post hoc test *p < 0.05; **p < 0.01, ***p < 0.001.

## Discussion

Platelets are established to aggregate in the periphery with leukocytes upon TLR1/2 engagement during *Leishmania* infection (Ihedioha et al., 2024, Chae et al., 2016). Aside from platelets, *Leishmania* species and their LPGs were reported to interact with TLR2 expressed on neutrophils, macrophages, dendritic cells or natural killer cells and modulate anti-leishmanial immune responses (Becker et al., 2003, Halliday et al., 2016, de Veer et al., 2003, Tolouei et al., 2013, Charmoy et al., 2007). The LPG-TLR2 interaction has reciprocal outcomes depending on which TLR2 co-receptor is up-regulated; the TLR2-TLR6 complex induces pro-inflammatory cytokines, whereas the TLR1-TLR2 heterodimer induces anti-inflammatory cytokines (Ghosh et al., 2024, Funderburg et al., 2011, Pandey et al., 2014). In addition, TLR6-silencing exacerbates the *L. major* infection, whereas TLR1-silencing improves protective response (Jafarzadeh et al., 2019). Immune pressure during *Leishmania* infection is maintained by IFN-γ-producing CD4 and CD8 T cells, IL-12-producing cells, and inducible nitric oxide synthase (iNOS) (Horta et al., 2012, Stobie et al., 2000). Impairment of these responses during latency has been shown in each case to promote parasite proliferation and the reappearance of lesions. However, the explanation as to why these control mechanisms fail to eliminate the parasite is not well understood. Our recent in vitro study showed that TLR1/2 plays a role in recognizing *Leishmania*-derived LPG and induction of DKK1 in activated platelets (Ihedioha et al., 2023). In turn, DKK1 potentiates Th2 cytokine expression and parasite survival (Chae et al., 2016). However, the mechanisms regulating how DKK1 is produced and modulates Th2 polarized responses are poorly understood. To gain a more comprehensive understanding of this process, mice with conditional deletion of MyD88 or DKK1 in platelets were used to evaluate whether the MyD88 activation signal and subsequent release of DKK1 from platelets regulate disease outcome by promoting a Th2 response in *Leishmania* infection. First, we demonstrated that the transmission of activation signals through MyD88 is crucial for platelet activation and early DKK1 production. Lack of platelet MyD88 or DKK1 resulted in impaired P-selectin expression. These findings highlighted MyD88 as the downstream protein regulating platelet activation and DKK1 production following interaction between LPG and platelet TLR1/2.

In this study, we have demonstrated that DKK1 produced via the TLR1/2-MyD88 dependent pathway elevates LPA and NPA formation, as well as the infiltration of activated neutrophils into the infection site of BALB/c-infected mice. This observation is consistent with our previous studies, which showed that DKK1 increases LPA and leukocytes in blood obtained from infected BALB/c mice (Chae et al., 2016). Also, our recent work has established that DKK1 signaling via its receptor (LRP6) elevates NPA formation and the migration of activated neutrophils to the infection site (Ihedioha et al., 2024). Further, these findings are in accordance with previous studies, which showed that activated platelets secrete platelet-derived growth factor, required for infiltration of a subpopulation of effector monocytes to the site of *Leishmania* infection (Goncalves et al., 2011). These findings provide direct evidence that DKK1 produced via the MyD88- dependent pathway plays a pivotal role in modulating systemic and local inflammatory responses through the Wnt3a pathway. The comparable LPA and NPA formation in all infected mice on day 14 PI suggests that as the expression level of LPG required for DKK1 production declines in the mammalian host with transformation of the parasite into the amastigote stage, LPG plays a minimal critical role in the formation of platelet-neutrophil aggregates needed for neutrophil infiltration to the infection site.

The impact of Th1 versus Th2 immunity on intracellular infections is attributed to classical versus alternative activation of macrophages leading to resistance or susceptibility to infection (Carneiro et al., 2020). Our previous paper demonstrated that DKK1 is an important regulator of leukocyte infiltration and polarization of immune responses in pathological type 2 cell-mediated inflammation (Chae et al., 2016). Because DKK1 produced via the MyD88 activation signal promotes the migration of leukocytes to the infection site, we tested the hypothesis that the Th2 response in infected BALB/c mice may be mediated by DKK1 produced via a MyD88-dependent pathway. The elevated production of IL-10 and IL-4 in BALB/c-infected mice, in contrast to the mice with platelets deficient in MyD88 or DKK1, suggests that DKK1 released following activation of MyD88 promotes Th2 cytokine production. This effect is associated with elevated IL-10-producing CD4 and CD8T-cells, cDC2 dendritic cells and M2 macrophages. In the context of *Leishmania* infection, a Th2 response is considered detrimental, as it hampers responses controlling parasite survival/growth and promotes disease progression (Scott and Novais, 2016). Also, M2 macrophages and cDC2 dendritic cells have been linked to the development of pathology; parasites survive and multiply within M2 macrophages (Lee et al., 2018). The impaired production of Th2 cytokines (specifically IL-10 and IL-4) in DKK1^(PKO)^ and MyD88^(PKO)^ infected mice confirms that the Th2 response in BALB/c infected mice may be mediated by DKK1 produced through the MyD88-dependent pathway.

Aside from the Th2 cell-polarizing conditions that underlie the increased susceptibility of BALB/c mice to *L. major*, healing forms of cutaneous leishmaniasis in human and mouse models are promoted by the presence of ongoing Th1 responses. Dendritic cells are important in determining the host response, serving as both host cells, APCs and sources of cytokines directing a Th1 or Th2 response. These interactions between dendritic cells and *Leishmania* parasites are complex and involve paradoxical functions that can block or activate T-cell responses, resulting in the control of infection or disease progression (Soong, 2008). The magnitude and profile of dendritic cell activation vary greatly, depending upon the cytokines released early in the microenvironment and dendritic cell subsets (Soong, 2008). It has been demonstrated that BALB/c mice produce IL-4 early following infection. While IL-4 is known to induce Th2 responses (Himmelrich et al., 2000, Radwanska et al., 2007), previous studies have also shown that the administration of recombinant IL-4 within 8 hours post-infection with *L. major* led to increased IL-12 mRNA expression by dendritic cells in vivo and rendered mice resistant to infection (Biedermann et al., 2001). Further, IL-4 has been shown to inhibit IL-10 production by dendritic cells (Yao et al., 2005). During *L. major* infection in BALB/c mice, abrogation of IL-4 receptor alpha on dendritic cells blocked IL- 12 but increased IL-10 production due to impaired dendritic cell instruction, resulting in increased susceptibility (Hurdayal et al., 2013). Further studies confirmed CD11b^+^Ly6C^+^ inflammatory dendritic cells as the most highly infected dendritic cell subset responsible for hosting and disseminating *L. major* parasites in CD11c^cre^IL-4Rα^−/lox^ mice (Hurdayal et al., 2019). Notably in the current study, the level of CD11c^+^CD11b^+^ cells are selectively diminished in DKK1 deficient MyD88^(PKO)^ and DKK1^(PKO)^ infected mice.

In contrast, IL-10 compromises healing responses in intensity or function, acting through effects on T cells, macrophages and dendritic cells (Anderson et al., 2007, Nagase et al., 2007, Couper et al., 2008, Abdoli et al., 2017, Saraiva et al., 2020). Adoptive transfer studies have shown that IL- 10 produced by the CD4^+^ CD25^−^ Foxp3^−^ T cells suppressed the healing response in cutaneous leishmaniasis (Anderson et al., 2007). In addition, it was shown that the majority of the CD4 T cells in the dermis and a smaller subset of CD4 T cells in the draining node, were CD4 T cells that produce both IL-10 and IFN-γ (Anderson et al., 2007). It has been found that DC-10 cells direct the development of IL-10 producing Th1 cells (Comi et al., 2018). In particular, IL-10 inhibits the up-regulation of antigen-presenting molecules and IL-12 production, thus inhibiting the capacity of dendritic cells to generate Th1 responses (Cavani et al., 2000). IL-10 exerts its actions through interaction with the IL-10 receptor, resulting in the activation of a series of intracellular signaling molecules, including STAT proteins (Cassatella et al., 1999, Riley et al., 1999). In the current study, we have demonstrated that the presence of platelet-derived DKK1 in vivo promotes the development of tolerogenic DC-10 dendritic cells and M2 macrophages. This effect on dendritic cells can be direct, as demonstrated in vitro that rDKK1 induces tolerogenic DC-10 cells and blocks activation by TNFα. IL-10 has been identified as a primary factor that can block dendritic cell maturation induced by various stimuli (Caux et al., 1994, Buelens et al., 1997, Allavena et al., 1998, Corinti et al., 2001). Notably, the tolerogenic effect of r-DKK1 and r-IL-10 treated dendritic cells in vitro were comparable. As DC-10 cells are important for Th1-IL-10 cell differentiation, this suggests that both IL-10 and DKK1 contribute to the development of Th1-IL-10 cells by limiting dendritic cell maturation and their capacity to initiate Th1 responses, resulting in parasite proliferation in BALB/c infected mice.

Our study has highlighted the importance of DKK1 produced through the platelet-MyD88 signaling pathway in dendritic cell polarization. This has been shown to be critical for regulating IL-10 levels and ultimately influencing the differentiation of Th2 and Th1-IL-10 cells. Consequently, DKK1 promotes disease progression and parasite survival in M2 macrophages by inducing Th2 and Th1-IL-10 responses. Thus, inhibiting the TLR1/2-MyD88 pathway or DKK1 production in platelets may be an attractive approach to limit parasite proliferation in *Leishmania* infection.

## Materials and methods

### Mice

WT BALB/c mice (4 weeks old) were purchased from the Jackson Laboratory. Sperm from mice heterozygous for MyD88 flox and PF4-Cre recombinase was used for in vitro fertilization, and pups carrying a heterozygous floxed allele of MyD88 and PF4-Cre were backcrossed to female BALB/c mice for eight generations. Thereafter, the heterozygous MyD88 floxed mice were intercrossed with mice expressing Cre recombinase under the control of the PF4 promoter to generate mice in which MyD88 was selectively deficient in platelets (MyD88^PKO^). PF4-Cre- DKK1 deficient mice (DKK1^PKO^) were generated by specific deletion of DKK1 in megakaryocytes and platelets. All experiments were conducted with age-matched littermates, unless specified otherwise, and all mice were housed at the University of Arizona Animal or University of Nebraska Medical Center Care Facilities. The mouse protocols were approved by the University of Arizona in accordance with the Association for Assessment and Accreditation of Laboratory Animal Care International (AAALAC). Mice strains were transferred to the University of Nebraska Medical Center (UNMC) and utilized under approved protocols.

### Genotyping of mouse lines used in the study

Genotyping of all the knockout mice was performed through standard PCR procedures (Jacquot et al., 2019). Genomic DNA from Proteinase K-digested tail tissue was extracted using a buffer composed of 50 mM Tris-HCl (pH 8.0), 100 mM NaCl, 1% SDS, and 100 mM EDTA (pH 8.0). The primers used for the PCR reactions are the following: mDKK1 Flox1 (5^’^-AGAACTAACCC CGGCCCCACAGCAGA-3^’^); mDKK1 Flox2 (5^’^-CTCCTCAGGGAAGACAACAAAGCCG- 3^’^); MyD88-O1MR9481 F (5^’^-GTTGTGTGTGTCCGACCGT-3^’^); MyD88-01MR9482 R (5^’^- GTCAGAAACAACCACCACCATGC-3^’^); PF4-Cre F (5^’^-CCCATACAGCACACCTTTTG-3^’^); PF4-Cre R (5^’^-TGCACAGTCAGCAGG TT -3^’^).

### Parasite strains and infection protocol

The WT L. *major* LV39 clone 5 was maintained at 26°C in M199 culture medium (Thermo Fisher Scientific) supplemented with 20% heat-inactivated FBS (Thermo Fisher Scientific), 20 mM HEPES (Sigma-Aldrich) and 50 µg/ml gentamycin (Thermo Fisher Scientific). Prior to using parasites for infection, metacyclic promastigotes were isolated from stationary-phase cultures by centrifugation through a 45% and 90% Percoll® gradient (GE Healthcare) as previously described (Ahmed et al., 2003, Jara et al., 2022). Live parasites were isolated at the 90%-45% interface. Metacyclic promastigotes were washed three times in cold phosphate-buffered saline (PBS-Thermo Fisher Scientific) by centrifugation, resuspended in PBS at 2X10^8^/ml and 10 μl containing 2x10^6^ metacyclic promastigotes were injected into the top of the right hind footpad.

### Platelet preparation and P-selectin expression

P-selectin expression was assessed in platelets obtained from infected BALB/c, DKK1^(PKO)^, MyD88^(PKO)^, and non-infected mice, as previously described with minor modifications (Ihedioha et al., 2023). Briefly, blood was drawn by maxillary bleeding into tubes containing 3.2% citrate buffer (G-Biosciences) from infected BALB/c, DKK1^(PKO)^, MyD88^(PKO)^ and non-infected mice at days 3 and 14 PI. In brief, blood was diluted with 250 μL modified Tyrode’s-HEPES buffer (134 mM NaCl, 0.34 mM Na_2_HPO4, 2.9 mM KCl, 12 mM NaHCO_3_,20 mM HEPES, 5 mM glucose, and 1 mM MgCl_2_, pH7.3) and centrifuged at 250 x g for 15 minutes at room temperature. Platelet-rich plasma was removed, and platelets were then isolated by centrifugation at 900 × g for 30 min. Isolated platelets (10x10^6^/ml) were washed at 550 x g for 10 minutes in the presence of PGE1 (140 nM; Sigma Aldrich) and indomethacin (10 mM; Thermo Fisher Scientific). Platelets were resuspended to the required density in modified Tyrode’s HEPES buffer (Sigma-Aldrich) and rested for 30 minutes at 37°C in the presence of 10 mM indomethacin before staining. Staining for P-selectin expression was done using PE-conjugated CD41 and APC-conjugated CD62P. Stained platelets (1 × 10^6^/ml in Tyrode’s-HEPES buffer; 100 μL) were analyzed using an LSR II flow cytometer, and the analysis was performed using FlowJo software. A total of 10,000 events were collected per sample. The mean fluorescent intensity of P-selectin was compared among the different groups.

### Cell isolation from infected footpad and lymph node

Neutrophils, dendritic cells, and macrophages were isolated from the footpad using an established protocol (Hameed et al., 2023). Briefly, infected footpads were cut right above the ankle and deskinned. Crosswise cutting was performed in small increments, starting from the bottom of the foot and moving towards the ankle, and minced using toothed forceps. To obtain more cells in the non-infected mice, PBS was injected into the two feet of each mouse, and the cells on both feet were harvested. Additionally, the number of mice in the non-infected group increased from 5 to 10 mice, and the cells were then pooled for analysis. Additionally, cells were harvested from both draining and non-draining lymph nodes, and a single-cell suspension was prepared. Cells were separated from tissues using a cell strainer (70 μm pore size- Thermo Fisher Scientific), centrifuged at 300 x g for 7 minutes, and resuspended in FACS buffer for cell staining.

### Analysis of dendritic and macrophage subtypes

The abundance of cDC1 and cDC2 subtypes in dermal dendritic cells obtained from the footpads of infected BALB/c, DKK1^(PKO),^ MyD88^(PKO),^ and non-infected BALB/c mice was assessed on day 14 post-infection using PE-conjugated CD11c, FITC-conjugated CD11b, and PECy5.5-conjugated CD8α antibodies. The cDC1 and cDC2 subtypes were identified by measuring the percentage of CD11b^-^CD8α^+^ and CD11b^+^CD8α^-^ cells within the CD11c^+^ gated population, respectively. In addition, the frequency of cDC1 and cDC2 was assessed using flow cytometry. The abundance of M1 and M2 subtypes in dermal macrophages obtained from the footpads of infected BALB/c, DKK1^(PKO)^, MyD88^(PKO)^, and non-infected BALB/c mice was assessed on day 14 post-infection using APC-conjugated F4/80, PE-conjugated CD38, and Alexa Fluor 700-conjugated CD206. The M1 and M2 subtypes were determined by measuring the percentage of CD38^+^CD206^-^ and CD38^-^ CD206^+^ cells within the F4/80^+^gated population, respectively.

### Neutrophil-platelet aggregate (NPA) and leukocyte-platelet aggregate (LPA) formation

Neutrophil-platelet aggregate assessment was performed as described previously with minor modifications (Ihedioha et al., 2023, Ihedioha et al., 2024). Briefly, cells (1x10^6^/ml) isolated from footpads of infected BALB/c, MyD88^(PKO),^ DKK1^(PKO)^ and non-infected BALB/c mice on days 3 and 14 PI, were stained with FITC conjugated Ly6G (BioLegend), and PE-conjugated CD41 (BioLegend) antibodies for 15 mins in the dark at room temperature. Similar staining was performed to assess leukocyte-platelet aggregates using cells (1 x 10^6^/ml) isolated from blood. Isolated cells were stained with Pacific blue-conjugated CD45 (BioLegend) and PE-conjugated CD41 (BioLegend). Samples were acquired using LSR II flow cytometry within 4 to 6 hours, and analysis was performed with FlowJo software. Live gating was performed on leukocyte-sized events to exclude single platelets. Leukocytes were identified by their forward and side scatter characteristics, as well as CD45 expression. Neutrophils were identified by their forward and side scatter characteristics and Ly6G expression. The Ly6G^+^ and CD41^+^ subpopulations identified NPA, while CD45^+^ and Ly6G^+^ subpopulations identified LPA.

### CD11b^+^ and MHC class II^+^ neutrophils

Assessment of neutrophil activation (CD11b^+^ and MHC class II^+^ cells) was performed by minor modification of previously described methods (Ihedioha et al., 2024). Briefly, cells isolated from footpads of infected BALB/c, MyD88^(PKO),^ DKK1^(PKO)^ on days 7 and 14 PI and non-infected BALB/c mice, were stained with Pacific-blue conjugated Ly6G (BioLegend), FITC conjugated CD11b (eBioscience), and PE-conjugated MHC class II (eBioscience) antibodies for 15 mins in the dark at room temperature. Stained samples were evaluated within 2 hours by LSR II flow cytometry and analyzed using FlowJo software. Neutrophils were identified by their forward and side scatter characteristics and Ly6G expression. The CD11b^+^ and MHC class II^+^ subpopulation identified activated neutrophils.

### Myeloperoxidase positive (MPO^+^) neutrophils

Activated neutrophils were further determined by measuring myeloperoxidase MPO^+^ neutrophils using the established flow cytometry intracellular cell staining method with minor modifications (Ihedioha et al., 2024). Briefly, cells (1x10^6^/ml) generated from the footpad of infected BALB/c, MyD88^(PKO),^ DKK1^(PKO)^ on days 3 and 14 PI and non-infected BALB/c mice, were fixed at room temperature in a tube with BD Cytofix^TM^ Fixation Buffer (BD Biosciences) that has been prewarmed to 37°C. The cells were then washed and permeabilized with BD Perm/Wash^TM^ Buffer (BD Biosciences). The permeabilized cells were stained in the dark at room temperature in BD Perm/Wash^TM^ Buffer with pacific-blue-conjugated Ly6G (day 3 PI), FITC-conjugated Ly6G (day 14 PI) and with either Alexa Fluor 647 mouse IgG1 isotype control or Alexa Fluor 647-conjugated myeloperoxidase antibody (BD Biosciences). Data acquisition and analysis were done using LSR II flow cytometry and FlowJo software, respectively.

### Plasma DKK1 ELISA

Plasma was collected from blood drawn from infected BALB/c, DKK1^(PKO)^, MyD88^(PKO)^ and non- infected mice as previously described (Ihedioha et al., 2023). Plasma was collected following blood centrifugation at 900 x g for 30 minutes. The concentration of DKK1 in plasma was determined by Enzyme-linked immunosorbent assays using a mouse DKK1 ELISA kit (Thermo Fisher Scientific) according to the manufacturer’s protocol.

### Ex-vivo lymph node cell stimulation and cytokine determination using ELISA

Single-cell suspensions from draining and non-draining lymph nodes of infected BALB/c, MyD88^(PKO)^, DKK1^(PKO)^ and non-infected BALB/c mice were activated with soluble leishmania antigen (SLAG) at a concentration of 50 µg/ml. Cells (1 × 10^6^ cells/mL) were cultured in a complete RPMI 1640 medium (Thermo Fisher Scientific) supplemented with 10% FBS, 1% penicillin/streptomycin (Gibco) and 10 mM HEPES at 37 °C under 5 % CO_2._ The supernatant was collected 72 hrs post-incubation with SLAG and stored at -80°C for cytokine analysis. The concentrations of IL-4, IL-10, and IFN-*γ* in the supernatant were determined using mouse Th1 and Th2 cytokine panel ELISA kit (eBioscience) according to the manufacturer’s protocol. For SLAG preparation, *L. major* parasites (5 × 10^8^ parasites/ml in M199 medium without FCS) were subjected to five repeated freeze-thaw cycles at -80°C and room temperature, respectively.

### Intracellular staining for IFN-γ and IL-10 in CD4^+^ and CD8^+^T cell

On day 14 PI, single-cell suspension obtained from draining and non-draining lymph node cells of non-infected BALB/c, infected BALB/c, MyD88^(PKO)^ and DKK1^(PKO)^ mice were stimulated with a cell stimulation cocktail (BD Biosciences; 2 ul/ml) for 5 hr, and BD Golgi Plug (BD Biosciences; 1 μl/ml) added for the final 3 hr. Cell stimulation was performed in complete RPMI 1640 medium supplemented with 10% FBS, 1% penicillin/streptomycin and 10 mM HEPES. The cells were surface-stained with the following antibodies (from eBioscience): PacBlue-conjugated anti-CD3, FITC-conjugated anti-CD4, and Alexa Fluor-conjugated anti-CD8 antibodies. Intracellular staining was performed as described in the manufacturer’s protocol (eBioscience) using APC-conjugated anti-IFN-γ (eBioscience) or PE-conjugated anti-IL-10 antibodies (BD Biosciences). PE mouse IgG1(BD Biosciences) and APC mouse IgG2a antibodies (eBioscience) served as isotype controls. Data acquisition and analysis were done using LSR II flow cytometry and FlowJo software, respectively.

### Generation of monocyte-derived dendritic cells, in vitro cell stimulation and functional analysis

CD11b^+^ monocytes were isolated and purified from mouse bone marrow cells using CD11b MicroBeads UltraPure mouse magnetic bead (Miltenyi Biotec) according to the manufacturer’s protocol. Isolated monocytes (1 × 10^6^ cells/mL) were differentiated into dendritic cells by 6 days of culturing with RPMI 1640 medium supplemented with 10% FBS, 1% penicillin/streptomycin, 10 mM HEPES, 50 µM 2-mercaptoethanol (Thermo Fisher Scientific), 20 ng/mL of IL-4 (Thermo Fisher Scientific) and 100 ng/mL of GM-CSF (Thermo Fisher Scientific) in 6-well tissue culture plates (Corning Costar). Cells were confirmed dendritic cells by flow cytometric analyses for CD11c^+^ expression. Immature dendritic cells were allowed to rest for 24 hours in complete RPM1 1640 media without cytokines to remove the residual effect of GM-CSF. Rested immature dendritic cells were stimulated with 100 ng/ml of recombinant DKK1 (R & D Systems), 20 ng/ml of recombinant IL-10 (Thermo Fisher Scientific), 10 ng/ml of recombinant TNF-α (Thermo Fisher Scientific) or recombinant TNF-α plus DKK1. The supernatant was collected at 24 and 48 hrs post-incubation and stored at -80◦C for cytokine analysis. The concentration of IL-10 and IL- 12p40 in the supernatant was determined using a mouse IL-12p40 as well as a Th1 and Th2 cytokine panel ELISA kit (eBioscience) according to the manufacturer’s protocol. For the detection of MHC II and co-stimulatory molecules, dendritic cells were stained with antibodies from eBioscience: FITC-conjugated CD11c, PE-conjugated MHC II, APC-Cy7-conjugated CD80, and V450-conjugated CD86. Data acquisition and analysis were done using LSR II flow cytometry and FlowJo software, respectively.

### Estimation of parasite burden and lesion size

Parasite burden was estimated by limiting dilution analysis at weeks 6 and 15 PI as previously described (Titus et al., 1985). In brief, cells from the infection site were suspended in Schneider medium supplemented with 20% heat-inactivated FBS and 1% Penicillin-streptomycin. After counting the cells, ten serial dilutions were prepared; for each dilution, eight wells (100 μL) were set up in 96-well microtiter plates. The cells were incubated at 26 °C for 10 days. The total number of positive wells (presence of motile promastigotes) and negative wells (absence of motile promastigotes) were identified by an inverted light microscope. The number of viable parasites in the infection site of each mouse was determined from the highest dilution at which promastigotes could be grown.

Lesion size was monitored weekly by using a Vernier caliper to measure the thickness of the infected hind footpad and compare it with that of the non-infected hind footpad. Lesion size = Size of infected footpad - Contralateral uninfected footpad (mm).

### Statistical analysis

Statistical tests were noted in figure legends. All data were shown as means ± SEM. Analyses were performed using GraphPad Prism 10 software. Statistical significance was identified with *P < 0.05; *P < 0.01; **P < 0.001; and ns, not significant. Graphical abstract was created with https://Biorender.com.

## Online supplemental material

This article includes five supplemental materials that demonstrate flow cytometry gating strategies, representative plots, and supplemental experimental figures. Fig. S1, related to Fig. 3, presents the analysis of LPA and NPA in non-infected BALB/c, infected BALB/c, MyD88^(PKO)^ and DKK1^(PKO)^ mice on days 3 and 14 PI. Fig. S2, relating to Fig. 4, presents the analysis of MPO^+^, CD11b^+^ and MHC class II^+^ neutrophils in non-infected BALB/c, infected BALB/c, MyD88^(PKO)^ and DKK1^(PKO)^ mice on days 3 and 14 PI. Fig. S3, relating to Fig.5, indicates the analysis of macrophages and dendritic cell subsets in non-infected BALB/c, infected BALB/c, MyD88^(PKO)^ and DKK1^(PKO)^ mice on days 7 and 14 PI. Fig. S4, relating to Fig.7, shows the analysis of CD8^+^IL- 10^+^, CD4^+^IL-10^+^ and CD4^+^IL-10^+^IFNγ^+^ T-cells in non-infected BALB/c, infected BALB/c, MyD88^(PKO)^ and DKK1^(PKO)^ mice on day 14 PI. Fig. S5, relating to Fig.8, addresses the analysis of MHC II^+^, CD86^+^ and CD80^+^ dendritic following in vitro treatment with rTNF-α, rDKK1 and rIL-10.

## Data availability

All data associated with this study are available in the paper or the online supplemental material.

## Acknowledgments

We thank INFRAFRONTIER/EMMA (www.infrafrontier.eu, PMID: 25414328) for providing the fertilized embryos (DKK1^fl/fl-/-^). We also appreciate the UNMC Flow Cytometry Research Facility (The UNMC Flow Cytometry Research Facility is administrated through the Office of the Vice Chancellor for Research and supported by state funds from the Nebraska Research Initiative (NRI) and The Fred and Pamela Buffett Cancer Center’s National Cancer Institute Cancer Support Grant (P30 CA036727). Major instrumentation has been provided by the Office of the Vice Chancellor for Research, The University of Nebraska Foundation, the Nebraska Banker’s Fund, and the NIH-NCRR Shared Instrument Program). In addition, we acknowledge Dr. Joann B Sweasy, Dr. Stephen M Beverley, Dr. Jennifer M Lundberg, and Dr. Eddie Wook-Jin Chae for their excellent suggestions and helpful advice. This work was supported by AI-137060 (awarded to Alfred Bothwell).

