## Supplemental Figures for "Platelet DKK1 promotes tolerogenic dendritic cells and non-healing responses in cutaneous leishmaniasis"

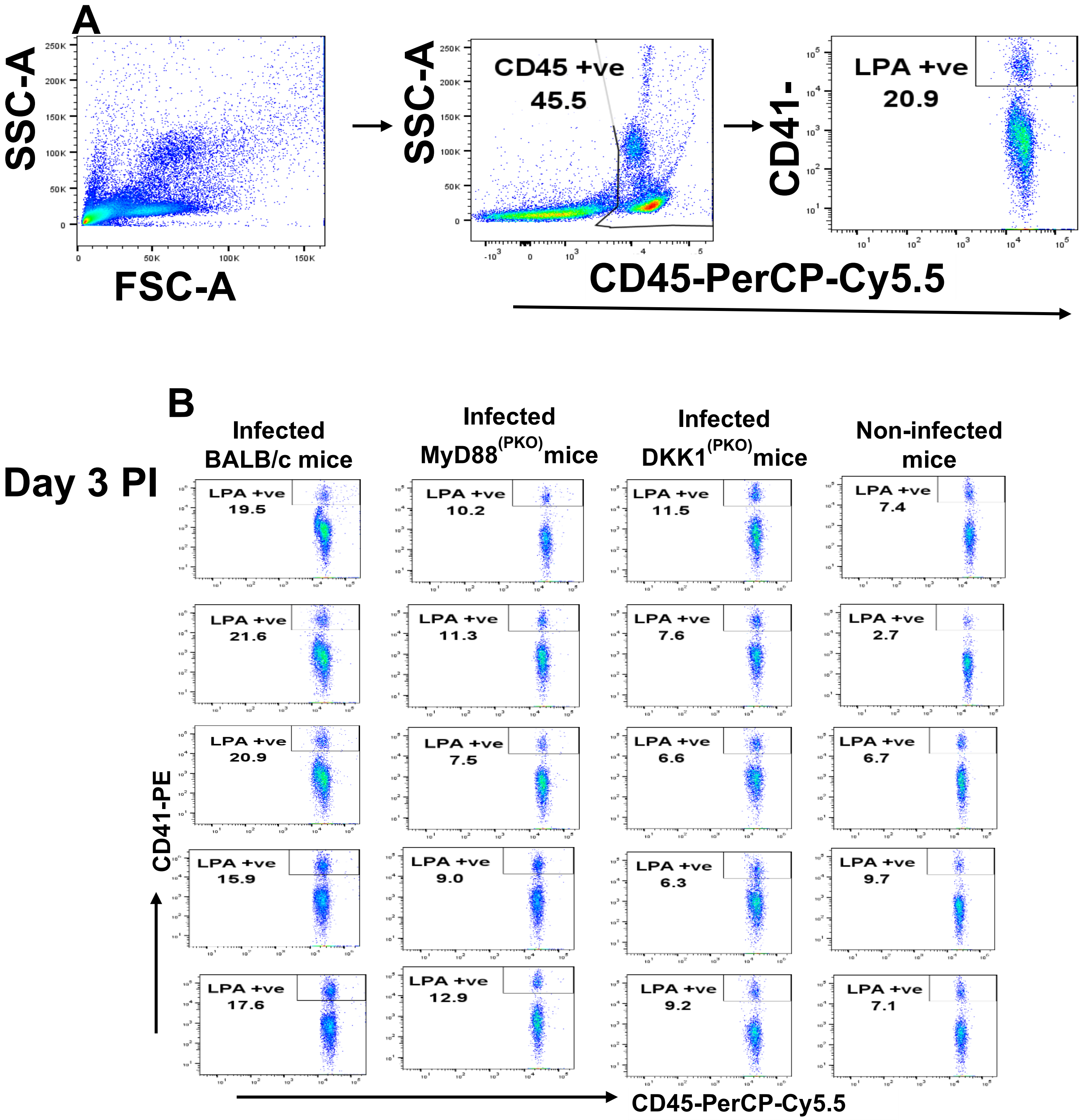

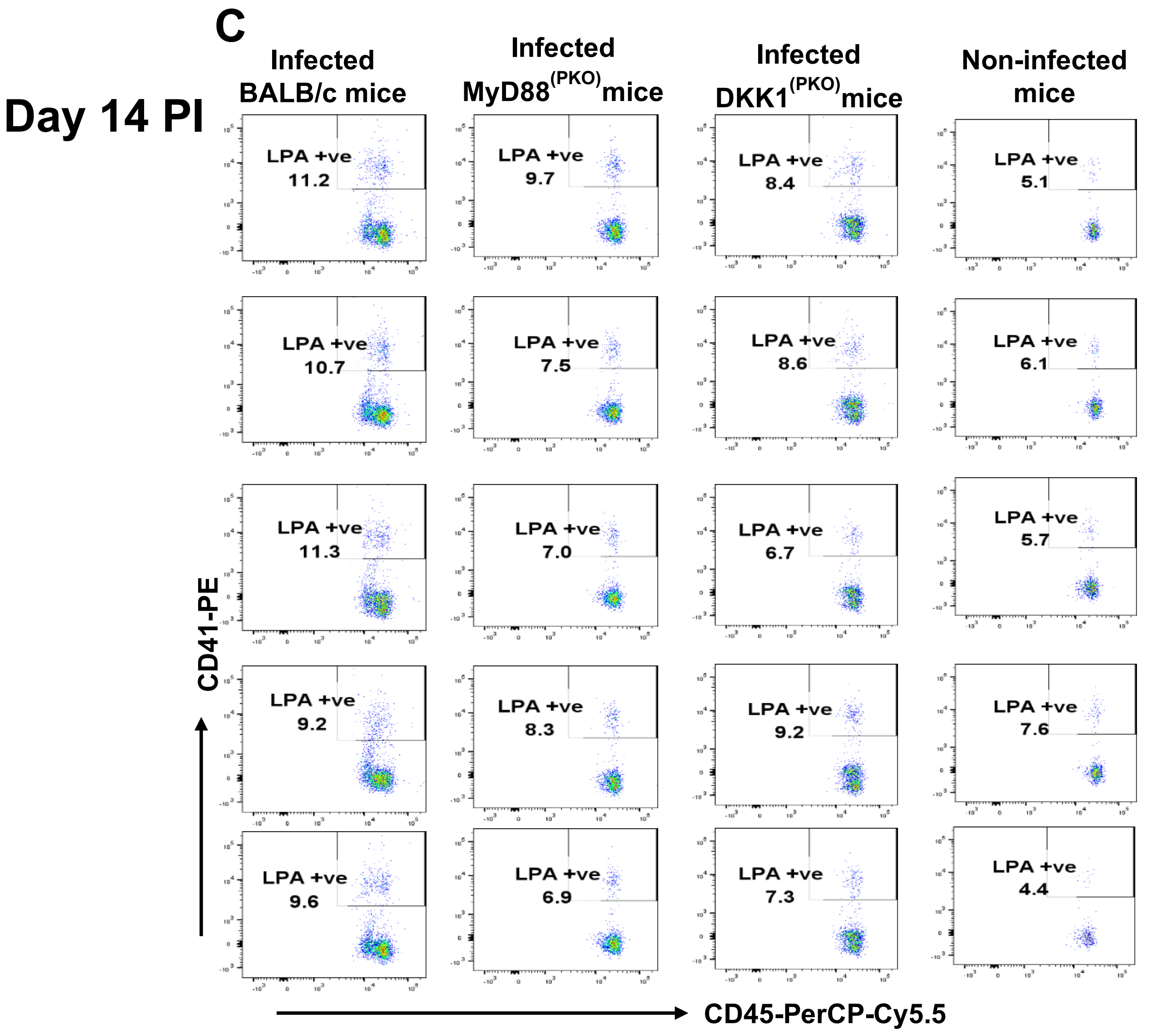
C
Day 14 PI
Infected BALB/c mice
Infected MyD88(PKO)mice
Infected DKK1(PKO)mice
Non-infected mice
LPA +ve 11.2
LPA +ve 9.7
LPA +ve 8.4
LPA +ve 5.1
LPA +ve 10.7
LPA +ve 7.5
LPA +ve 8.6
LPA +ve 6.1
LPA +ve 11.3
LPA +ve 7.0
LPA +ve 6.7
LPA +ve 5.7
LPA +ve 9.2
LPA +ve 8.3
LPA +ve 9.2
LPA +ve 7.6
LPA +ve 9.6
LPA +ve 6.9
LPA +ve 7.3
LPA +ve 4.4
CD41-PE
CD45-PerCP-Cy5.5

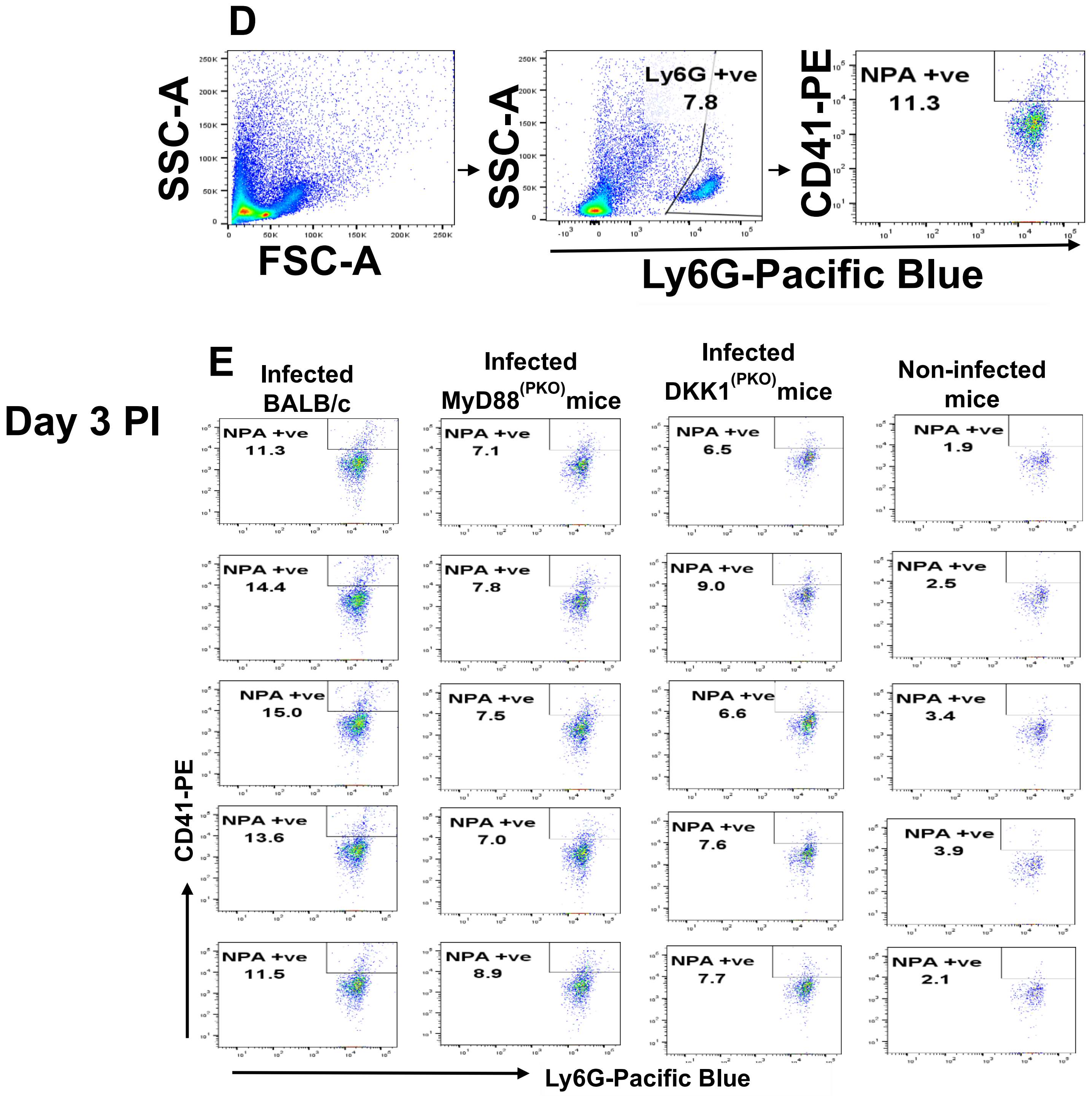
D
SSC-A
FSC-A
SSC-A
Ly6G +ve
7.8
CD41-PE
NPA +ve
11.3
Ly6G-Pacific Blue
E
Day 3 PI
Infected BALB/c
Infected MyD88(PKO)mice
Infected DKK1(PKO)mice
Non-infected mice
NPA +ve 11.3
NPA +ve 7.1
NPA +ve 6.5
NPA +ve 1.9
NPA +ve 14.4
NPA +ve 7.8
NPA +ve 9.0
NPA +ve 2.5
NPA +ve 15.0
NPA +ve 7.5
NPA +ve 6.6
NPA +ve 3.4
NPA +ve 13.6
NPA +ve 7.0
NPA +ve 7.6
NPA +ve 3.9
NPA +ve 11.5
NPA +ve 8.9
NPA +ve 7.7
NPA +ve 2.1
CD41-PE
Ly6G-Pacific Blue

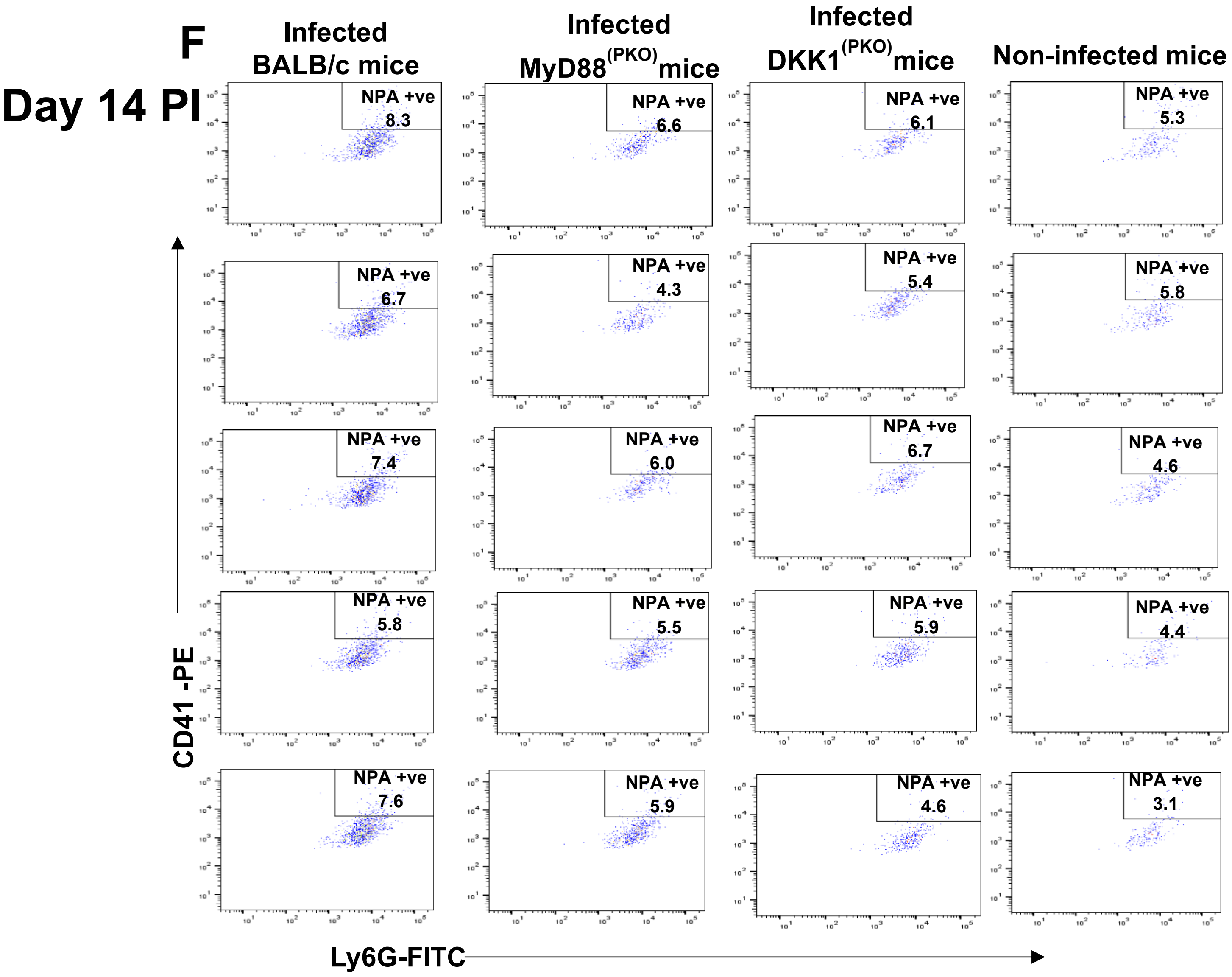


**Figure S1: Impaired LPA and NPA formation in infected MyD88$^{(PKO)}$ and DKK1$^{(PKO)}$ mice.** BALB/c, MyD88$^{(PKO)}$, and DKK1$^{(PKO)}$ mice were challenged with infective metacyclic promastigote (2 x $10^6$ parasites, n = 5) of WT *L. major* strain via the footpad. Control mice for LPA detection (n = 5) and NPA detection (n = 10/2 feet per mouse) were given 0.9% NaCl saline. Blood was collected via the maxillary vein on days 3 and 14 PI for the determination of LPA formation. Cells from the infected footpad were collected on days 3 and 14 PI for assessing NPA formation. Samples were analyzed by flow cytometry for LPA and NPA. Representative flow cytometry dot plots showing the analysis of LPA **(A)** and NPA **(D)** performed on day 3 PI. The dot plots shown in **(B, C, E, & F)** are from each sample in all the experimental groups obtained on day 3 and 14 PI, respectively. In all the experiments, BALB/c-infected and non-infected mice served as positive and negative controls, respectively.

A

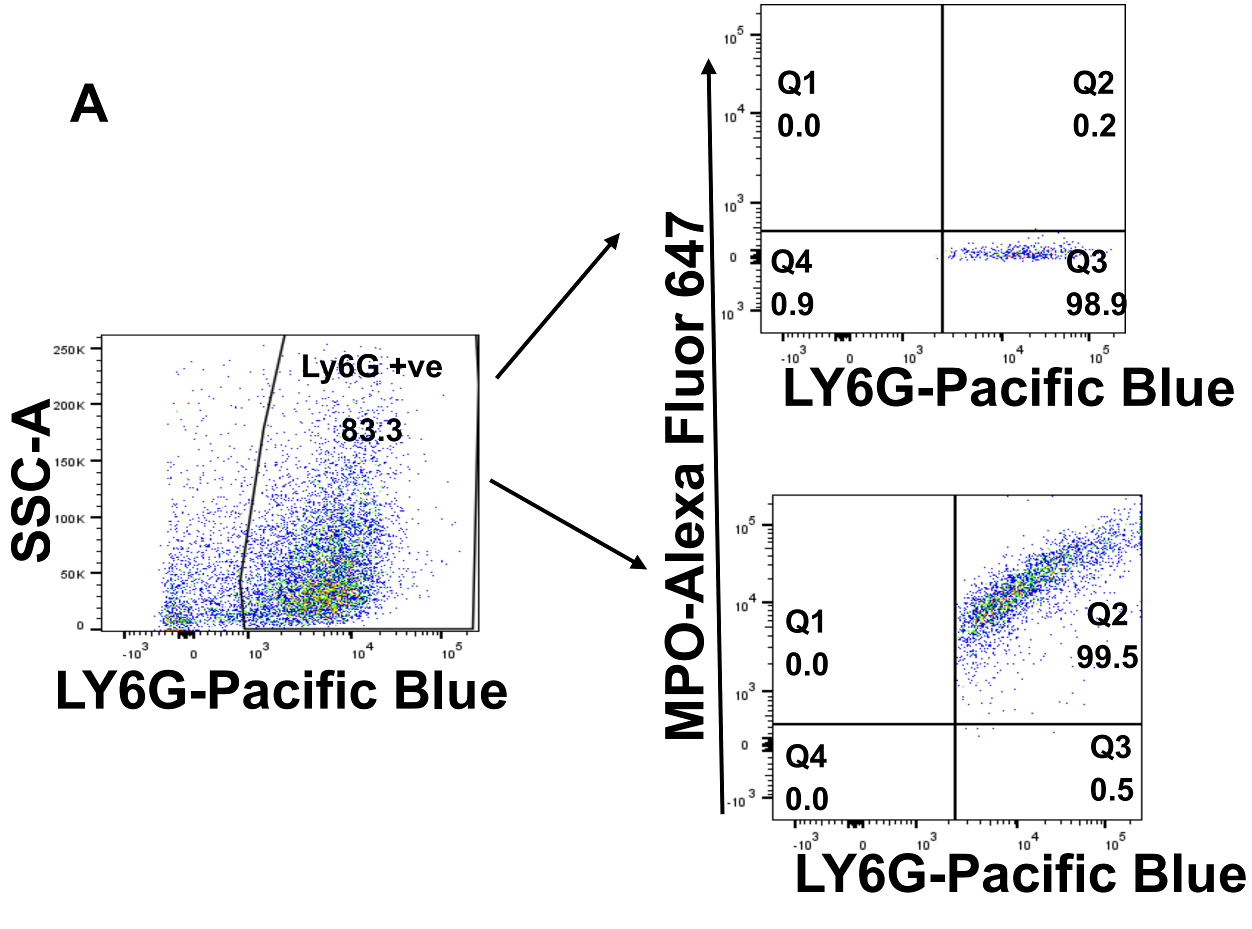
SSC-A
Ly6G +ve
83.3
LY6G-Pacific Blue
MPO-Alexa Fluor 647
Q1
0.0
Q2
0.2
Q4
0.9
Q3
98.9
LY6G-Pacific Blue
Q1
0.0
Q2
99.5
Q4
0.0
Q3
0.5
LY6G-Pacific Blue

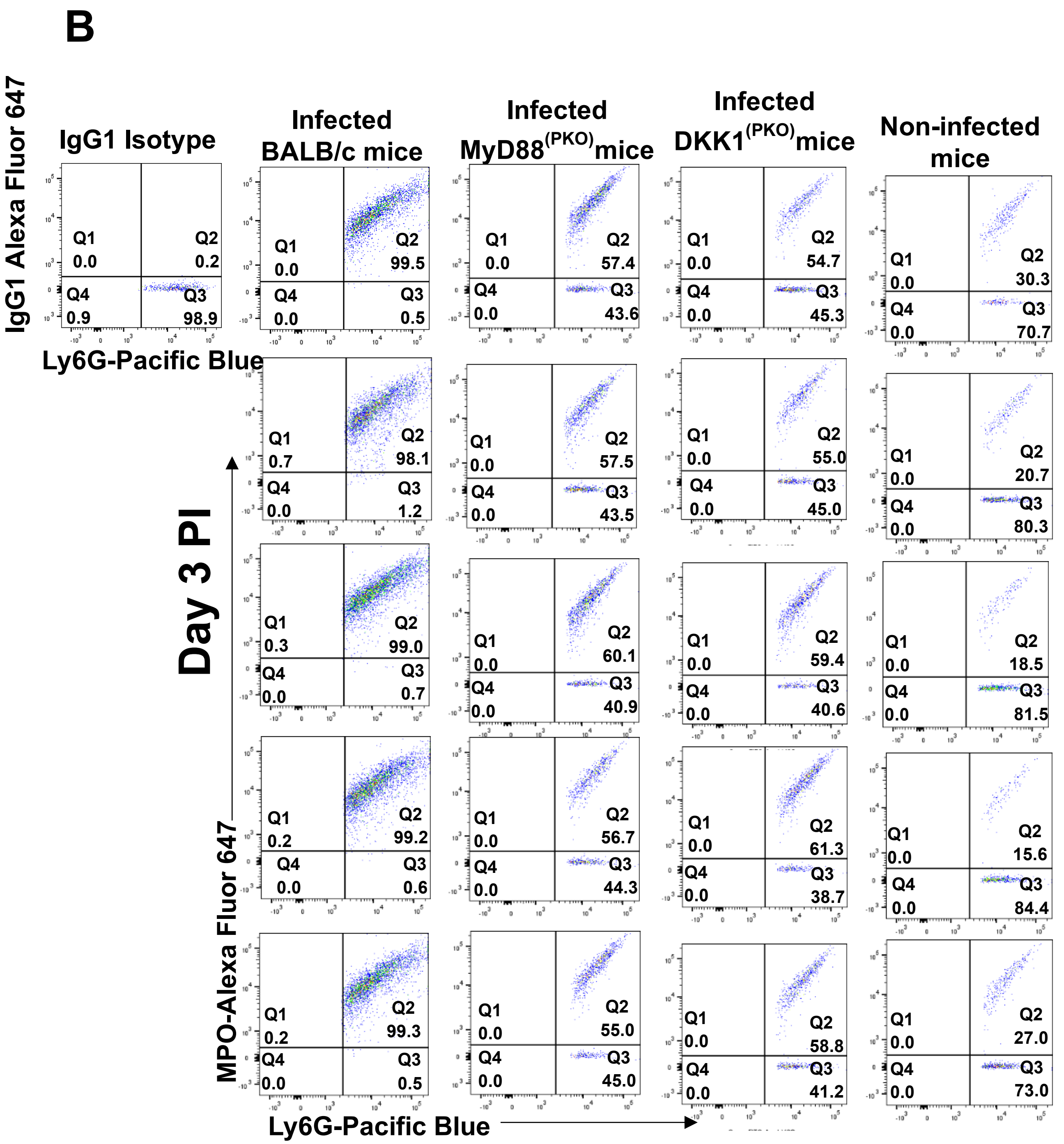
B
IgG1 Isotype
Infected BALB/c mice
Infected MyD88(PKO) mice
Infected DKK1(PKO) mice
Non-infected mice
IgG1 Alexa Fluor 647
Ly6G-Pacific Blue
Day 3 PI
MPO-Alexa Fluor 647
Ly6G-Pacific Blue
Q1 0.0 Q2 0.2 Q4 0.9 Q3 98.9
Q1 0.0 Q2 99.5 Q4 0.0 Q3 0.5
Q1 0.0 Q2 57.4 Q4 0.0 Q3 43.6
Q1 0.0 Q2 54.7 Q4 0.0 Q3 45.3
Q1 0.0 Q2 30.3 Q4 0.0 Q3 70.7
Q1 0.7 Q2 98.1 Q4 0.0 Q3 1.2
Q1 0.0 Q2 57.5 Q4 0.0 Q3 43.5
Q1 0.0 Q2 55.0 Q4 0.0 Q3 45.0
Q1 0.0 Q2 20.7 Q4 0.0 Q3 80.3
Q1 0.3 Q2 99.0 Q4 0.0 Q3 0.7
Q1 0.0 Q2 60.1 Q4 0.0 Q3 40.9
Q1 0.0 Q2 59.4 Q4 0.0 Q3 40.6
Q1 0.0 Q2 18.5 Q4 0.0 Q3 81.5
Q1 0.2 Q2 99.2 Q4 0.0 Q3 0.6
Q1 0.0 Q2 56.7 Q4 0.0 Q3 44.3
Q1 0.0 Q2 61.3 Q4 0.0 Q3 38.7
Q1 0.0 Q2 15.6 Q4 0.0 Q3 84.4
Q1 0.2 Q2 99.3 Q4 0.0 Q3 0.5
Q1 0.0 Q2 55.0 Q4 0.0 Q3 45.0
Q1 0.0 Q2 58.8 Q4 0.0 Q3 41.2
Q1 0.0 Q2 27.0 Q4 0.0 Q3 73.0

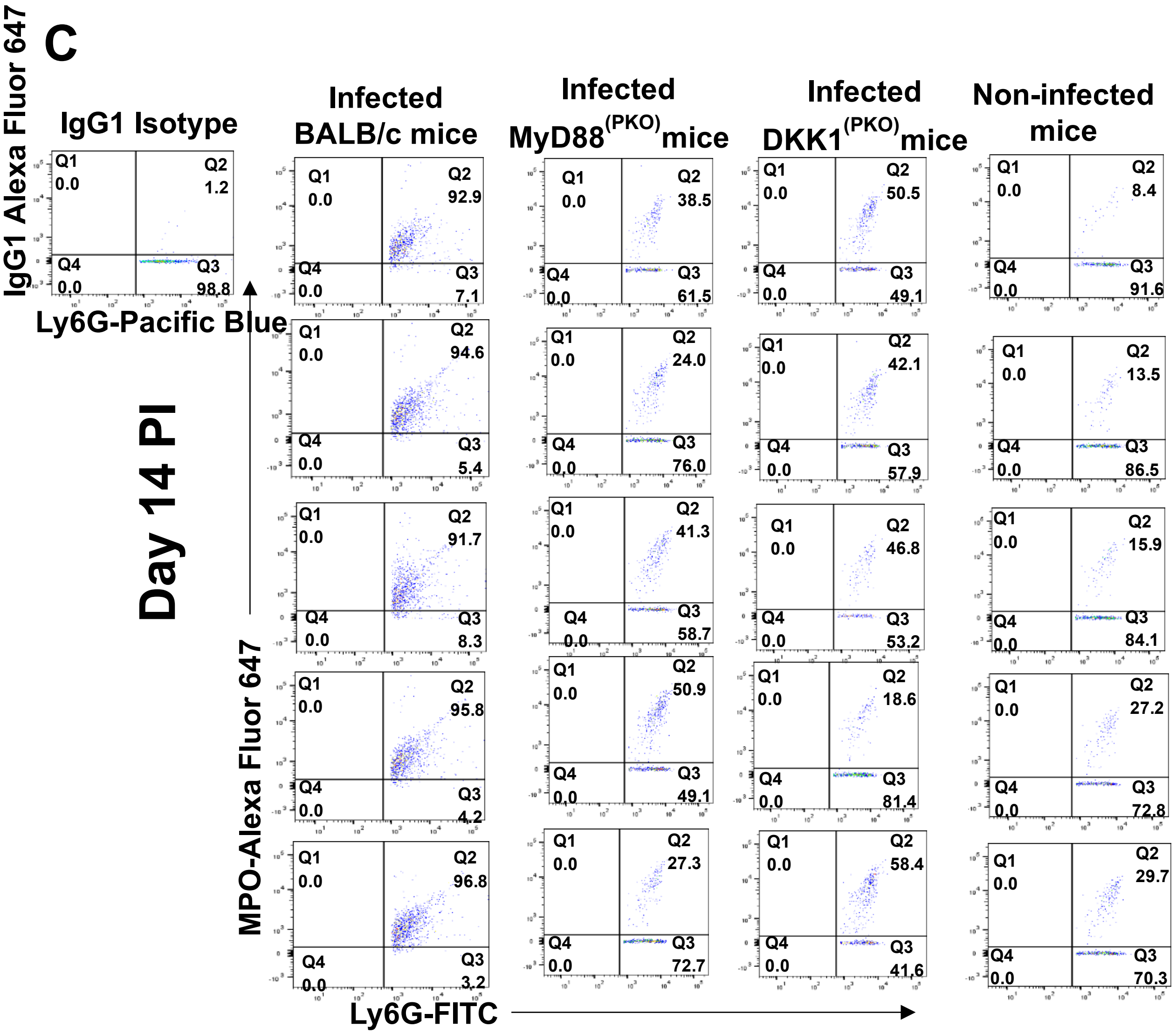

C
IgG1 Isotype
IgG1 Alexa Fluor 647
Ly6G-Pacific Blue
Q1 0.0 Q2 1.2 Q4 0.0 Q3 98.8
Infected BALB/c mice
Infected MyD88(PKO)mice
Infected DKK1(PKO)mice
Non-infected mice
Day 14 PI
MPO-Alexa Fluor 647
Ly6G-FITC
Q1 0.0 Q2 92.9 Q4 0.0 Q3 7.1
Q1 0.0 Q2 38.5 Q4 0.0 Q3 61.5
Q1 0.0 Q2 50.5 Q4 0.0 Q3 49.1
Q1 0.0 Q2 8.4 Q4 0.0 Q3 91.6
Q1 0.0 Q2 94.6 Q4 0.0 Q3 5.4
Q1 0.0 Q2 24.0 Q4 0.0 Q3 76.0
Q1 0.0 Q2 42.1 Q4 0.0 Q3 57.9
Q1 0.0 Q2 13.5 Q4 0.0 Q3 86.5
Q1 0.0 Q2 91.7 Q4 0.0 Q3 8.3
Q1 0.0 Q2 41.3 Q4 0.0 Q3 58.7
Q1 0.0 Q2 46.8 Q4 0.0 Q3 53.2
Q1 0.0 Q2 15.9 Q4 0.0 Q3 84.1
Q1 0.0 Q2 95.8 Q4 0.0 Q3 4.2
Q1 0.0 Q2 50.9 Q4 0.0 Q3 49.1
Q1 0.0 Q2 18.6 Q4 0.0 Q3 81.4
Q1 0.0 Q2 27.2 Q4 0.0 Q3 72.8
Q1 0.0 Q2 96.8 Q4 0.0 Q3 3.2
Q1 0.0 Q2 27.3 Q4 0.0 Q3 72.7
Q1 0.0 Q2 58.4 Q4 0.0 Q3 41.6
Q1 0.0 Q2 29.7 Q4 0.0 Q3 70.3

D

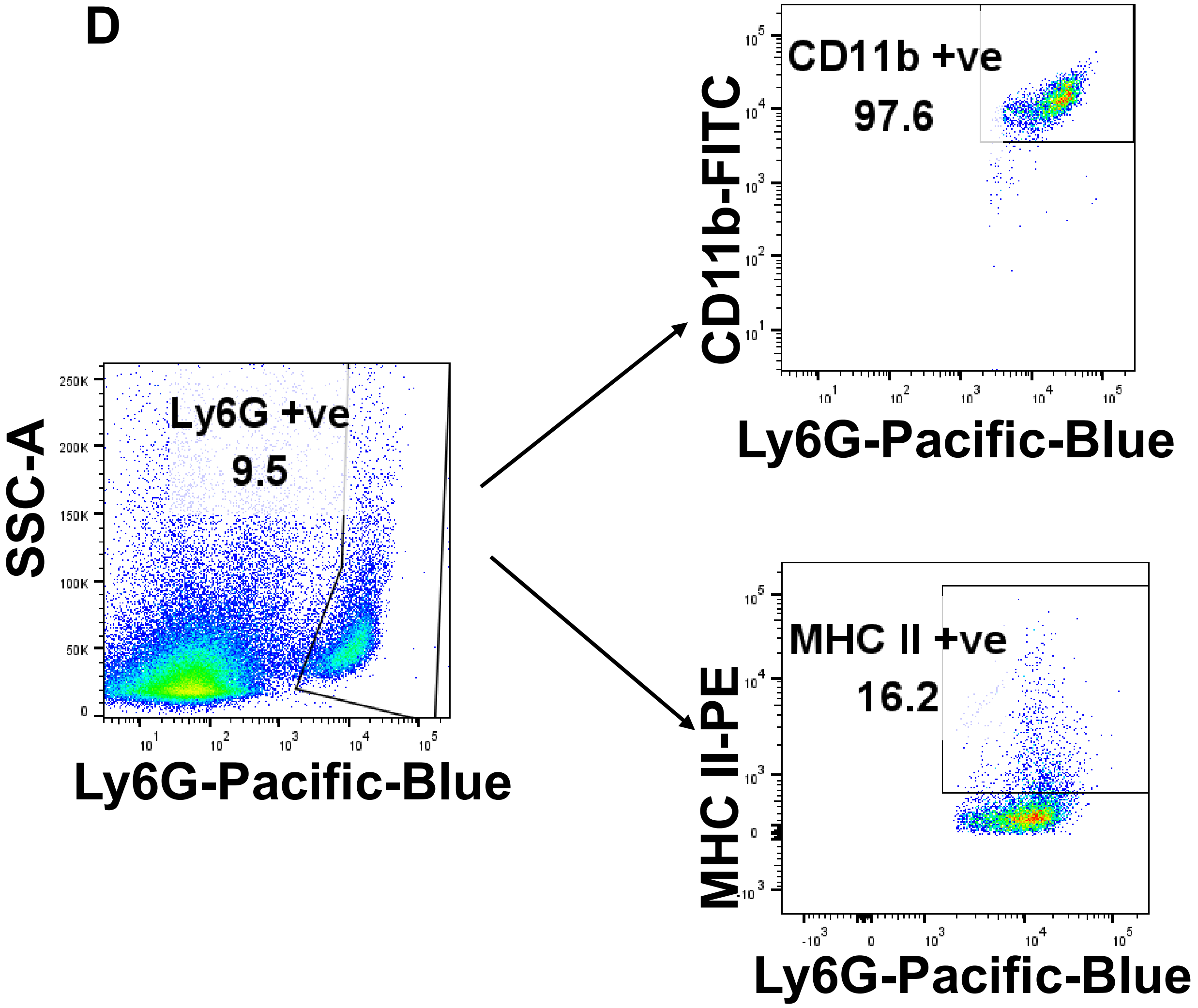
SSC-A
Ly6G +ve
9.5
Ly6G-Pacific-Blue
CD11b-FITC
CD11b +ve
97.6
Ly6G-Pacific-Blue
MHC II-PE
MHC II +ve
16.2
Ly6G-Pacific-Blue

E

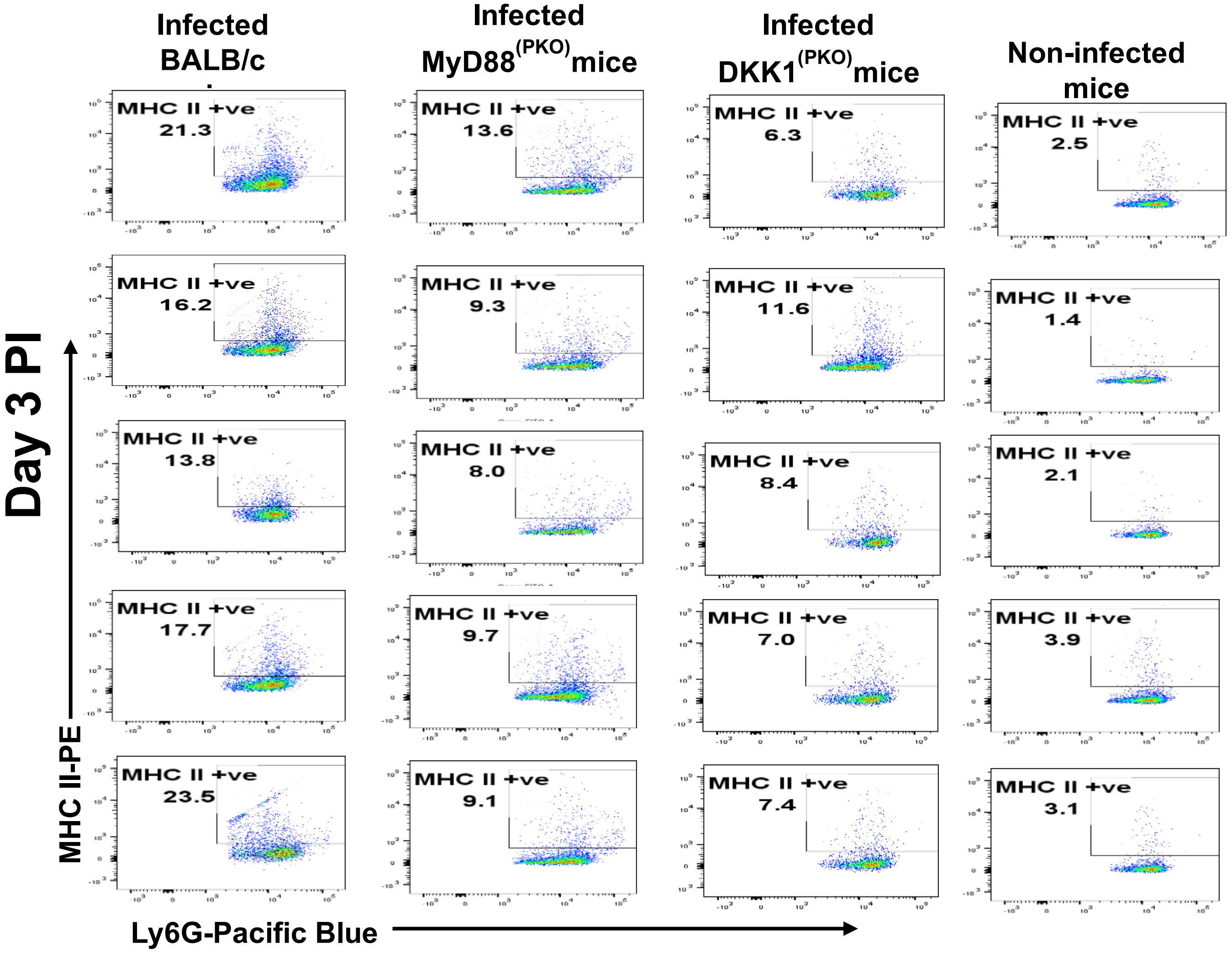
Infected BALB/c
Infected MyD88(PKO)mice
Infected DKK1(PKO)mice
Non-infected mice
Day 3 PI
MHC II-PE
Ly6G-Pacific Blue
MHC II +ve
21.3
13.6
6.3
2.5
16.2
9.3
11.6
1.4
13.8
8.0
8.4
2.1
17.7
9.7
7.0
3.9
23.5
9.1
7.4
3.1

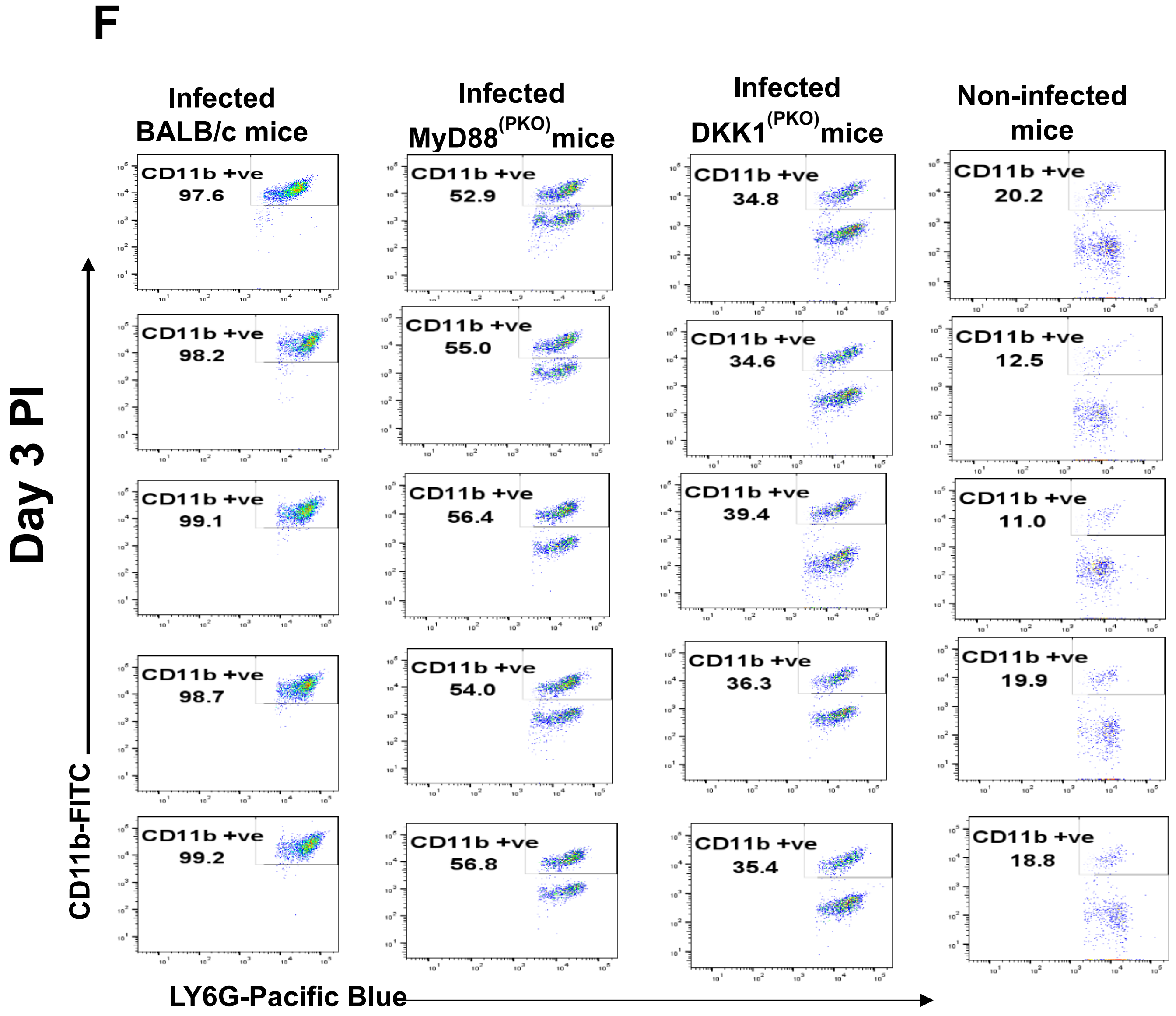
F
Infected BALB/c mice
Infected MyD88(PKO) mice
Infected DKK1(PKO) mice
Non-infected mice
Day 3 PI
CD11b-FITC
LY6G-Pacific Blue
CD11b +ve 97.6
CD11b +ve 98.2
CD11b +ve 99.1
CD11b +ve 98.7
CD11b +ve 99.2
CD11b +ve 52.9
CD11b +ve 55.0
CD11b +ve 56.4
CD11b +ve 54.0
CD11b +ve 56.8
CD11b +ve 34.8
CD11b +ve 34.6
CD11b +ve 39.4
CD11b +ve 36.3
CD11b +ve 35.4
CD11b +ve 20.2
CD11b +ve 12.5
CD11b +ve 11.0
CD11b +ve 19.9
CD11b +ve 18.8

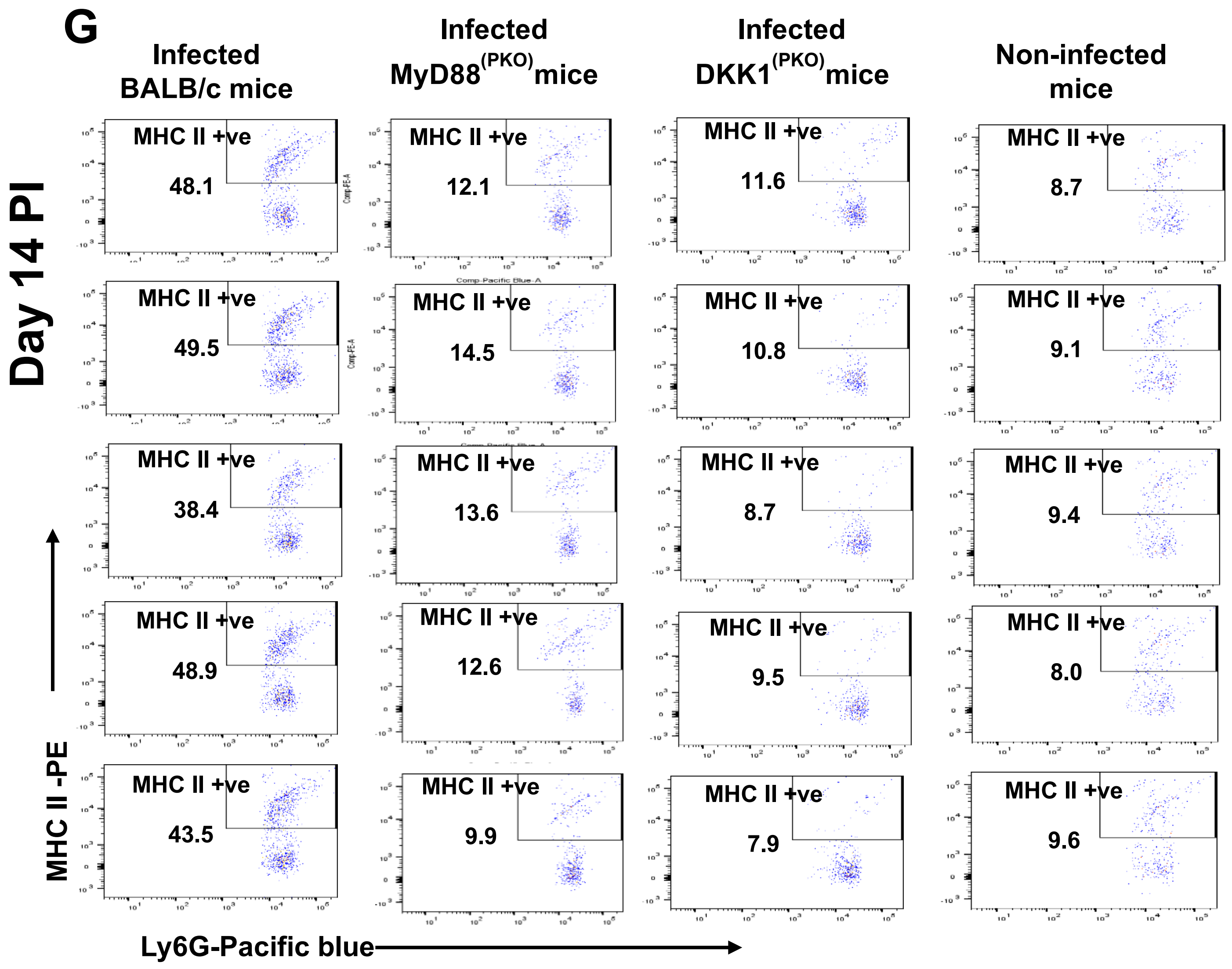
G
Infected BALB/c mice
Infected MyD88(PKO)mice
Infected DKK1(PKO)mice
Non-infected mice
Day 14 PI
MHC II +ve
48.1
12.1
11.6
8.7
49.5
14.5
10.8
9.1
38.4
13.6
8.7
9.4
48.9
12.6
9.5
8.0
43.5
9.9
7.9
9.6
MHC II -PE
Ly6G-Pacific blue

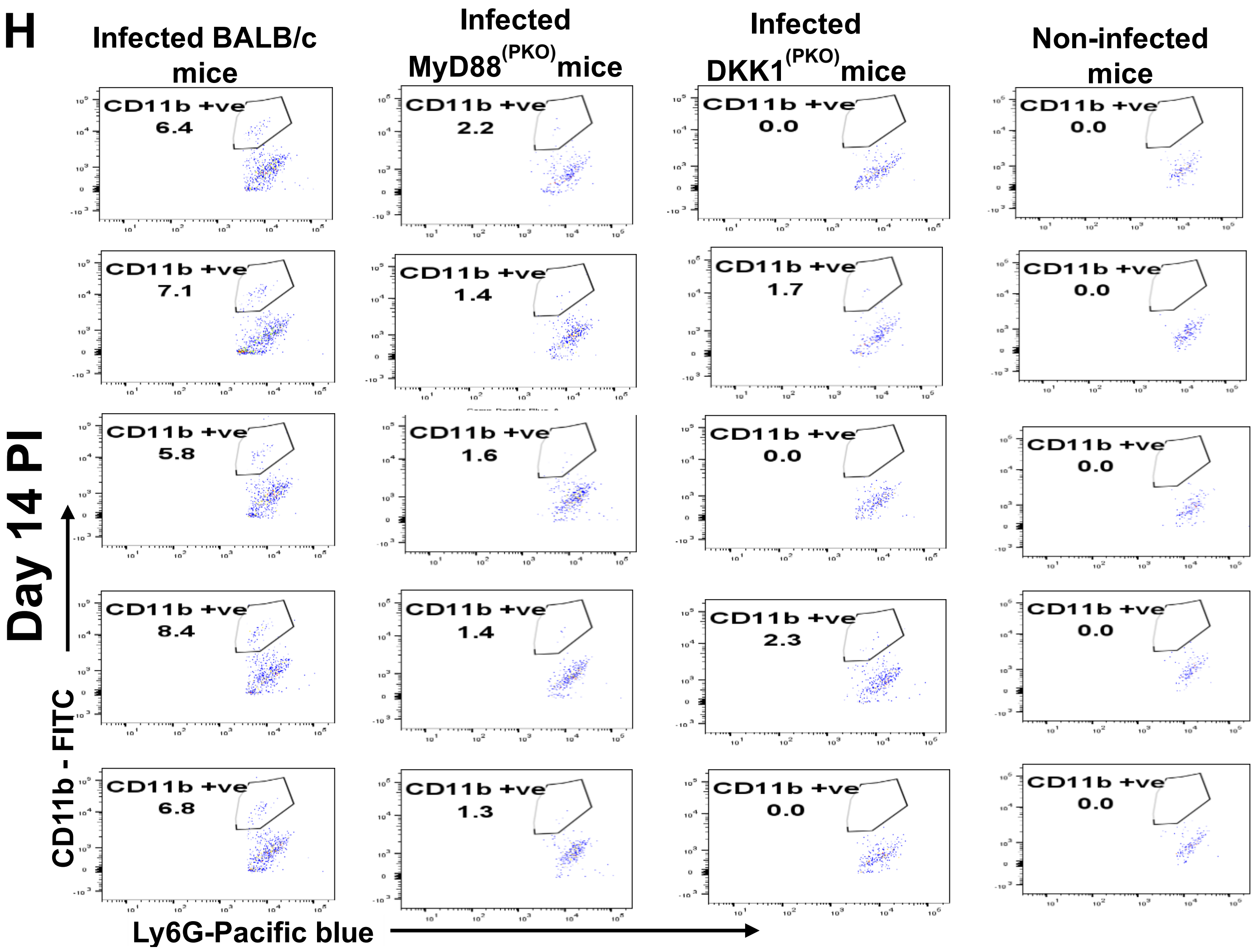


**Figure S2: Decreased MPO$^+$, CD11b$^+$ and MHC class II$^+$ neutrophils in MyD88$^{(PKO)}$ and DKK1$^{(PKO)}$-infected mice.** The BALB/c, MyD88$^{(PKO)}$ and DKK$^{(PKO)}$ mice were challenged with infective metacyclic promastigote (2 x $10^6$ parasites, n = 5) of *L. major* via the footpad. Non-infected BALB/c mice (n = 10/2 feet per mouse) were given 0.9% NaCl saline. Neutrophils were isolated from the footpads of all infected and non-infected mice at day 3 and 14 PI. Neutrophil samples were analyzed by flow cytometry for MPO$^+$CD11b$^+$ MHC class II+ neutrophils. Representative flow cytometry dot plots showing the analyses of MPO$^+$ neutrophils and IgG1 isotype control for non-specific antibody staining on day 3 PI **(A)**. In addition, a representative flow cytometry dot plot showing the analyses of CD11b$^+$ and MHC class II$^+$ neutrophils performed on day 3 PI **(D)** is indicated. A dot plot of each sample in all the experimental groups obtained on days 3 and 14 PI is presented in **(B, C, E, F, G, & H),** respectively. In all experiments, BALB/c mice infected and non-infected served as positive and negative controls, respectively.

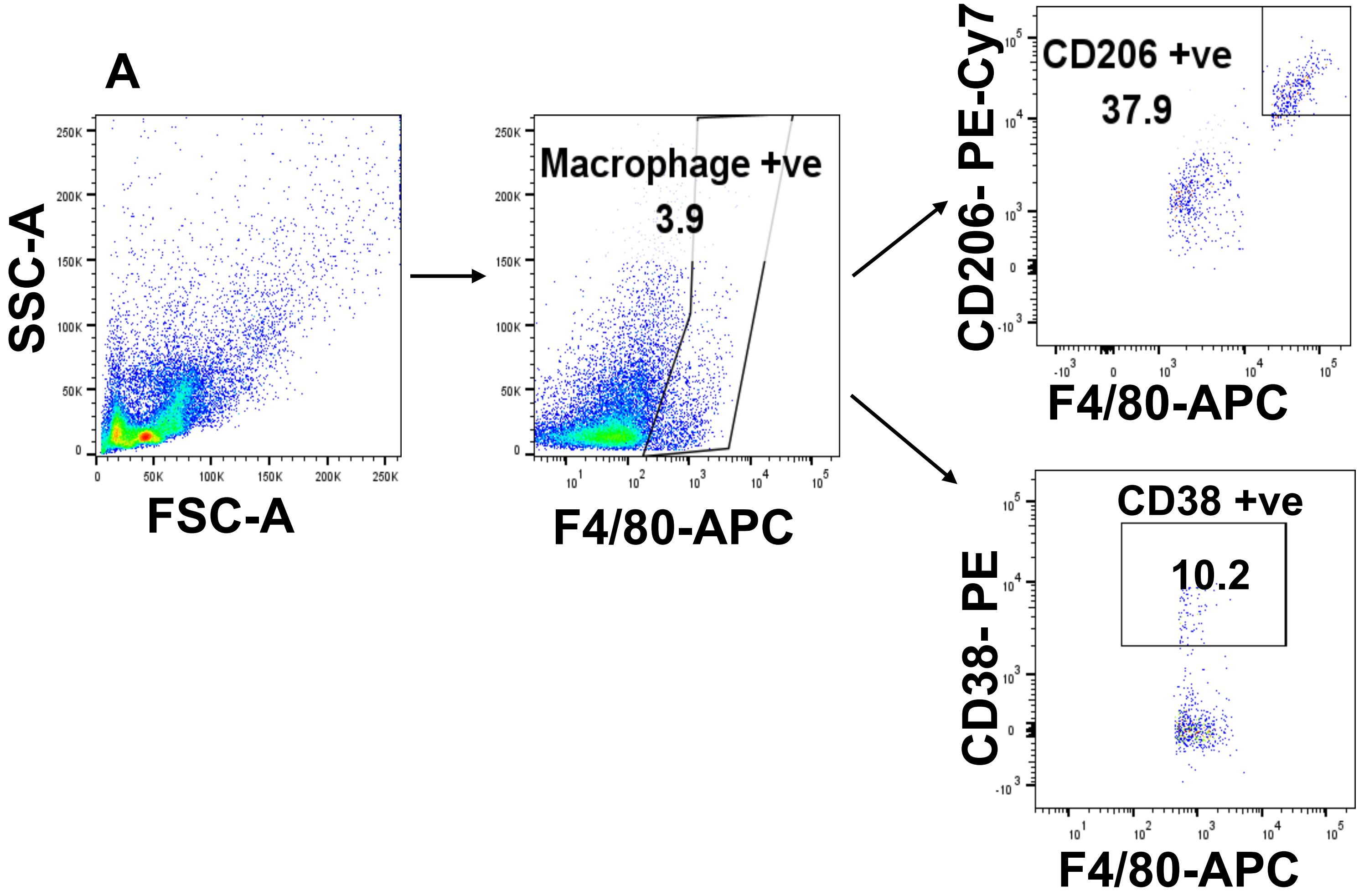
A
SSC-A
FSC-A
Macrophage +ve
3.9
F4/80-APC
CD206- PE-Cy7
CD206 +ve
37.9
F4/80-APC
CD38- PE
CD38 +ve
10.2
F4/80-APC

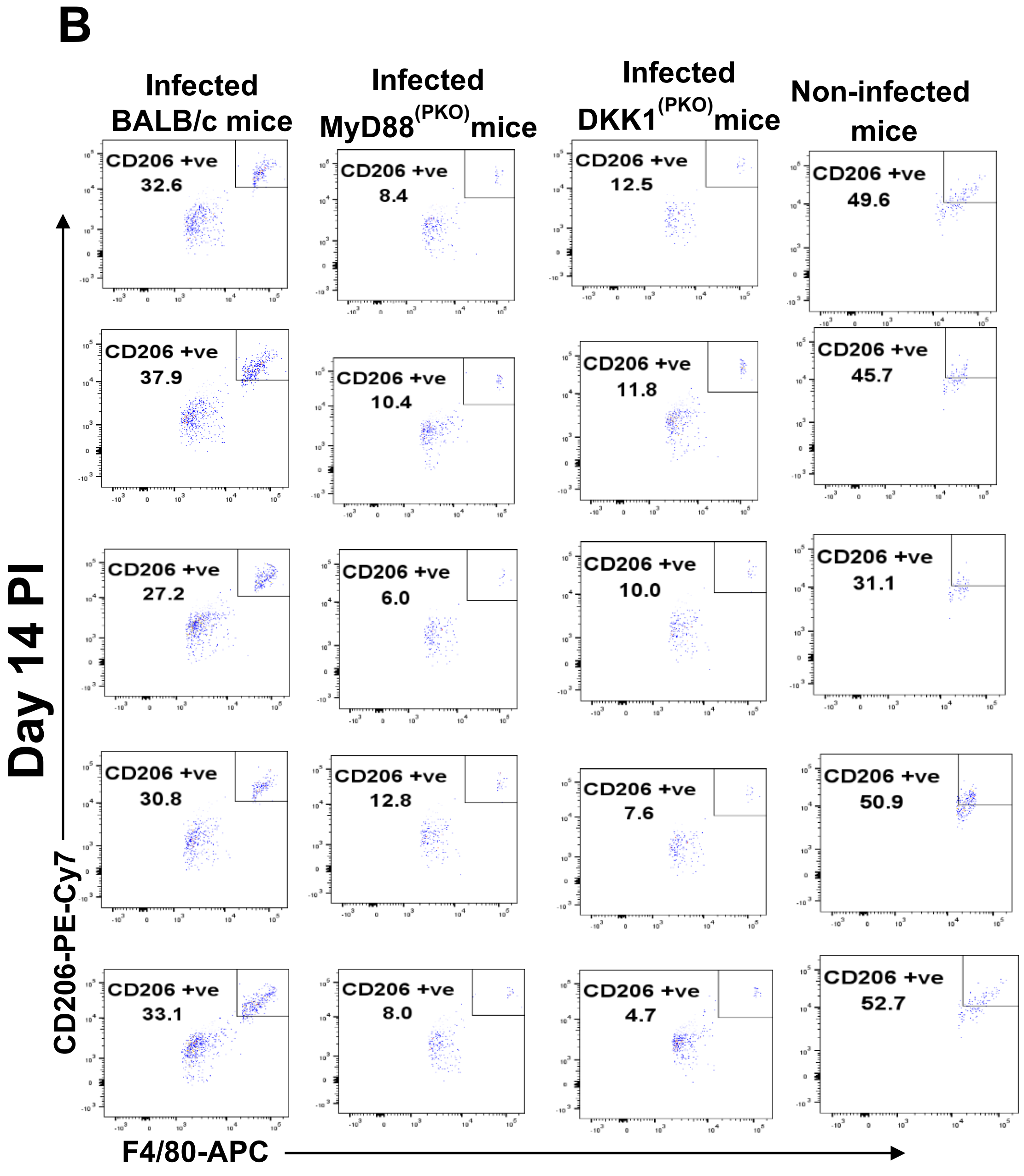
B
Infected BALB/c mice
Infected MyD88(PKO)mice
Infected DKK1(PKO)mice
Non-infected mice
Day 14 PI
CD206 +ve 32.6
CD206 +ve 8.4
CD206 +ve 12.5
CD206 +ve 49.6
CD206 +ve 37.9
CD206 +ve 10.4
CD206 +ve 11.8
CD206 +ve 45.7
CD206 +ve 27.2
CD206 +ve 6.0
CD206 +ve 10.0
CD206 +ve 31.1
CD206 +ve 30.8
CD206 +ve 12.8
CD206 +ve 7.6
CD206 +ve 50.9
CD206 +ve 33.1
CD206 +ve 8.0
CD206 +ve 4.7
CD206 +ve 52.7
CD206-PE-Cy7
F4/80-APC

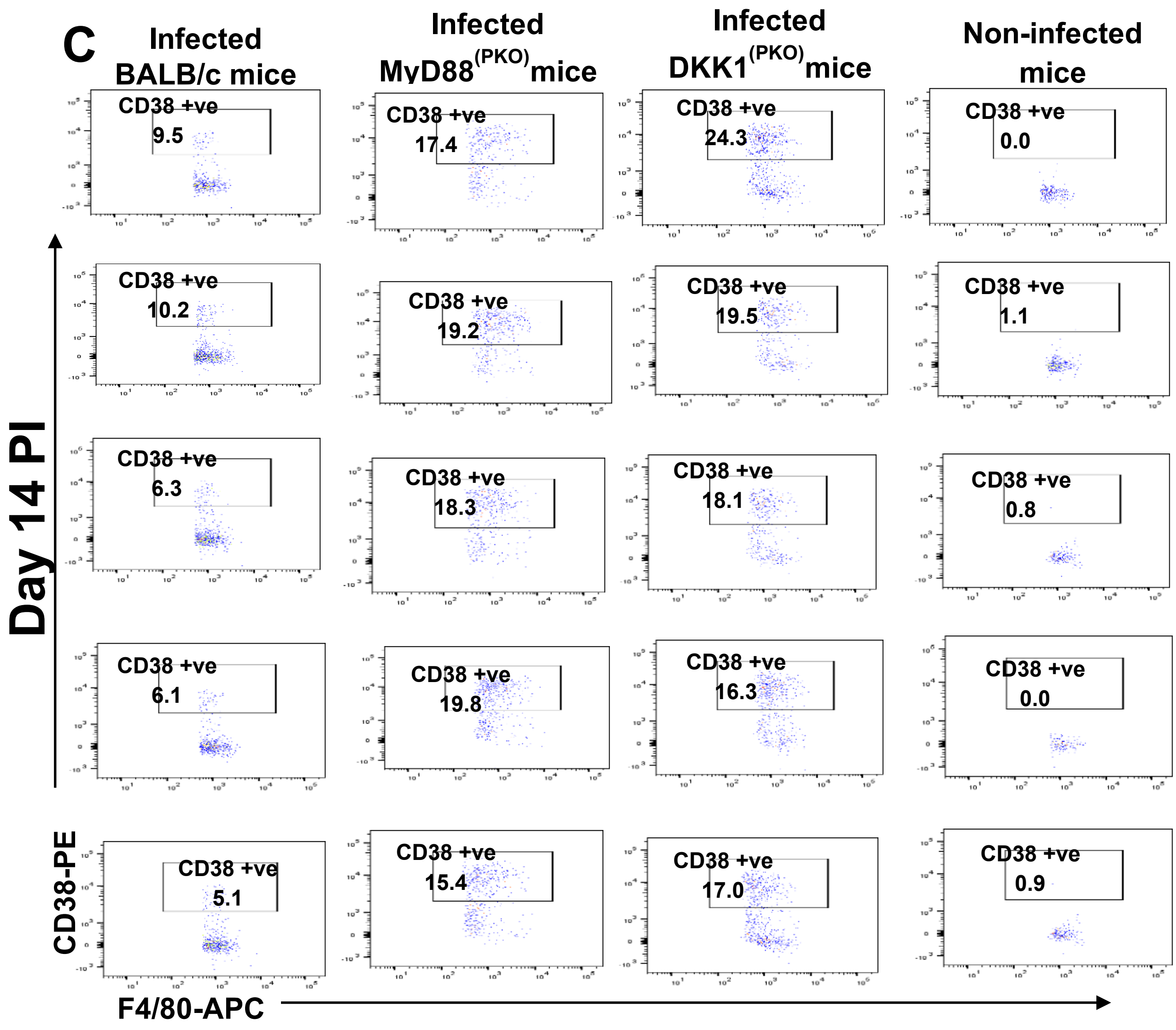
C
Infected BALB/c mice
Infected MyD88(PKO)mice
Infected DKK1(PKO)mice
Non-infected mice
Day 14 PI
CD38 +ve
9.5
17.4
24.3
0.0
10.2
19.2
19.5
1.1
6.3
18.3
18.1
0.8
6.1
19.8
16.3
0.0
5.1
15.4
17.0
0.9
CD38-PE
F4/80-APC

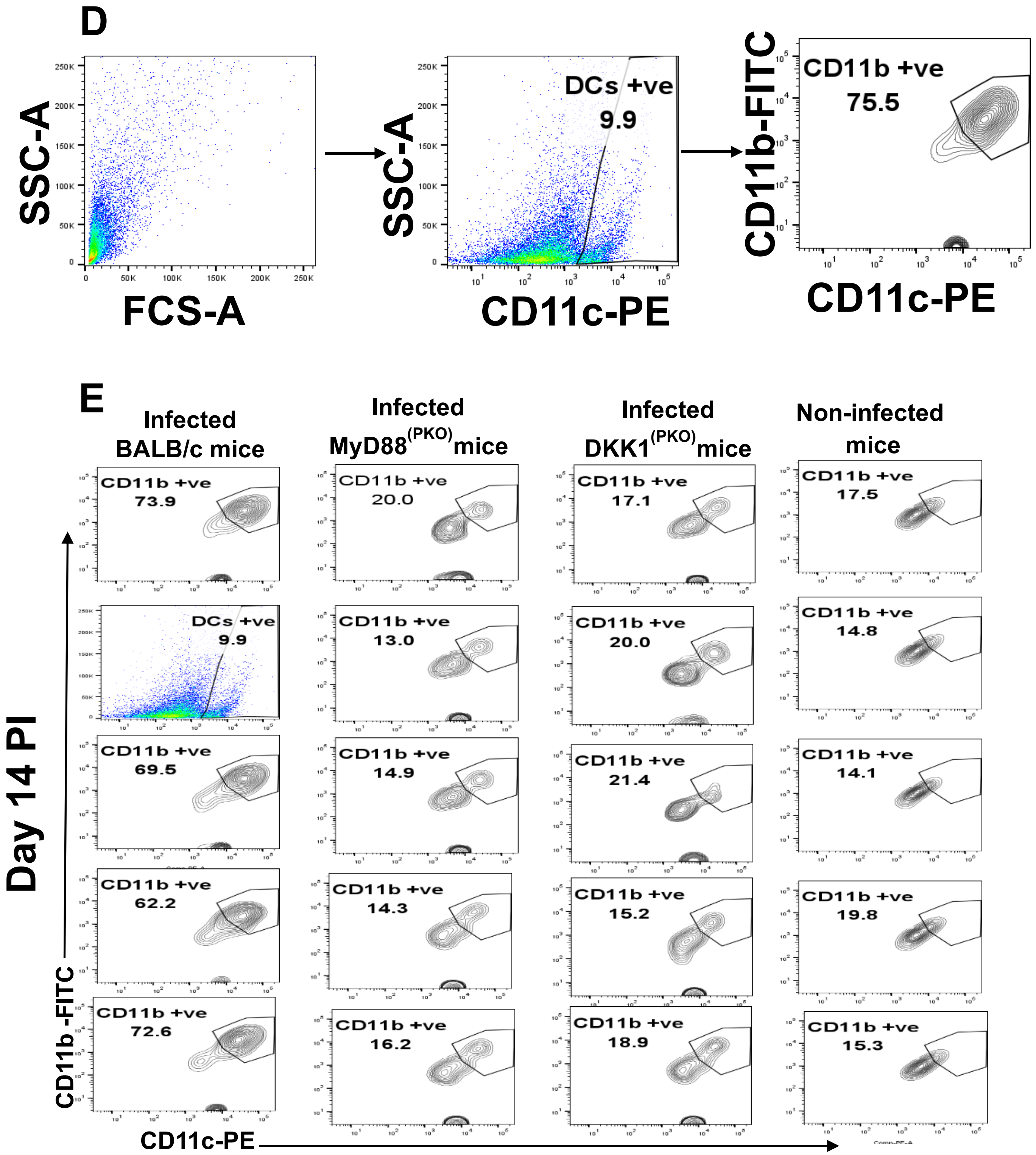
D
SSC-A
FCS-A
SSC-A
CD11c-PE
DCs +ve
9.9
CD11b-FITC
CD11c-PE
CD11b +ve
75.5
E
Infected BALB/c mice
Infected MyD88(PKO)mice
Infected DKK1(PKO)mice
Non-infected mice
Day 14 PI
CD11b -FITC
CD11c-PE
CD11b +ve 73.9
DCs +ve 9.9
CD11b +ve 69.5
CD11b +ve 62.2
CD11b +ve 72.6
CD11b +ve 20.0
CD11b +ve 13.0
CD11b +ve 14.9
CD11b +ve 14.3
CD11b +ve 16.2
CD11b +ve 17.1
CD11b +ve 20.0
CD11b +ve 21.4
CD11b +ve 15.2
CD11b +ve 18.9
CD11b +ve 17.5
CD11b +ve 14.8
CD11b +ve 14.1
CD11b +ve 19.8
CD11b +ve 15.3

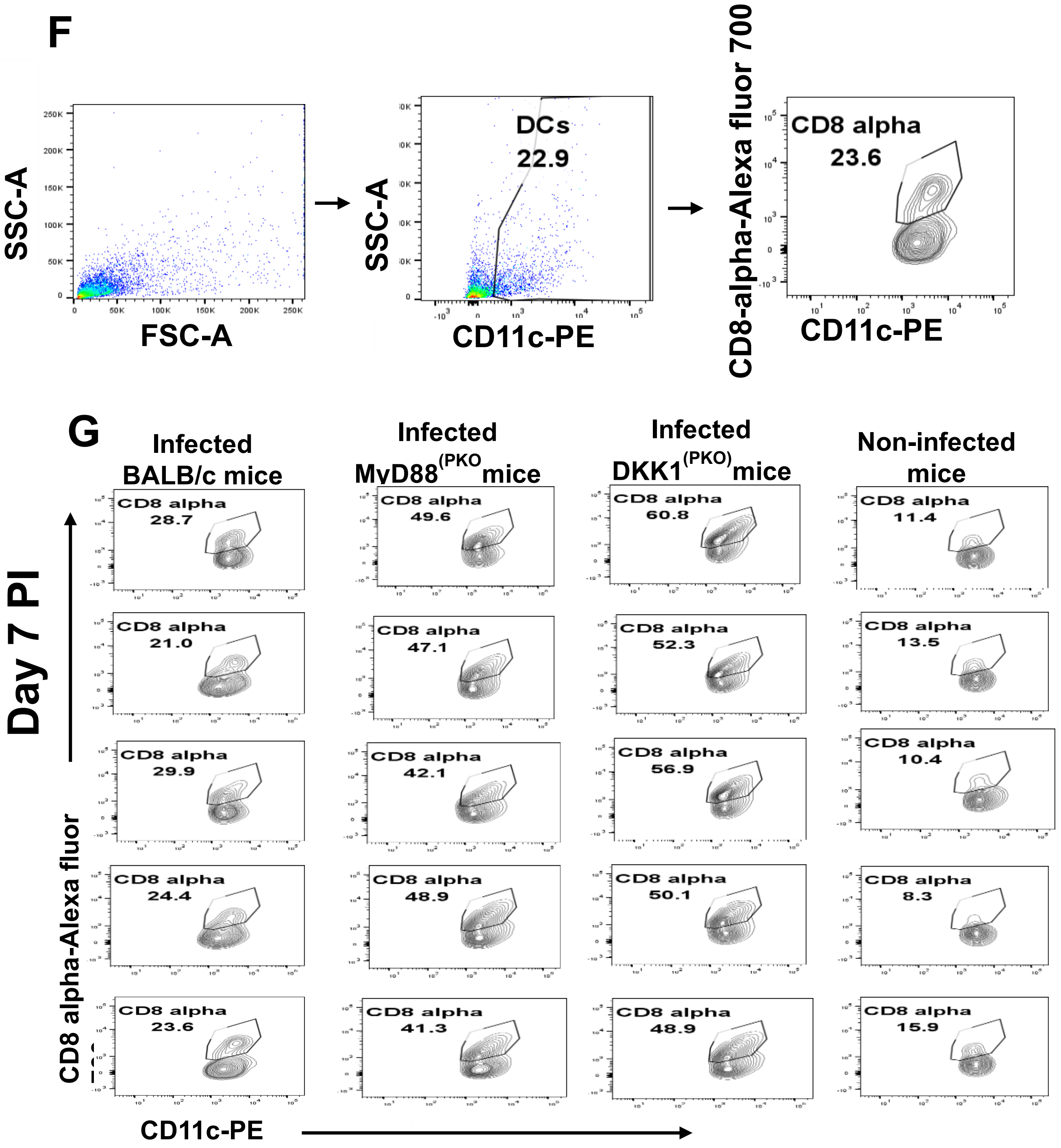


**Figure S3: Increased $CD38^+$ macrophages and $CD8\alpha^+$dendritic cells in $MyD88^{(PKO)}$ and $DKK1^{(PKO)}$ infected mice.** BALB/c, $MyD88^{(PKO)}$ and $DKK1^{(PKO)}$ mice were challenged with infective metacyclic promastigote (2 x $10^6$ parasites, n = 5) of *L. major* via the footpad. Non-infected BALB/c mice (n = 10/2 feet per mouse) were given 0.9% NaCl saline. Macrophages and dendritic cells were isolated from the footpads of all infected and non-infected mice on day 14 PI. Isolated macrophage and dendritic cell samples were analyzed by flow cytometry for M1($CD38^+$) and M2($CD206^+$) macrophages, as well as cDC1($CD8\alpha^+$) and cDC2 ($CD11b^+$) dendritic cells. Representative flow cytometry dot plots showing the analysis of $CD38^+$ and $CD206^+$ macrophages **(A)**, as well as $CD11b^+$ and $CD8\alpha^+$dendritic cells performed on day 14PI **(D) and (F).** A dot plot of each macrophage **(B & C)** and dendritic cell **(E & G)** sample in all the experimental groups is indicated**.** WT-infected and non-infected mice served as positive and negative controls in all experiments, respectively.

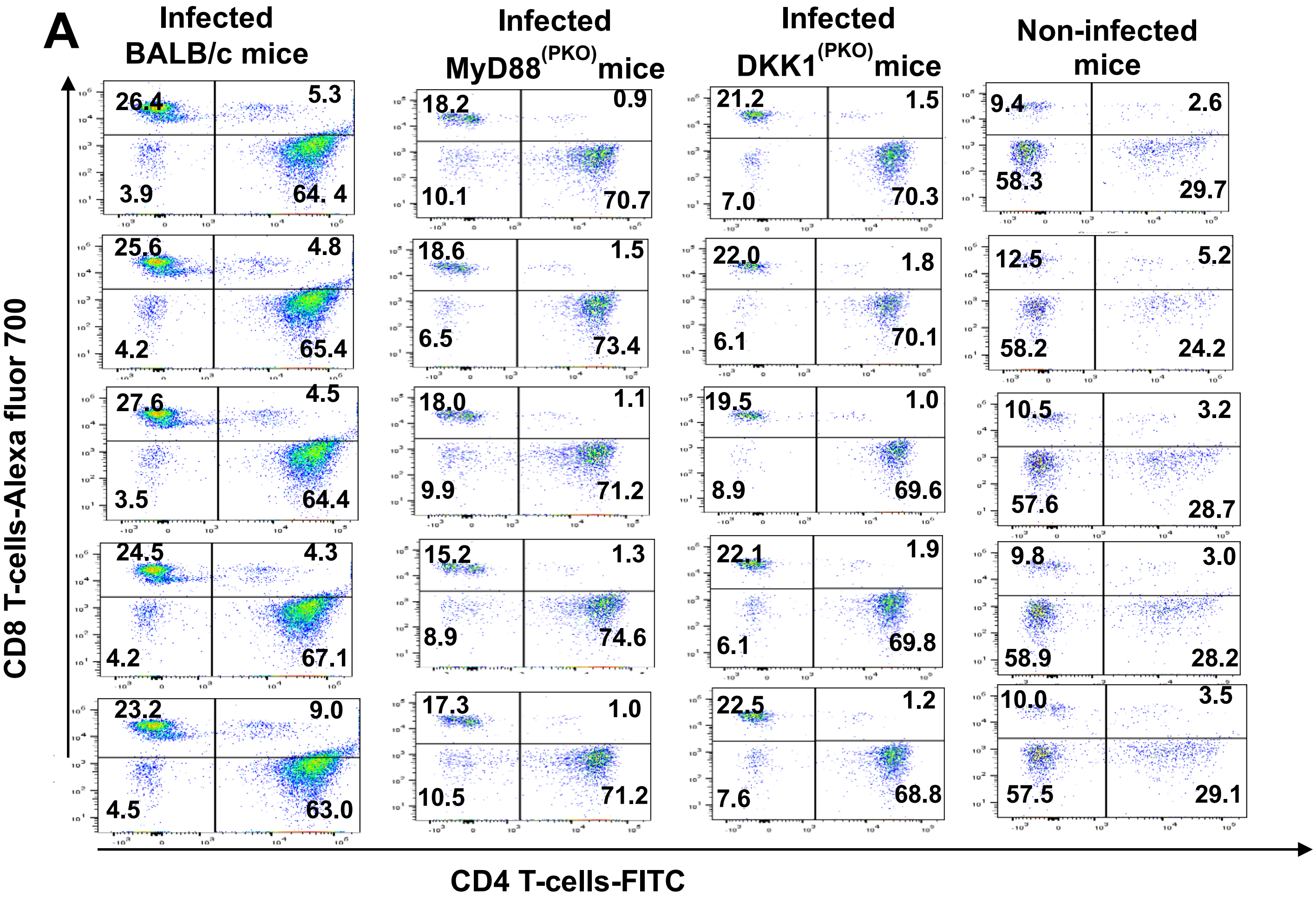
A
Infected BALB/c mice
Infected MyD88(PKO) mice
Infected DKK1(PKO) mice
Non-infected mice
CD8 T-cells-Alexa fluor 700
CD4 T-cells-FITC
26.4
5.3
3.9
64. 4
18.2
0.9
10.1
70.7
21.2
1.5
7.0
70.3
9.4
2.6
58.3
29.7
25.6
4.8
4.2
65.4
18.6
1.5
6.5
73.4
22.0
1.8
6.1
70.1
12.5
5.2
58.2
24.2
27.6
4.5
3.5
64.4
18.0
1.1
9.9
71.2
19.5
1.0
8.9
69.6
10.5
3.2
57.6
28.7
24.5
4.3
4.2
67.1
15.2
1.3
8.9
74.6
22.1
1.9
6.1
69.8
9.8
3.0
58.9
28.2
23.2
9.0
4.5
63.0
17.3
1.0
10.5
71.2
22.5
1.2
7.6
68.8
10.0
3.5
57.5
29.1

**B**

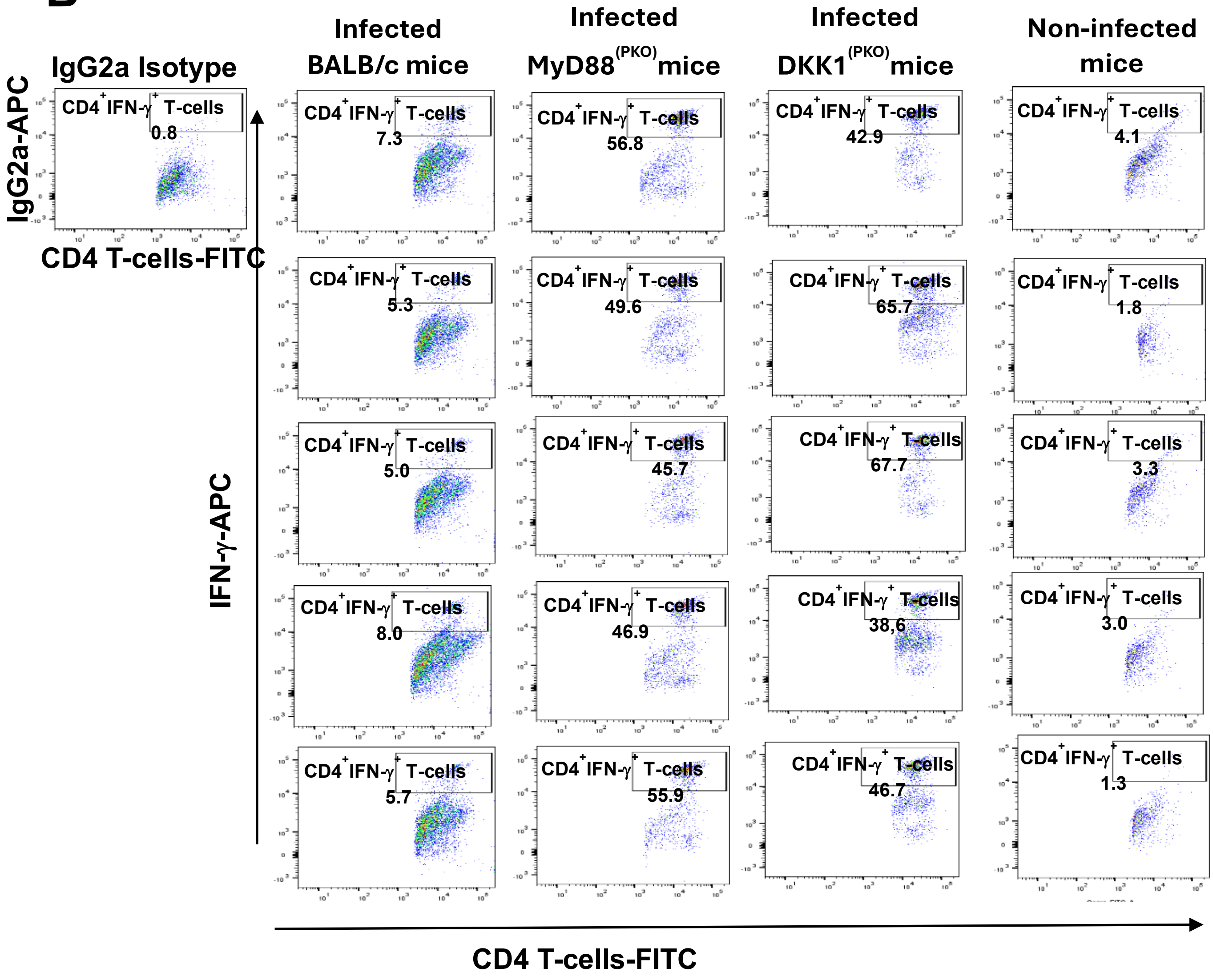
IgG2a Isotype
Infected BALB/c mice
Infected MyD88(PKO) mice
Infected DKK1(PKO) mice
Non-infected mice
IgG2a-APC
CD4 T-cells-FITC
IFN-γ-APC
CD4+IFN-γ+ T-cells
0.8
7.3
5.3
5.0
8.0
5.7
56.8
49.6
45.7
46.9
55.9
42.9
65.7
67.7
38,6
46.7
4.1
1.8
3.3
3.0
1.3
CD4 T-cells-FITC

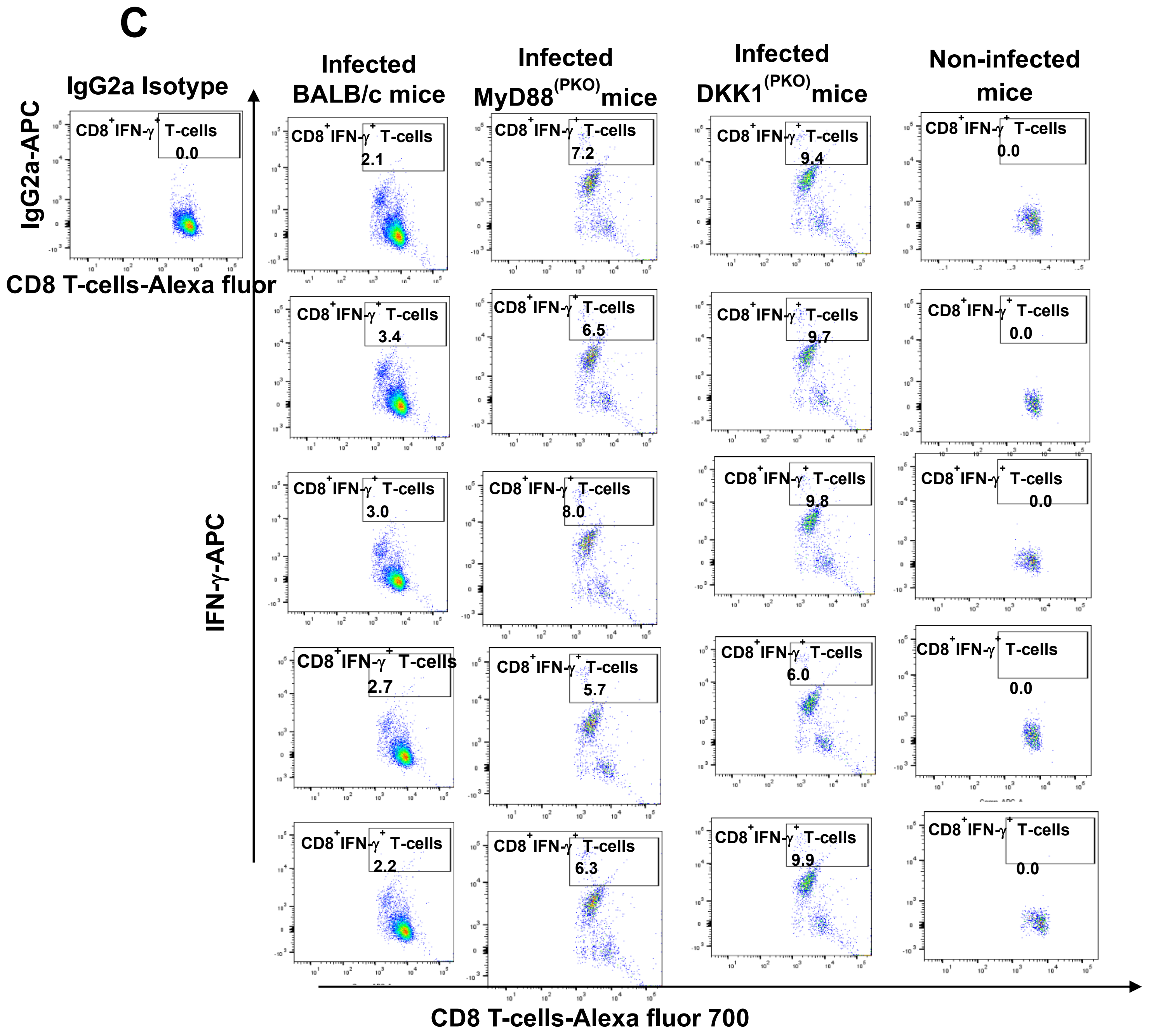
C
IgG2a Isotype
Infected BALB/c mice
Infected MyD88(PKO) mice
Infected DKK1(PKO) mice
Non-infected mice
IgG2a-APC
CD8 T-cells-Alexa fluor
IFN-γ-APC
CD8 T-cells-Alexa fluor 700
CD8+IFN-γ+ T-cells
0.0
2.1
3.4
3.0
2.7
2.2
7.2
6.5
8.0
5.7
6.3
9.4
9.7
9.8
6.0
9.9

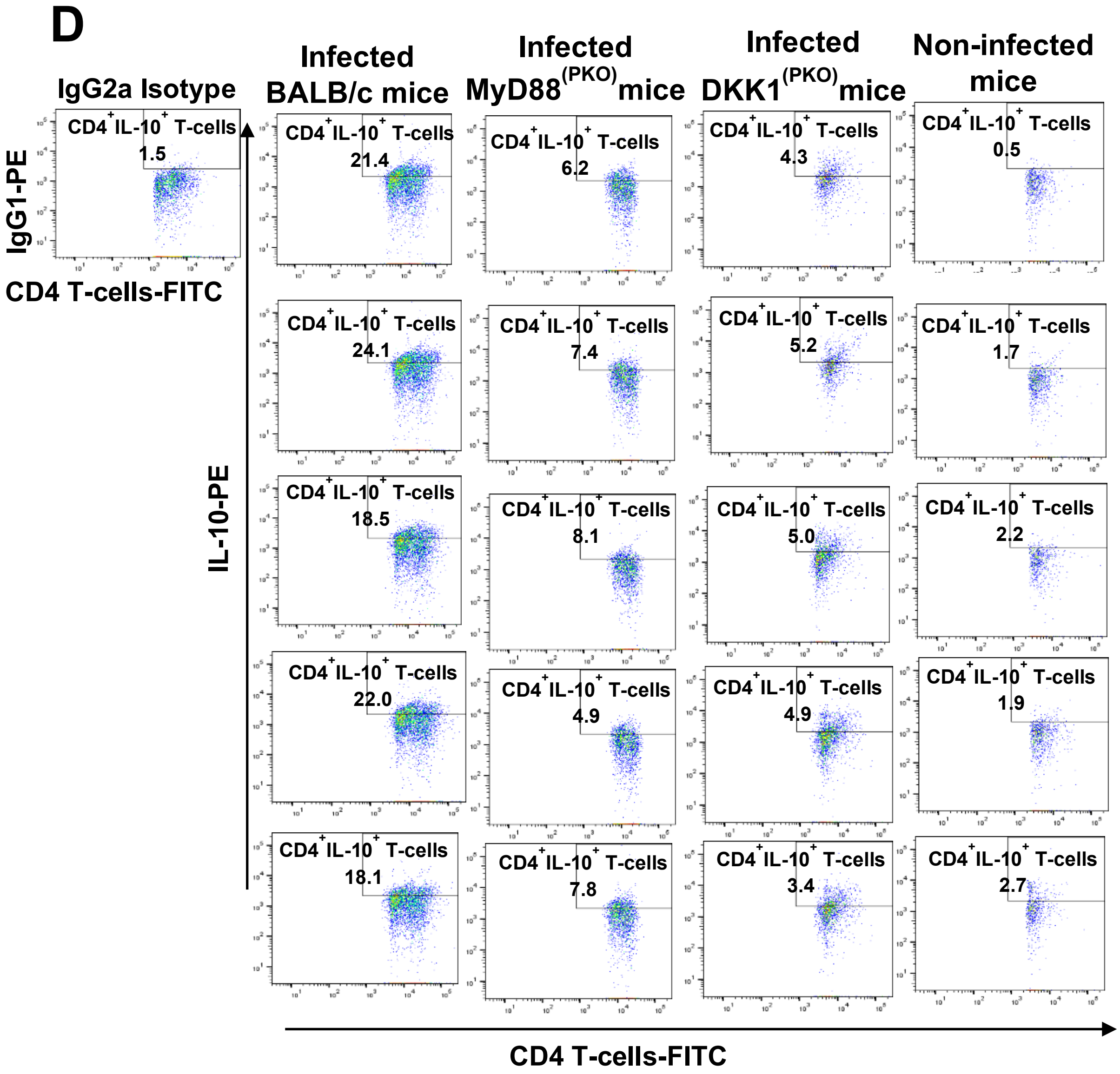
D
IgG2a Isotype
Infected BALB/c mice
Infected MyD88(PKO)mice
Infected DKK1(PKO)mice
Non-infected mice
IgG1-PE
IL-10-PE
CD4 T-cells-FITC
CD4+IL-10+ T-cells
1.5
21.4
6.2
4.3
0.5
24.1
7.4
5.2
1.7
18.5
8.1
5.0
2.2
22.0
4.9
4.9
1.9
18.1
7.8
3.4
2.7
CD4 T-cells-FITC

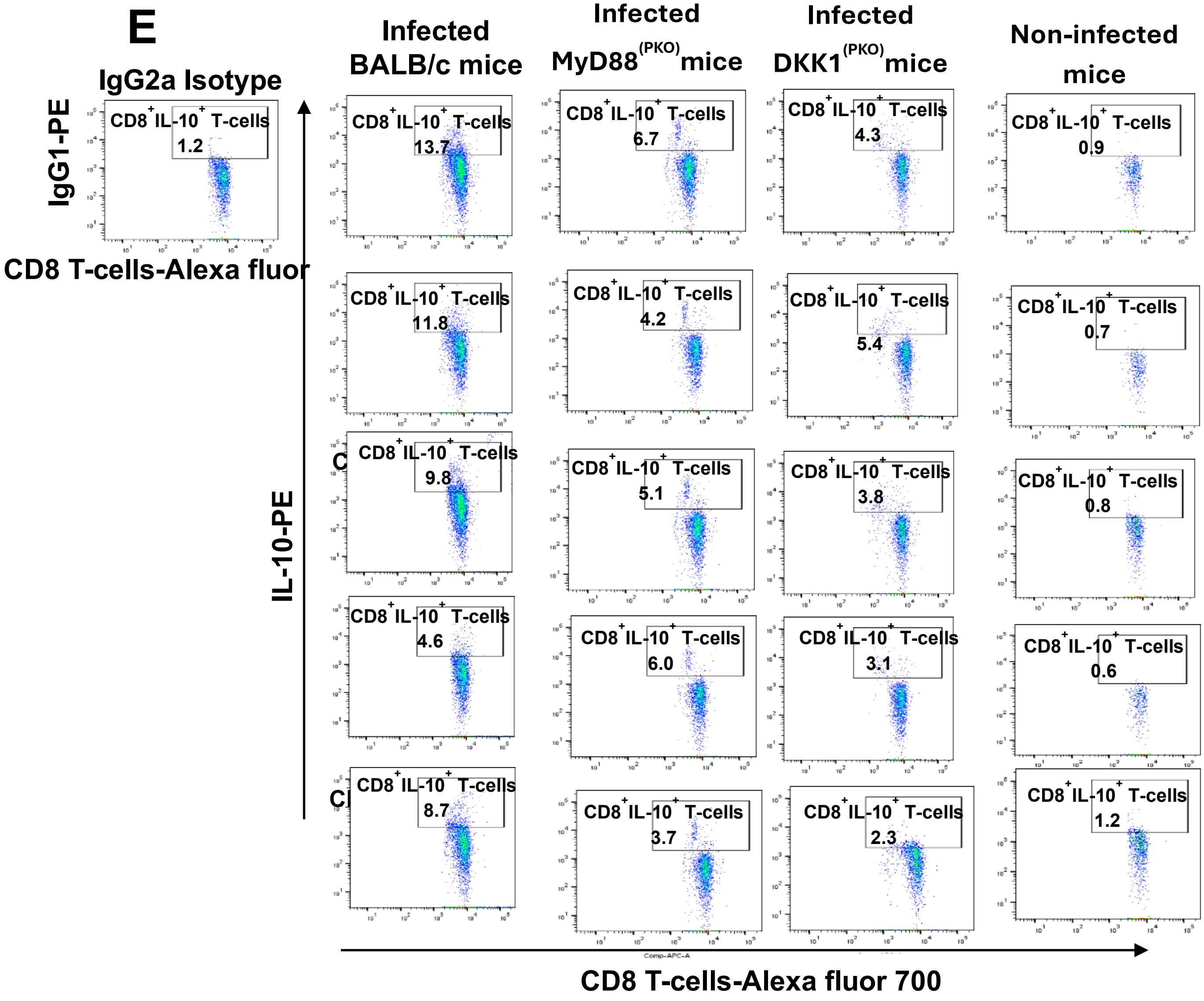
E
IgG2a Isotype
IgG1-PE
CD8+IL-10+ T-cells
1.2
CD8 T-cells-Alexa fluor
Infected BALB/c mice
Infected MyD88(PKO) mice
Infected DKK1(PKO) mice
Non-infected mice
IL-10-PE
CD8+IL-10+ T-cells 13.7
CD8+IL-10+ T-cells 6.7
CD8+IL-10+ T-cells 4.3
CD8+IL-10+ T-cells 0.9
CD8+IL-10+ T-cells 11.8
CD8+IL-10+ T-cells 4.2
CD8+IL-10+ T-cells 5.4
CD8+IL-10+ T-cells 0.7
CD8+IL-10+ T-cells 9.8
CD8+IL-10+ T-cells 5.1
CD8+IL-10+ T-cells 3.8
CD8+IL-10+ T-cells 0.8
CD8+IL-10+ T-cells 4.6
CD8+IL-10+ T-cells 6.0
CD8+IL-10+ T-cells 3.1
CD8+IL-10+ T-cells 0.6
CD8+IL-10+ T-cells 8.7
CD8+IL-10+ T-cells 3.7
CD8+IL-10+ T-cells 2.3
CD8+IL-10+ T-cells 1.2
CD8 T-cells-Alexa fluor 700

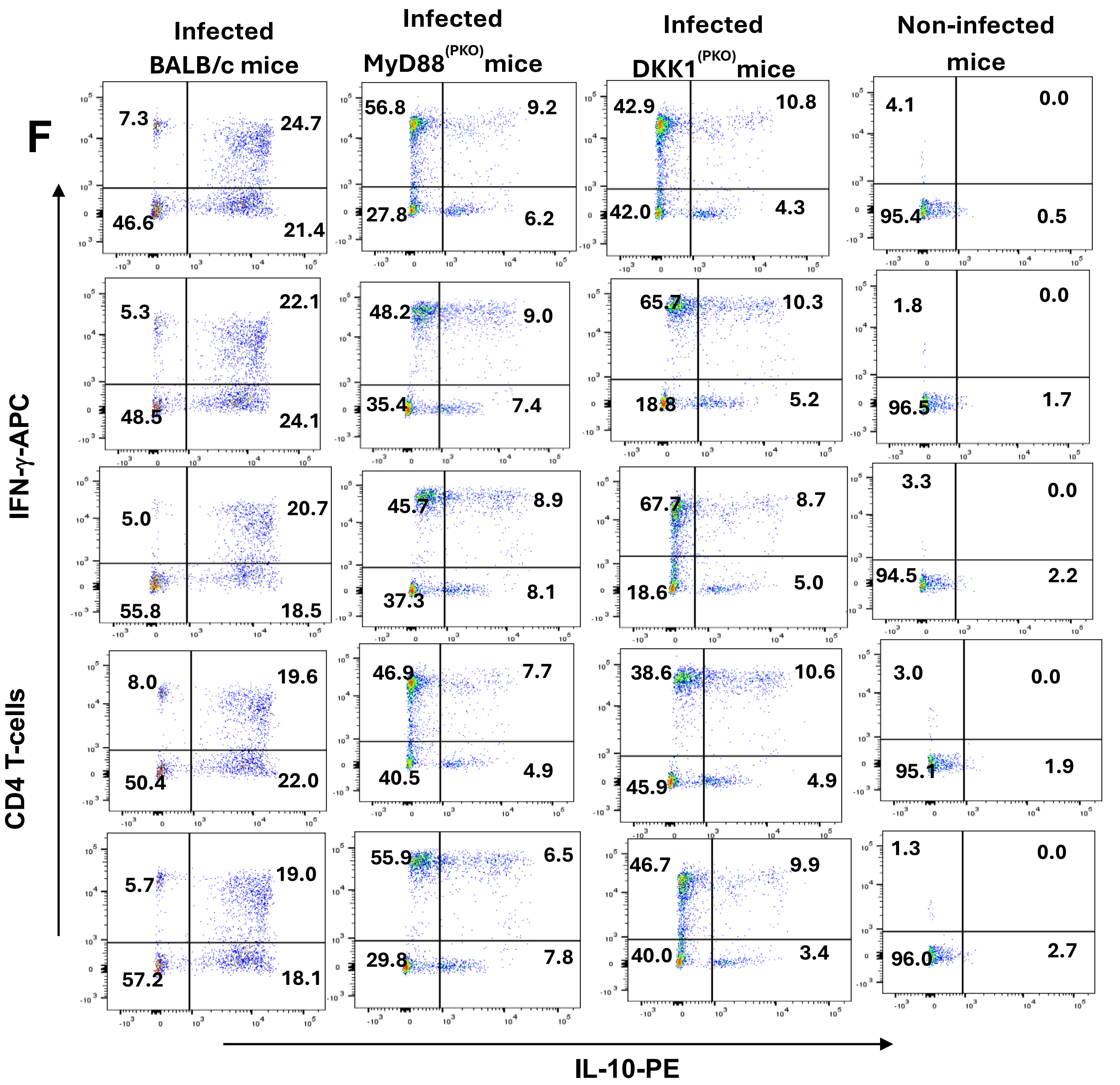


**Figure S4: Decreased $CD8^+IL\text{-}10^+$, $CD4^+IL\text{-}10^+$ and $CD4^+IL\text{-}10^+IFNg^+$ T-cells in $MyD88^{(PKO)}$ and $DKK1^{(PKO)}$ -infected mice on day 14 PI.** BALB/c, $MyD88^{(PKO)}$ and $DKK1^{(PKO)}$ mice were challenged with infective metacyclic promastigote (2 x $10^6$ parasites, n = 5) of *L. major* via the footpad. Non-infected BALB/c mice (n =5) were given 0.9% NaCl saline. Two weeks post-infection, the draining and non-draining lymph node cells were isolated. Lymph node cells were incubated with a cell stimulation cocktail for 5 hr. After a further 3 hr in BFA, cells were stained for intracellular IL-10 and IFN-γ. The stained lymph node cells from each mouse in BALB/c, $MyD88^{(PKO)}$ and $DKK1^{(PKO)}$ infected mice were determined for the percentage of $CD4^+$ and $CD8^+$ T-cells, percentage and MFI of $CD8^+IFN\text{-}\gamma^+$ or $CD8^+IL\text{-}10^+$ T-cells, percentage and MFI of $CD4^+IFN\text{-}\gamma^+$ or $CD4^+IL\text{-}10^+$ T-cells, and percentage of $CD4^+$ $IFN\text{-}\gamma^+$ $IL\text{-}10^+$ T-cells by flow cytometry. A dot plot of each sample in all the experimental groups is presented in **(A, B, C, D, E & F).**

A

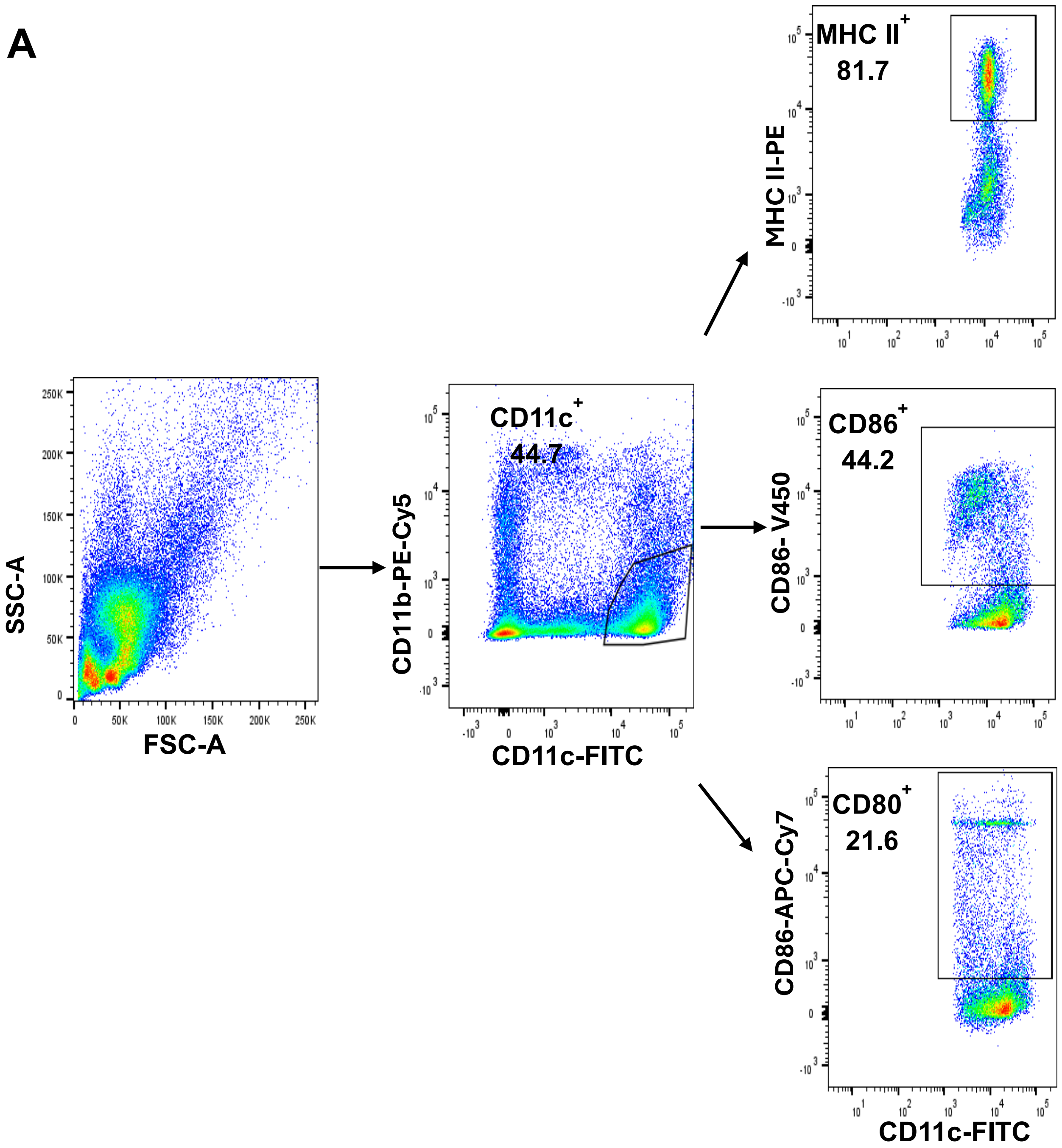
SSC-A
FSC-A
CD11b-PE-Cy5
CD11c-FITC
CD11c+
44.7
MHC II-PE
MHC II+
81.7
CD86- V450
CD86+
44.2
CD86-APC-Cy7
CD80+
21.6
CD11c-FITC

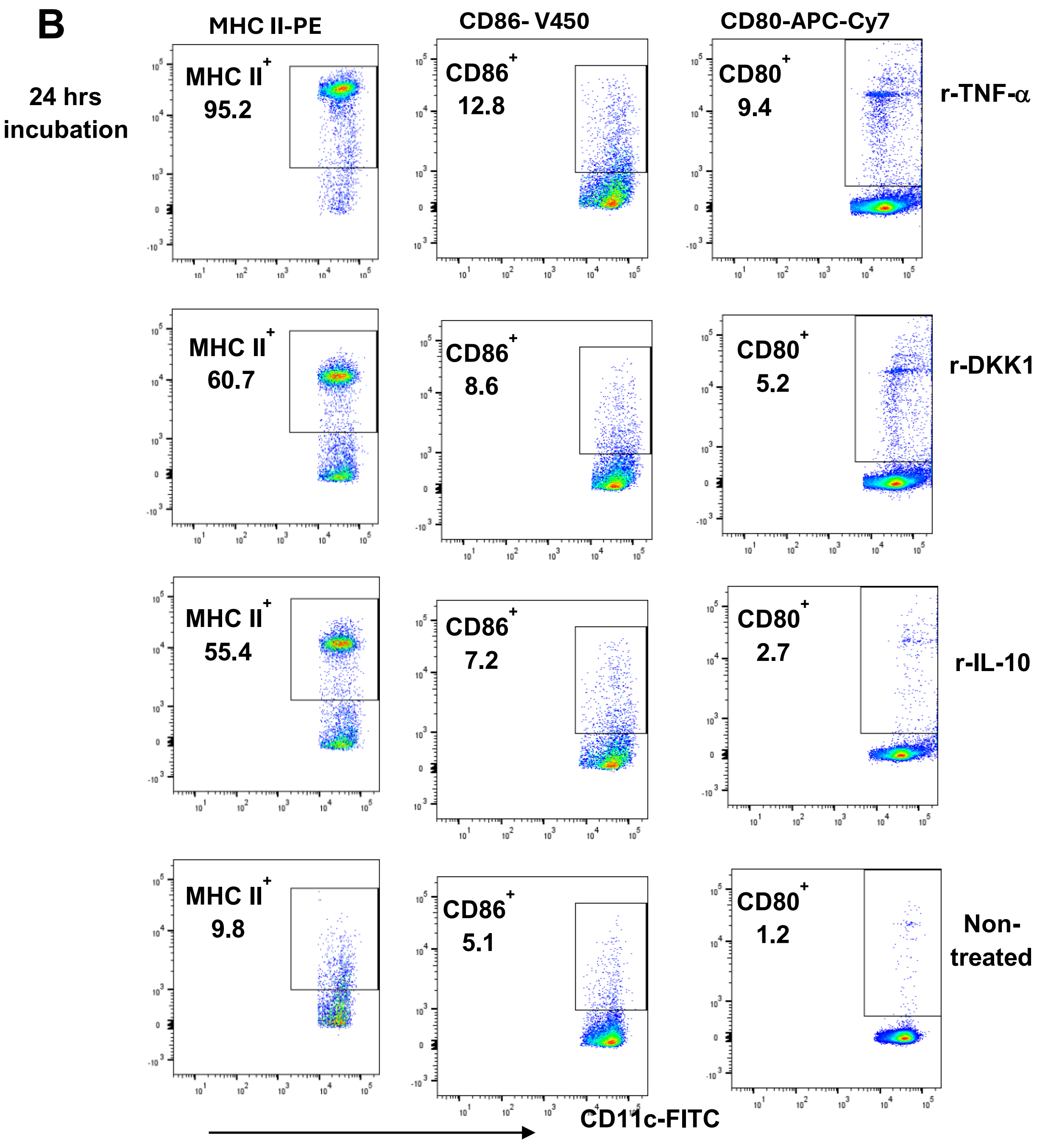
B
MHC II-PE
CD86- V450
CD80-APC-Cy7
24 hrs incubation
MHC II+ 95.2
CD86+ 12.8
CD80+ 9.4
r-TNF-α
MHC II+ 60.7
CD86+ 8.6
CD80+ 5.2
r-DKK1
MHC II+ 55.4
CD86+ 7.2
CD80+ 2.7
r-IL-10
MHC II+ 9.8
CD86+ 5.1
CD80+ 1.2
Non-treated
CD11c-FITC

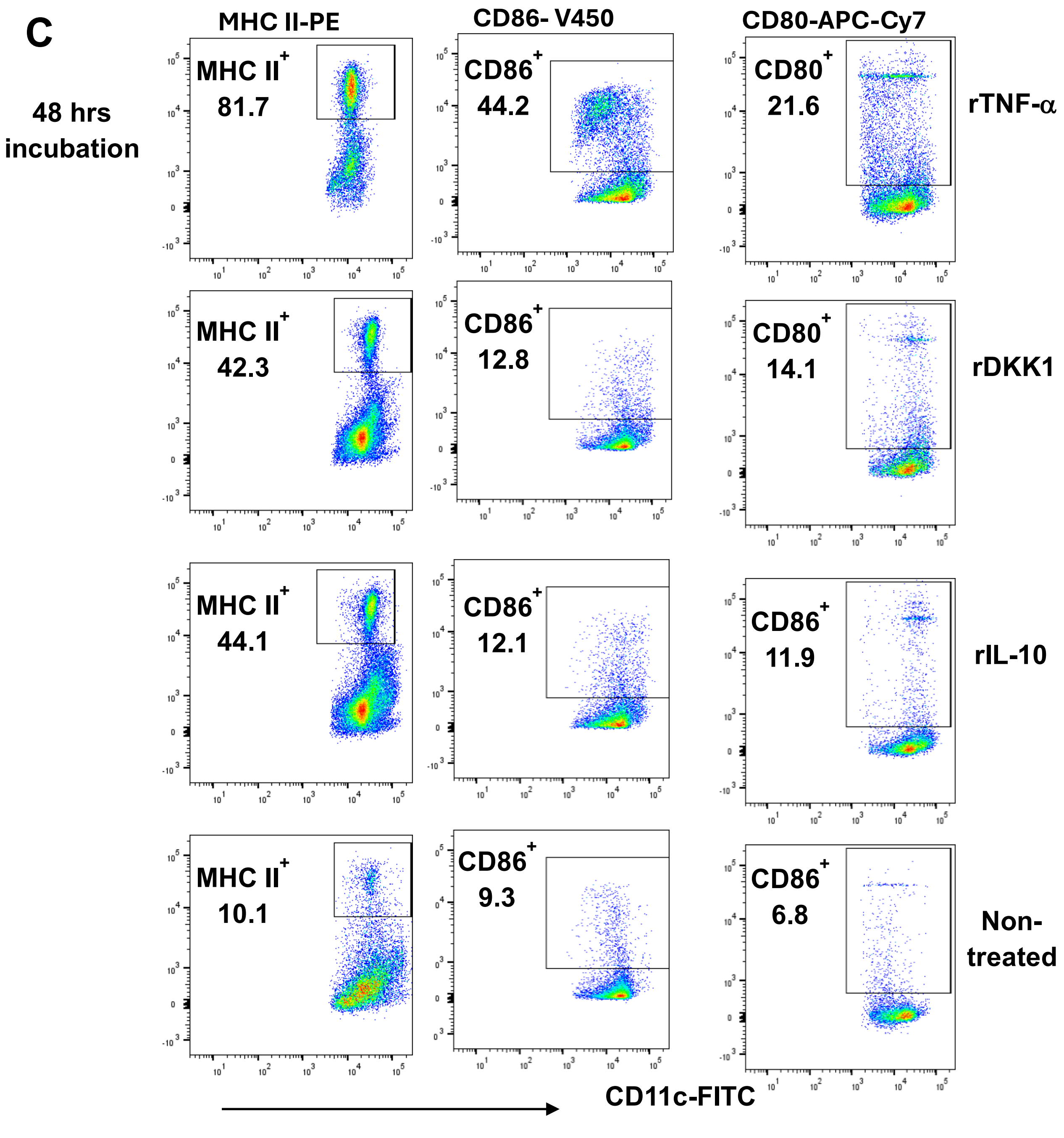


**Figure S5: rDKK1 inhibits the percentage of MHC II$^+$, CD86$^+$ and CD80$^+$ dendritic cells.** Monocyte-derived dendritic cells were incubated in rDKK1(100 ng/ml), rIL-10 (20 ng/ml) and rTNF-α (10 ng/ml). Cells harvested at 24 and 48 hrs post-incubation were used to determine the percentage of MHC II$^+$, CD86$^+$ and CD80$^+$ dendritic cells by flow cytometry. Representative flow cytometry dot plots generated 24 hrs post-incubation showed the analysis of MHC II$^+$, CD86$^+$ and CD80$^+$ is indicated (**A**). A dot plot of each sample in all experimental conditions is presented in (**B**) & (**C**).
